# Genome editing excisase origins illuminated by somatic genome of *Blepharisma*

**DOI:** 10.1101/2021.12.14.471607

**Authors:** Minakshi Singh, Kwee Boon Brandon Seah, Christiane Emmerich, Aditi Singh, Christian Woehle, Bruno Huettel, Adam Byerly, Naomi Alexandra Stover, Mayumi Sugiura, Terue Harumoto, Estienne Carl Swart

## Abstract

Massive DNA excision occurs regularly in ciliates, ubiquitous microbial eukaryotes with somatic and germline nuclei in the same cell. Tens of thousands of internally eliminated sequences (IESs) scattered throughout a copy of the ciliate germline genome are deleted during development of the streamlined somatic genome. *Blepharisma* represents one of the two earliest diverging ciliate classes, and, unusually, has dual pathways of somatic nuclear development, making it ideal for investigating the functioning and evolution of these processes. Here, we report the somatic genome assembly of *Blepharisma stoltei* strain ATCC 30299 (41 Mb), arranged as numerous alternative telomere-capped minichromosomes. This genome encodes eight PiggyBac transposase homologs liberated from transposons. All are subject to purifying selection, but just one, the putative IES excisase, has a complete catalytic triad. We propose PiggyBac homologs were ancestral excisases that enabled evolution of extensive, natural genome editing.

## Introduction

DNA excision in ciliates is a spectacular and widespread form of natural genome editing with profound consequences for what germline and somatic genomes mean (Arnaiz et al., 2012; Chen et al., 2014; Hamilton et al., 2016; Swart and Nowacki, 2015). Though the responsible processes are under active study, much remains to be learnt from these master DNA manipulators, including how and why this remarkable situation arose in them.

Knowledge of ciliate genome editing mechanisms is dominated by *Tetrahymena* and *Paramecium* (class Oligohymenophorea), with additional input from *Oxytricha*, *Stylonychia* and *Euplotes* (class Spirotrichea) (Chalker et al., 2013; Vogt et al., 2013). The remaining nine ciliate classes await detailed characterization. To advance investigation of natural genome editing and tackle questions about its origin we focused on the ciliate species *Blepharisma stoltei*. Together with its sister-class, Karyorelictea, the class Heterotrichea, to which this ciliate species belongs, represent the earliest branching ciliate lineages, more distantly related to current model ciliates than those models are to each other (Lynn, 2010). Furthermore, the genus *Blepharisma* exhibits distinctive alternative somatic nuclear developmental pathways, which have the potential to disentangle genome editing processes from indirect influences of preceding pathways.

*Blepharisma* is a distinctive genus of single-celled ciliates known for the red, light-sensitive pigment, blepharismin, in their sub-pellicular membranes (Giese, 1973), and unusual nuclear/developmental biology (Figure 1) (Miyake et al., 1991). To date molecular investigations and genomics of ciliates have predominantly focused on oligohymenophoreans and spirotrichs (Figure 2, Table S1). In recent years, publication of a draft genome for the heterotrich ciliate, *Stentor,* has facilitated revival of this genus for investigations of cellular regeneration (Slabodnick and Marshall, 2014; Slabodnick et al., 2017; Zhang et al., 2021). However, significant hurdles still need to be overcome to investigate genome editing in *Stentor coeruleus* since requisite cell mating has not been observed in the reference somatic genome strain (personal communication, Mark Slabodnick), and very high lethality has been reported for other strains in which mating occurred (Rapport et al., 1976). We therefore focused on *Blepharisma* which is amenable to such investigations, with controlled induction of mating, and, critically, established procedures for investigating cellular and nuclear development from more than a century of meticulous cytology (Friedl et al., 1983; Giese, 1973; Harumoto et al., 1998; Inaba, 1965; Kobayashi et al., 2015; Kumazawa, 1979; Miyake and Harumoto, 1990; Miyake et al., 1979; Repak, 1968; Salvini et al., 1983; Sugiura et al., 2010; Terazima and Harumoto, 2004; Young, 1937).

**Figure 1.**
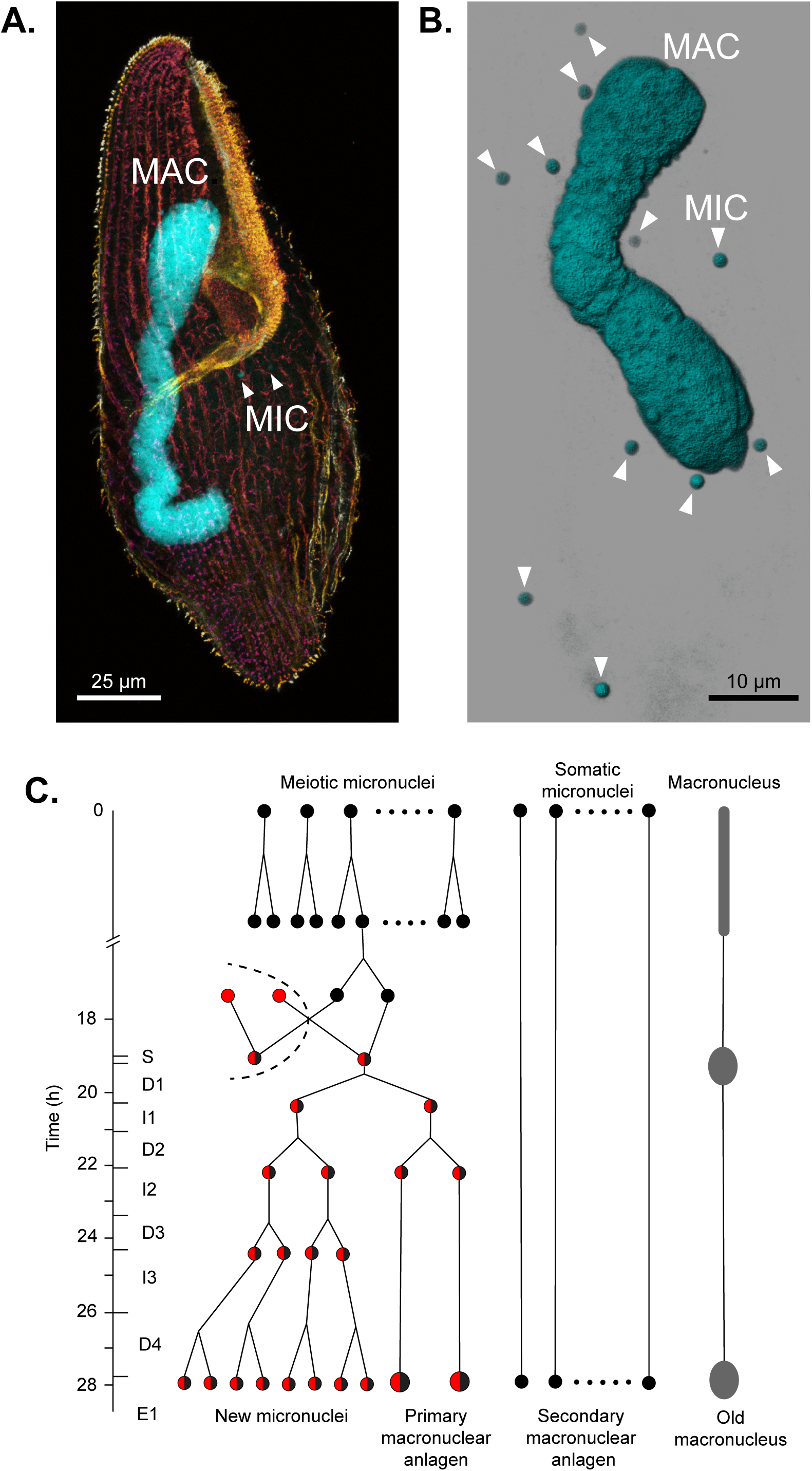
*Blepharisma* nuclei and nuclear development during conjugation. **A.** *B. stoltei* ATCC 30299 cell stained with anti-alpha tubulin-Alexa488 (depth-color coded red to yellow) and DAPI (cyan). **B.** Snapshot of a 3D reconstruction (Imaris, Bitplane) from CLSM images of Hoechst 33342 (dsDNA dye, Invitrogen™) fluorescence (Ex405 nm / Em420-470 nm). **C.** Schematic of nuclear processes occurring during conjugation (classified according to, and modified from (Miyake et al., 1991)). Nuclear events occurring before and up to, but not including fusion of the gametic nuclei (syngamy) are classified into sixteen pre-gamic stages where the MICs undergo meiosis and the haploid products of meiotic MICs are exchanged between the conjugating cells, followed by karyogamy. After karyogamy, cells are classified into 10 stages S (synkaryon), D1 (1^st^ mitosis), I1 (1^st^ interphase), D2 (2^nd^ mitosis), I2 (2^nd^ interphase), D3 (3^rd^ mitosis), I3 (3^rd^ interphase), D4 (4^th^ mitosis), E1 (1^st^ embryonic stage), E2 (2^nd^ embryonic stage). After E2, the exconjugants divide further and are classified into 6 stages of cell division (CD1-6) which we did not follow here. See also Figure 4.

**Figure 2.**
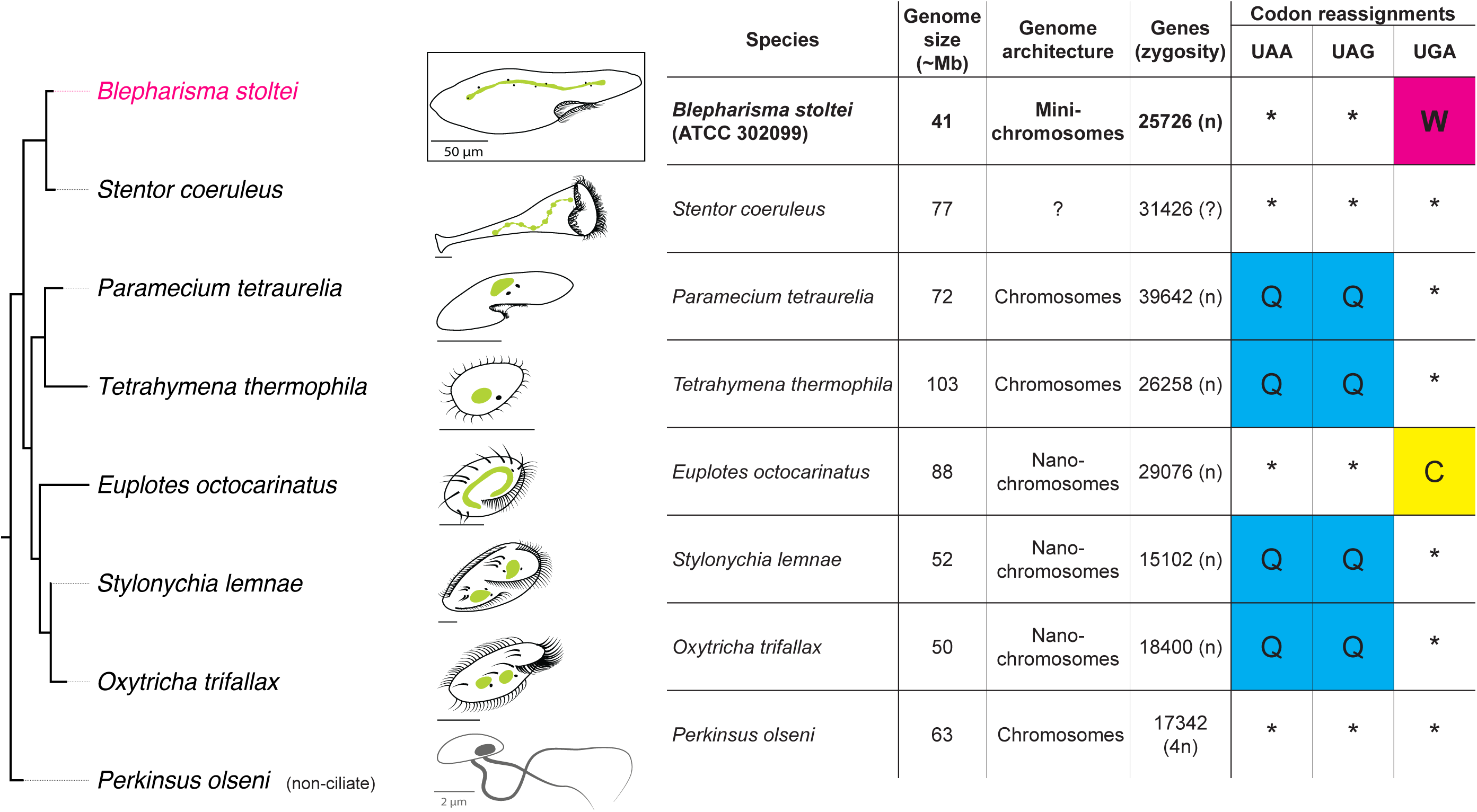
Basic properties of ciliate MAC genomes. In cell diagrams MACs are green and MICs are small black dots in close proximity to MACs. Citations for genome properties are in Data S1. See also Figure S1.

The *Blepharisma stoltei* strains used in the present study were originally isolated in Germany (strain ATCC 30299) and Japan (strain HT-IV), with the former continuously cultured for over fifty years, and the latter for over a decade. The cells are comparatively straightforward to maintain, e.g., stable cultures can be established in a simple salt medium on a few grains of rice. Due to their distinctive pigmentation and large size several *Blepharisma* species are excellent subjects for introducing cell biology concepts to non-specialists, and are thus readily available for educational purposes from commercial suppliers. They are ideal subjects for behavioral and developmental investigations, e.g., as voracious predators of smaller ciliates and other unicellular species, and also exhibit pronounced phenotypic plasticity, including forming cysts and giant, cannibal cells under suitable conditions (Giese, 1973).

Like all ciliates (Prescott, 1994), *Blepharisma* cells have two types of nuclei: a macronucleus (MAC) which is very large and transcriptionally active during vegetative growth, and a small, generally transcriptionally inactive micronucleus (MIC), which serves as the germline (Figure 1A, B). In vegetative propagation (asexual replication) of *Blepharisma*, cell fission results in half of the MAC pinching off before distributing to each of the resulting daughter cells together with the mitotically divided MICs. Upon starvation, *Blepharisma* cells, like other ciliates, are also capable of sexual processes initiated by conjugation. Essential for developmental investigations, the intricate ballet of nuclear movements and morphological changes occurring during *Blepharisma* conjugation is well-documented (Miyake et al., 1991) (Figure 1C). During this process half of the MICs in each of the cells undergo meiosis (meiotic MICs) and the rest do not (somatic MICs) (Figure 1C). One of the meiotic MICs eventually gives rise to two haploid gametic nuclei. One gametic MIC (the migratory nucleus) from each conjugating cell is exchanged with that of its partner. In parallel in partnered cells, subsequent fusion of the migratory and stationary haploid nuclei generates a zygotic nucleus (synkaryon), and after successive mitotic divisions gives rise to both new MICs and new MACs (known as anlagen). The new MACs continue to mature, eventually growing in size and DNA content (Miyake et al., 1991).

Conveniently for investigations of development and genome editing, *Blepharisma* is one of only two ciliate genera, along with *Euplotes* (Katashima, 1959; Kimball, 1942; Luporini et al., 1983; Vallesi et al., 1995), where conjugation has been shown to be mediated through pheromone-like substances called gamones. *Blepharisma* has two mating types, distinguished by their gamone production. Mating type I cells release gamone 1, a ∼30 kDa glycoprotein (Miyake and Beyer, 1974; Sugiura and Harumoto, 2001); mating type II cells release gamone 2, calcium-3-(2’- formylamino-5’-hydroxybenzoyl) lactate, a small-molecule effector (Kubota et al., 1973). *Blepharisma* cells commit to conjugation when complementary mating types recognize each other’s gamones, with the cells remaining paired while meiosis and fertilization occur and eventually new MACs begin to form.

As in model ciliates, we show in an accompanying paper that MIC-specific sequences are removed to form a functional *Blepharisma* MAC genome (Seah, et al. 2022). Like other ciliates the resulting MAC genome appears to have been freed of mobile elements and other forms of junk DNA contained in the MIC genome (Klobutcher and Herrick, 1997). However, this situation is an oversimplification of the actual MAC genome content (Seah, et al. 2022). In the best studied ciliates, genome editing is thought to be coordinated or assisted by small RNAs (sRNAs) (Chalker et al., 2013). Specific MIC-limited DNA segments — internally eliminated sequences (IESs) — are excised by domesticated transposases (Arnaiz et al., 2012; Chalker et al., 2013; Klobutcher and Herrick, 1995; Prescott, 1994). Large scale genome-wide DNA amplification accompanies genome editing, producing thousands of copies in mature MACs of larger ciliate species (Klobutcher and Herrick, 1997; Prescott, 1994).

We were motivated to investigate genome editing in *Blepharisma*, as, unlike model ciliates, these cells can produce two kinds of anlagen, and because one of their two developmental pathways skips the preceding series of mitoses, meioses, nuclear exchanges and fertilization (Miyake et al., 1991) (Figure 1C). Primary anlagen mature in the conventional manner from zygotic nuclei. Somatic MICs which have not undergone meiosis can give rise to secondary anlagen, which can develop into mature macronuclei (Miyake et al., 1991). This occurs frequently in strains with a high selfing frequency (conjugation among cells within a clonal population), in preference to development of primary MAC anlagen (Miyake et al., 1991). This alternative pathway of MAC development has also been observed experimentally after removal of primary MAC anlagen by microsurgery (Miyake et al., 1991). As conjugation progresses, the old (maternal) MACs are progressively degraded (Miyake et al., 1991). Since the *B*. *stoltei* MIC genome has numerous gene-interrupting IESs (Seah et al. 2022), in principle, editing of DNA needs to occur in both primary and secondary anlagen to produce functional MAC genomes.

Here we provide essential somatic genome and transcriptomic resources for *B. stoltei.* From long-read sequencing, the *B. stoltei* MAC genome appears to be organized as numerous alternative minichromosomes. Among *Blepharisma*’s MAC-encoded transposase genes we identified were PiggyBac transposase homologs, which, thus far only reported in the distantly related ciliates *Paramecium* and *Tetrahymena*. A few *Blepharisma* PiggyBac homologs are substantially upregulated in MAC development, including the main candidate IES excisase. Consistent with ancient origins of ciliate genome editing, *Blepharisma* shares pronounced development-specific upregulation of homologs known to be involved in this process. *Blepharisma* therefore represents an invaluable outgroup for investigations of genome editing evolution.

## Results

### A compact somatic genome with a minichromosomal architecture

The draft *Blepharisma stoltei* ATCC 30299 MAC genome is compact (41 Mb) and AT rich (66%), like most sequenced ciliate MAC genomes (Figure 2; Table S1, 2, Figure S1A). The genome is gene-dense (25,711 predicted genes), with short intergenic regions, tiny, predominantly 15 and 16 bp introns (Figure S4; Supplemental information, “Tiny spliceosomal introns”) and untranslated regions (UTRs) (Figure 3A). *B. stoltei* uses an alternative nuclear genetic code with UGA codons reassigned from stops to tryptophan (Figure S1B).

**Figure 3.**
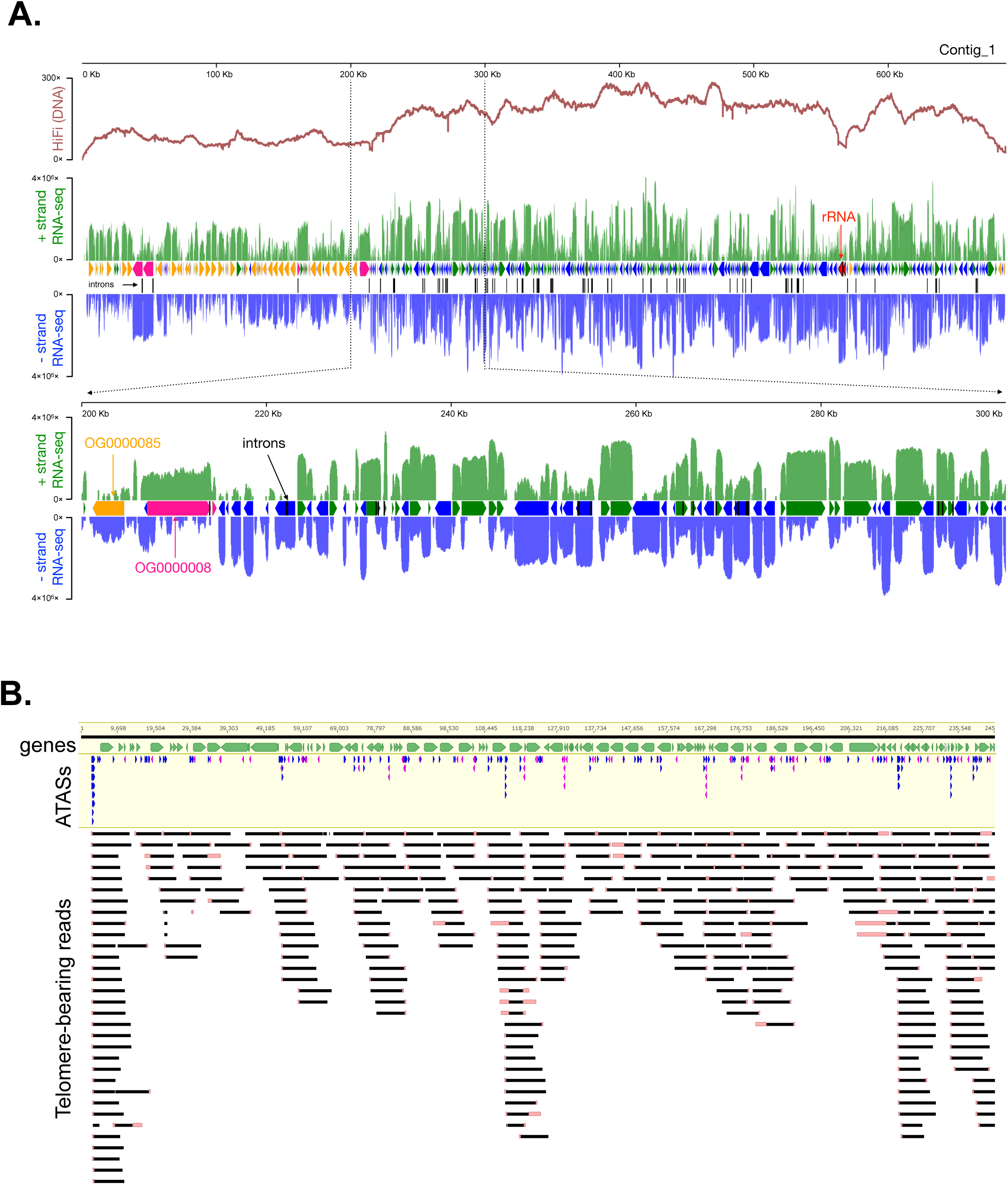
A gene-dense somatic genome with a minichromosomal architecture. **A.** HiFi (DNA) and RNA-seq coverage across a representative *B. stoltei* ATCC30299 MAC genome contig (Contig_1). Y scale is linear for HiFi reads and logarithmic (base 10) for RNA-seq. Plus strand (relative to the contig) RNA-seq coverage is green; minus strand RNA-seq coverage is blue. Between the RNA-seq coverage graphs each arrow represents a predicted gene. Two orthogroups classified by OrthoFinder are shown. **B.** Mapping of a subset telomere-containing HiFi reads to a *B. stoltei* MAC genome contig region, with alternative telomere addition sites (ATASs) shown by blue (5’) or mauve (3’) arrows. Pink bars at read ends indicate soft-masking, typically of telomeric repeats. See also Figure S2-5.

From joint variant calling of reads from strains ATCC 30299 and HT-IV, strain ATCC 30299 appears to be virtually homozygous, with only 1277 heterozygous single-nucleotide polymorphisms (SNPs) compared to 193725 in strain HT-IV (i.e., individual heterozygosity of 3.08 × 10^-5^ vs. 4.67 × 10^-3^ respectively). Low SNP levels were likely beneficial for overall genomic contiguity, since heterozygosity poses significant algorithmic challenges for assembly software (Chin et al., 2016). For brevity’s sake, we refer to this genome as the *Blepharisma* MAC genome (and “*Blepharisma*” for the associated strain). Though the final assembly comprises 64 telomere-to-telomere sequences, chromosomes and their ends are meaningless given the extensive natural fragmentation of the *Blepharisma* MAC genome (characterized in the next section), hence we simply refer to “contigs”.

The basic telomere unit of *Blepharisma* is a permutation of CCCTAACA, like its heterotrich relative *Stentor coeruleus* (Slabodnick et al., 2017) (Figure S2). Since a compelling candidate for a telomerase ncRNA (TERC) could not be found in either *Blepharisma* or *Stentor* using Infernal (Nawrocki et al., 2009) and RFAM models (RF00025 - ciliate TERC; RF00024 - vertebrate TERC), it was not possible to delimit the repeat ends. Heterotrichs may use a different or very divergent ncRNA. In contrast to the extremely short (20 bp) MAC telomeres of spirotrichs like *Oxytricha* with extreme MAC genome fragmentation (Swart et al., 2013), sequenced *Blepharisma* MAC telomeres are moderately long (Figure S2A), with a mode of 209 bp (∼26 repeats of the 8 bp motif), extending to a few kilobases.

With a moderately strict definition of possessing at least three consecutive telomeric repeats, one in eight reads in the *Blepharisma* HiFi library were telomere-bearing. Telomeric reads are distributed across the entire genome (Figure 3B). Typically, a minority of mapped reads are telomere-bearing at individual internal positions, and so we term them alternative telomere addition sites (ATASs) (Figure 3B). We identified 46705 potential ATASs, the majority of which (38686) were represented by only one mapped HiFi read.

The expected distance between telomeres, and hence the average MAC DNA molecule length, is about 130 kb. This is consistent with the raw input MAC DNA lengths, which were mostly longer than 10 kb and as long as 1.5 Mb (Figure S3A, B), and the small fraction (1.3%) of *Blepharisma*’s HiFi reads bound by telomeres on both ends. Excluding the length of the telomeres, telomere-bound reads may be as short as 4 kb (Figure S2B). Given the frequency of telomere-bearing reads, we expect many additional two-telomere DNA molecules longer than 12 kb, the approximate maximum length of the HiFi reads (Figure S3A, B).

Since the lengths of the sequenced two-telomere DNA molecules on average imply that they encode multiple genes, we propose classifying them as “minichromosomes”. This places them between the “nanochromosomes” of ciliates like *Oxytricha* and *Stylonychia*, which typically encode single genes and a few kilobases long (Aeschlimann et al., 2014; Swart et al., 2013), and *Paramecium tetraurelia* and *Tetrahymena thermophila* MAC chromosomes which are hundreds of kilobases to megabases long (Aury et al., 2006; Sheng et al., 2020; Zagulski et al., 2004). The *Paramecium bursaria* MAC genome is considerably more fragmented than those of other previously examined *Paramecium* species, and have thus also been classified as minichromosomes (Cheng et al., 2020).

### Key features of gene expression during new MAC development

To gain an overview of the molecular processes during *Blepharisma* genome editing, we examined gene expression trends across development. Complementary *B. stoltei* strains were treated with gamones of the opposite mating type, before mixing to initiate conjugation (Miyake et al., 1991; Sugiura et al., 2012). Samples for morphological staging and RNA-seq were taken at intervals from the time of mixing (“0 hour” time point) up to 38 hours.

During *Blepharisma* conjugation, meiosis begins around 2 h after conjugating cell pairs form and continues up to 18 h, by when gametic nuclei generated by meiosis have been exchanged (Figure 1C; Figure 4). This is followed by karyogamy and mitotic multiplication of the zygotic nucleus (22 hours). At 26 h, new, developing primary MACs can be observed in the conjugating pairs, as large, irregular bodies (Figure 4). These nuclei mature into the new MACs of the exconjugant cell by 38 h, after which cell division generates two daughter cells. Smaller secondary MACs, derived directly from MICs without all the intermediate nuclear stages, can also be seen from 22 h, eventually disappearing, giving way to the primary MACs (Figure 4).

**Figure 4.**
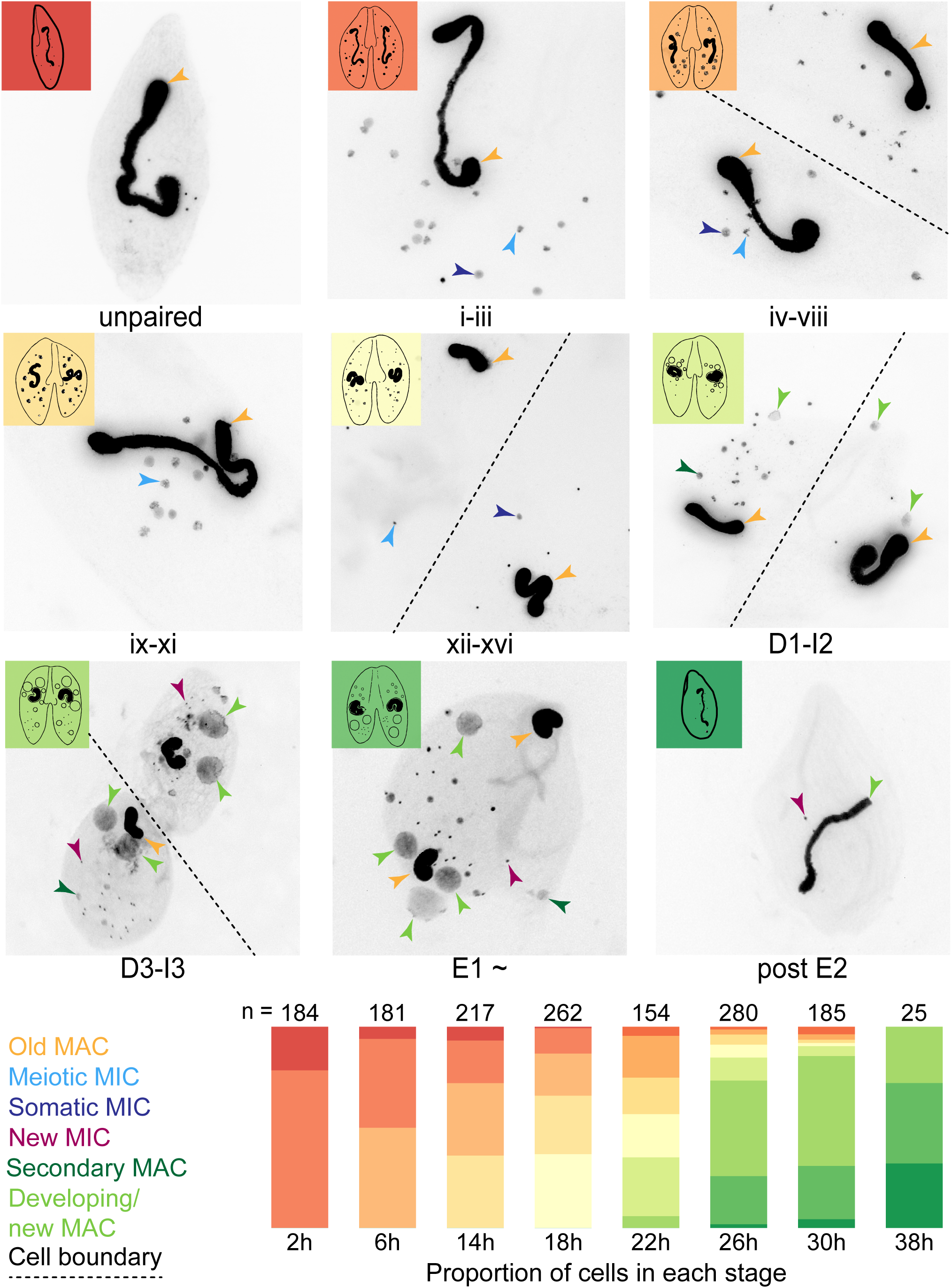
Developmental staging of *B. stoltei* for RNA-seq. Classification of nuclear morphology into stages is according to previous descriptions (Miyake et al., 1991). Nuclear events occurring before and up to, but not including fusion of the gametic nuclei (syngamy) are classified into sixteen stages indicated by roman numerals. These are the pre-gamic stages of conjugation where the MICs undergo meiosis and the haploid products of meiotic MICs are exchanged between the conjugating cells. Stages after syngamy are classified into 10 stages as in Figure 1. Illustration of various cell stages adapted from (Suzuki, 1957)). Stacked bars show the proportion of cells at each time point at different stages of development, preceded by the number of cells inspected (n). See also Figure S6.

Examining gene expression at 26 h, when the majority of cells are forming a new MAC (Figure 4), we observe two broad trends: relatively stable constitutive gene expression (Table S5; Data S3), e.g., an actin homolog (ENA accession: BSTOLATCC_MAC19444) and a bacteria-like globin protein (BSTOLATCC_MAC21846), versus pronounced development-specific upregulation (Table S6; Data S3), e.g., a histone (BSTOLATCC_MAC21995) an HMG box protein (BSTOLATCC_MAC14030), and a translation initiation factor (eIF4E, BSTOLATCC_MAC5291). We eschewed a shallow Gene Ontology (GO) enrichment analysis, instead favoring close scrutiny of a smaller subset of genes strongly upregulated during new MAC formation. For this, computational gene annotations in combination with BLASTP searches and examination of literature associated with homologs was used. Ranking the relative gene expression at 26 h vs. the average expression of starved, gamone treated, and 0 h cells, in descending order, revealed numerous genes of interest, including homologs of proteins known to be involved in genome editing in model ciliates (Table S6).

Among the top 100 genes ranked this way (69× to 825× upregulation) nine contain transposase domains from PFAM: DDE_Tnp_1_7, DDE_3, MULE and DDE_Tnp_IS1595 (e.g., BSTOLATCC_MAC2188, BSTOLATCC_MAC14490, BSTOLATCC_MAC18054, BSTOLATCC_MAC18052, respectively). We also observe small RNA (sRNA) biogenesis and transport proteins, i.e., a Piwi protein (BSTOLATCC_MAC5406) and a Dicer-like protein (BSTOLATCC_MAC1138; “Supplemental information”, “Homologs of small RNA-related proteins involved in ciliate genome editing” and Figure S8), and a POT1 telomere-binding protein homolog (POT1.4; BSTOLATCC_MAC1496; Supplemental information “Telomere-binding protein paralogs”). Numerous homologs of genes involved in DNA repair and chromatin are also present among these highly developmentally upregulated genes (“Supplemental information”, “Development-specific upregulation of proteins associated with DNA repair and chromatin” and “Development-specific histone variant upregulation”). The presence of proteins involved in either transcription initiation or translation initiation among these highly upregulated genes suggests a possible manner in which regulation of development-specific gene expression may be coordinated (“Supplemental information”, “Development-specific upregulation of proteins associated with initiation of transcription and translation”).

### A single *Blepharisma* PiggyBac homolog has a complete catalytic triad

In *Paramecium tetraurelia* and *Tetrahymena thermophila*, PiggyBac transposases are responsible for IES excision during genome editing (Baudry et al., 2009; Cheng et al., 2010). These transposases appear to have been domesticated, i.e., their genes are no longer contained in transposons but are encoded in the somatic genome where they play an essential genome development role (Baudry et al., 2009; Cheng et al., 2010). PiggyBac homologs typically have a DDD catalytic triad, rather than the more common DDE triad of other DDE/D transposases (Yuan and Wessler, 2011). The DDD catalytic motif is present in *Paramecium* PiggyMac (Pgm) and *Tetrahymena* PiggyBac homologs Tpb1 and Tpb2 (Bischerour et al., 2018; Cheng et al., 2010). Among ciliates, domesticated PiggyBac transposases have so far only been reported in these model oligohymenophorean genera. Notably they have not been detected in either the MAC or MIC genome of the spirotrich *Oxytricha trifallax* (Chen et al., 2014; Swart et al., 2013).

We detected more transposase domains (9 distinct PFAM identifiers) in *Blepharisma* than any other ciliate species we examined (Figure 5A). Using HMMER searches with the domain characteristic of PiggyBac homologs, DDE_Tnp_1_7 (PF13843), we found eight homologs in *B. stoltei* ATCC MAC genome and five additional ones within IESs, none of which were flanked by terminal repeats (identified by RepeatModeler). We also found PiggyBac homologs in the MAC genomes of *B. stoltei* HT-IV and *B. japonicum* R1072.

**Figure 5.**
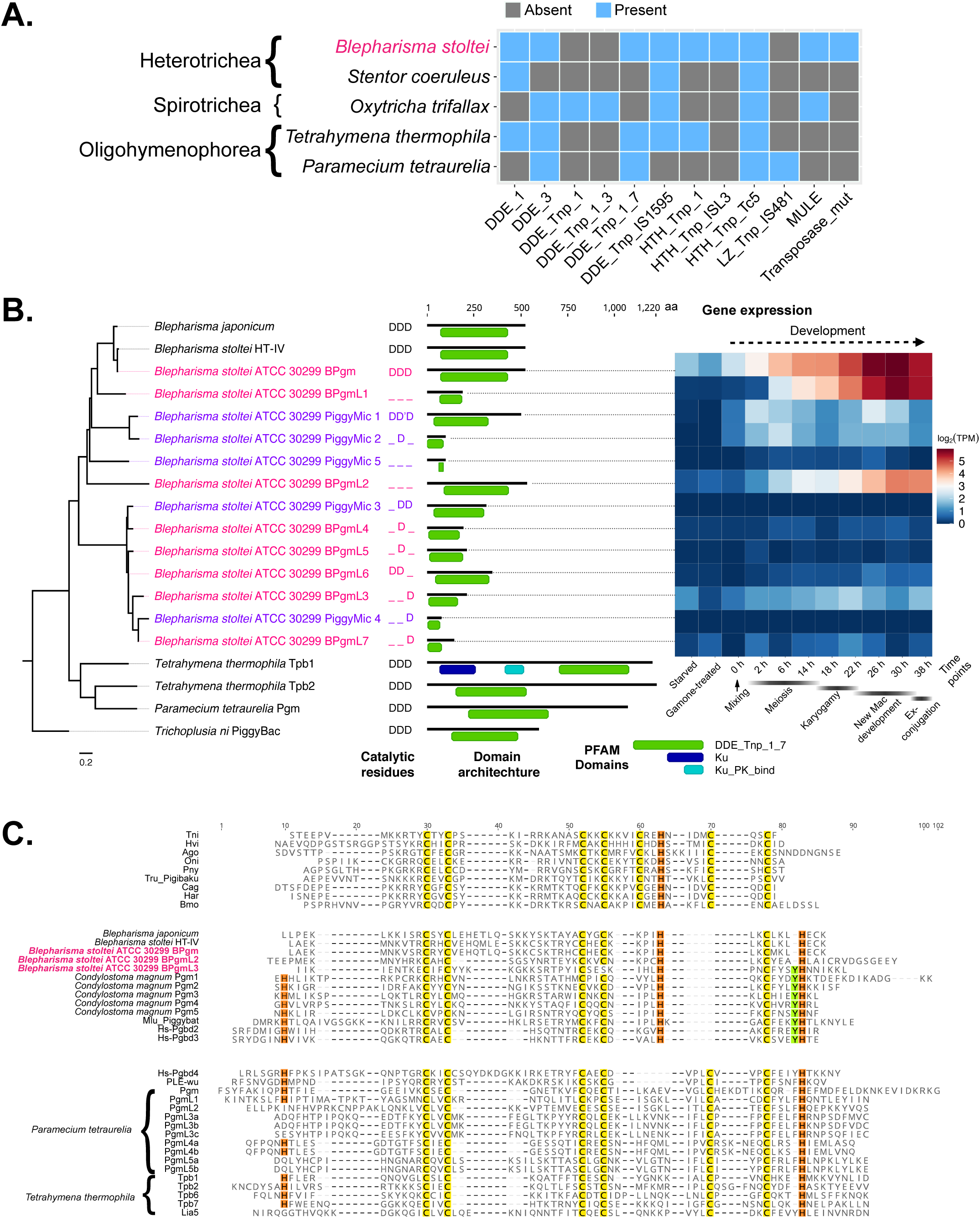
MAC genome-encoded transposases in ciliates and properties of a putative *Blepharisma* IES excisase. **A.** Presence/absence matrix of PFAM transposase domains detected in predicted MAC genome-encoded ciliate proteins. Ciliate classes are indicated before the binomial species names. **B**. DDE_Tnp_1_7 domain phylogeny with PFAM domain architecture and gene expression heatmap for *Blepharisma*. “Mixing” indicates when cells of the two complementary mating types were mixed. Outgroup: PiggyBac element from *Trichoplusia ni*. Catalytic residues: D-aspartate, D’-aspartate residue with 1 aa translocation. **C.** Cysteine-rich domains of PiggyBac homologs. PBLE transposases: Ago (*Aphis gossypii*); Bmo (*Bombyx mori*); Cag (*Ctenoplusia agnata*); Har (*Helicoverpa armigera*); Hvi (*Heliothis virescens*); PB-Tni (*Trichoplusia ni*); Mlu (PiggyBat from *Myotis lucifugus*); PLE-wu (*Spodoptera frugiperda*). Domesticated PGBD transposases: Oni (*Oreochromis niloticus*); Pny (*Pundamilia nyererei*); Lia5, Tpb1, Tpb2, Tpb6 and Tpb7 (*Tetrahymena thermophila*); Pgm, PgmL1, PgmL2, PgmL3a/b/c, PgmL4a/b, PgmL5a/b (*Paramecium tetraurelia*); Tru (*Takifugu rubripes*); Pgbd2, Pgbd3 and Pgbd4 (*Homo sapiens*).

Reminiscent of *Paramecium tetraurelia*, which, among ten PiggyMac homologs, has just one homolog with a complete catalytic triad (Bischerour et al., 2018), the DDD triad is preserved in just a single *Blepahrisma* PiggyBac homolog (Figure 5B; Contig_49.g1063, BSTOLATCC_MAC17466). This gene is strongly upregulated during development from 22 to 38 h, when new MACs develop and IES excision is required (Figure 5B). In a multiple sequence alignment the canonical catalytic triad second aspartate of a lower-expressed, MIC-limited PiggyBac is offset by one amino acid (Data S5).

There are significant similarities in the basic properties of *Blepharisma* and *Paramecium* IESs, detailed in the *Blepharisma* MIC genome report (Seah et al. 2022). Consequently, adopting the *Paramecium* nomenclature, we refer to the primary candidate IES excisase as *Blepharisma* PiggyMac (BPgm) and the other somatic homologs as BPgm-Likes (BPgmLs). By extension, we refer to their close relatives which are germline-limited as PiggyMics (Figure 5B).

Other than the PFAM DDE_Tnp_1_7 domain, three *Blepharisma* MAC genome-encoded PiggyBac homologs also possess a short, characteristic cysteine-rich domain (CRD) (Figure 5C), which is absent from the other BPgmLs and PiggyMics. PiggyBac CRDs have been classified into three different groups and are essential for *Paramecium* IES excision (Guérineau et al., 2021). In *Blepharisma*, the CRD consists of five cysteine residues arranged as CxxC-CxxCxxxxH-Cxxx(Y)H (where C, H, Y and x respectively denote cysteine, histidine, tyrosine and any other residue). Two *Blepharisma* homologs possess this CRD without the penultimate tyrosine residue, while the third contains a tyrosine residue before the final histidine. This -YH feature towards the end of the CxxC-CxxCxxxxH-Cxxx(Y)H CRD is shared by all the PiggyBac homologs we found in *Condylostoma*, the bat PiggyBac-like element (PBLE) and human PiggyBac element-derived (PGBD) proteins PGBD2 and PGBD3. In contrast, PiggyBac homologs from *Paramecium* and *Tetrahymena* have a CRD with six cysteine residues arranged in the variants of the motif CxxC-CxxC-Cx{2-7}Cx{3,4}H, and group together with human PGDB4 and *Spodoptera frugiperda* PBLE (Figure 5C).

### PiggyBac transposases are subject to purifying selection and originated early in ciliate evolution

Previous experiments involving individual or paired gene knockdowns of most of the ten *Paramecium tetraurelia* PiggyMac(-like) paralogs led to substantial IES retention, even though only one PiggyMac gene (Pgm) has the complete catalytic triad, indicating that all these proteins are functional (Bischerour et al., 2018). To examine functionally constraints on *Paramecium* PiggyMac homologs we examined non-synonymous (d_N_) to synonymous substitution rates (d_S_), i.e. ω = d_N_/d_S_, for pairwise codon sequence alignments using two closely related *Paramecium* species (*P*. *tetraurelia* and *P*. *octaurelia*). All d_N_/d_S_ values for pairwise comparisons of each of the catalytically incomplete *P. tetraurelia* PgmLs versus the complete Pgm, were less than 1, ranging from 0.01 to 0.25 (Table S7). All d_N_/d_S_ values for pairwise comparisons between *P. tetraurelia* and *P. octaurelia* PiggyBac orthologs were also substantially less than 1, ranging from 0.02 to 0.11 (Table S8). Since d_N_/d_S_= 1 indicates genes evolving neutrally (Yang and Nielsen, 2000), none of these genes are likely pseudogenes, and all appear subject to similar purifying selection.

Only one of *Blepharisma*’s eight MAC and five MIC PiggyBac homologs has the complete, characteristic DDD triad necessary for catalysis. In pairwise comparisons of each of the MAC homologs with incomplete/missing triads versus the complete one d_N_/d_S_ ranges from 0.0076 to 0.1351 (Table S9). The pairwise non-synonymous to synonymous substitution rates of the PiggyMics in comparison to the BPgm were also much less than 1 (range 0.007 to 0.2), indicating they are also subject to purifying selection.

We detected PiggyBac homologs in two other heterotrichs, but not the oligohymenophorean *Ichthyophthirius multifiliis* (“Supplemental information”). To determine whether the *Blepharisma* PiggyBac homologs share a common ciliate ancestor with the oligohymenophorean PiggyBacs, or whether they arose from independent acquisitions in major ciliate groups, we created a large phylogeny of PiggyBac homologs representative of putative domesticated transposases from *Blepharisma stoltei* ATCC 30299, *Condylostoma magnum*, *Paramecium* spp., *Tetrahymena thermophila*, as well as PiggyBac-like elements (PBLEs (Bouallègue et al., 2017)) from diverse eukaryotes (Figure 6; Data S1). All the heterotrichous ciliates PiggyBac homologs, ie. BPgm, BPgmLs 1-7 and PiggyMics grouped together with the *Condylostoma* Pgms. The ciliate Pgms and PgmLs largely cluster as a single clade, with the exception of PiggyMic 5, which appears as a low-support outgroup to opisthokont, archaeplastid and stramenopile PiggyBac-like elements. PiggyMic 5 has the shortest detected DDE_Tnp_1_7 domain (26 a.a.), and appeared poorly aligned relative to the other homologs.

**Figure 6.**
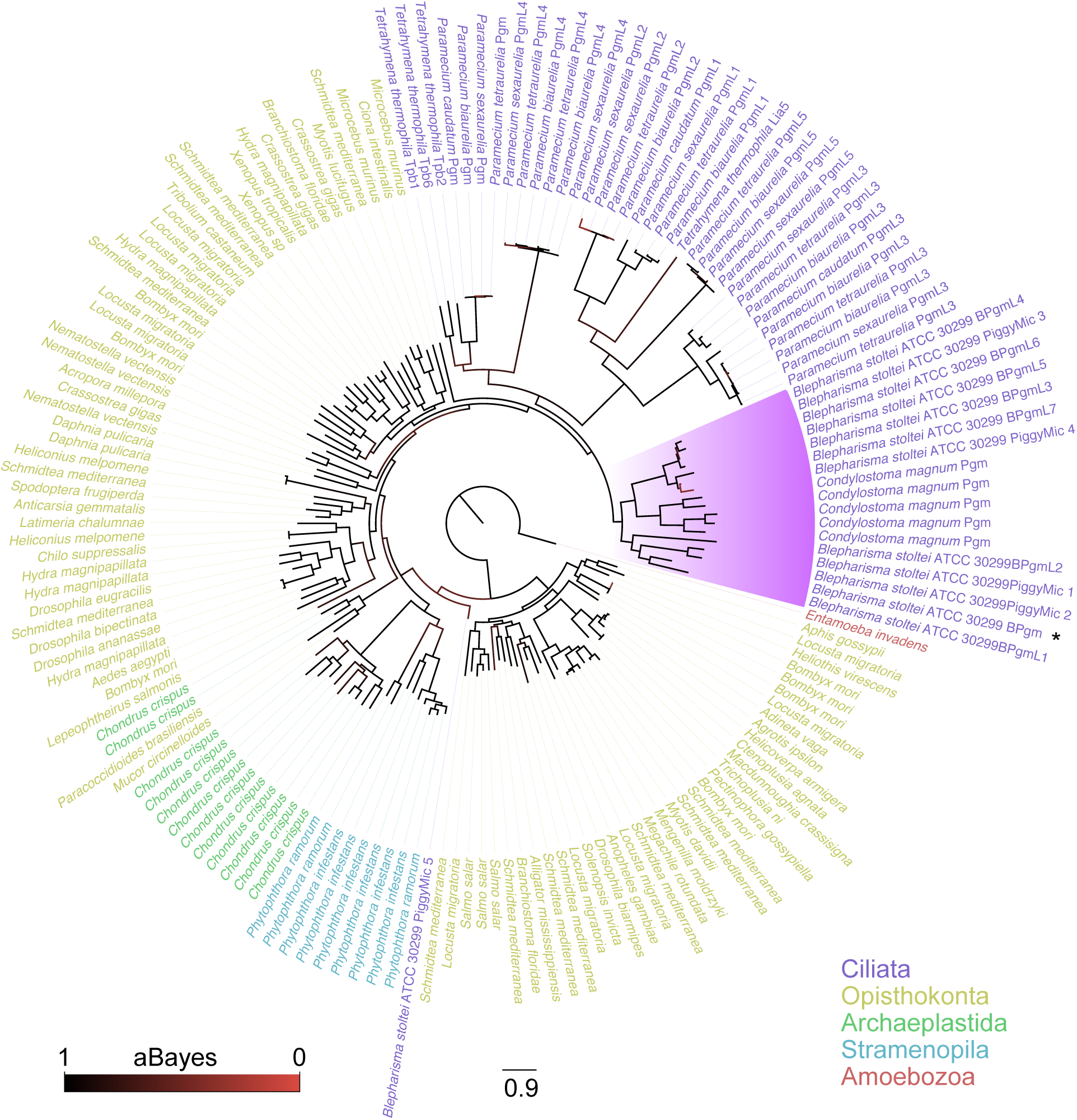
Phylogeny of ciliate PiggyBac homologs, eukaryotic PBLEs and PGBD5 homologs. Highlighted clade contains all PiggyBac homologs found in Heterotrichea, containing MAC and MIC-limited homologs of PiggyMac from *Blepharisma* and PiggyMac homologs of *Condylostoma magnum.* The tree is rooted at the PiggyBac-like element of *Entamoeba invadens*.

### *Blepharisma*’s MAC genome encodes additional domesticated transposases

Three *Blepharisma* MAC genome-encoded proteins possess PFAM domain DDE_1 (PF03184; Figure 7). The most common domain combinations for this domain, aside from proteins with it alone (5898 sequences; PFAM version 35), are with an N-terminal PFAM domain HTH_Tnp_Tc5 (PF03221) alone (2240 sequences), and both an N-terminal CENP-B_N domain (PF04218) and central HTH_Tnp_Tc5 domain (1255 sequences). The CENP-B_N domain is characteristic of numerous transposases, notably the Tigger and PogoR families (Gao et al., 2020). Though pairwise sequence identity is low amongst the *Blepharisma* DDE_1 proteins (avg. 28.3%) in their multiple sequence alignment, the CENP-B_N domain in one of them appears to align reasonably well to corresponding regions in the two proteins lacking this domain, suggesting it deteriorated beyond the recognition capabilities of HMMER3 and the given PFAM domain model. BLASTp matches for all three proteins in GenBank are annotated either as Jerky or Tigger homologs (Jerky transposases belong to the Tigger transposase family (Gao et al., 2020)). Given that none of the *Blepharisma* MAC DDE_1 domain proteins appears to have a complete catalytic triad, it is unlikely they are involved in transposition or IES excision.

**Figure 7.**
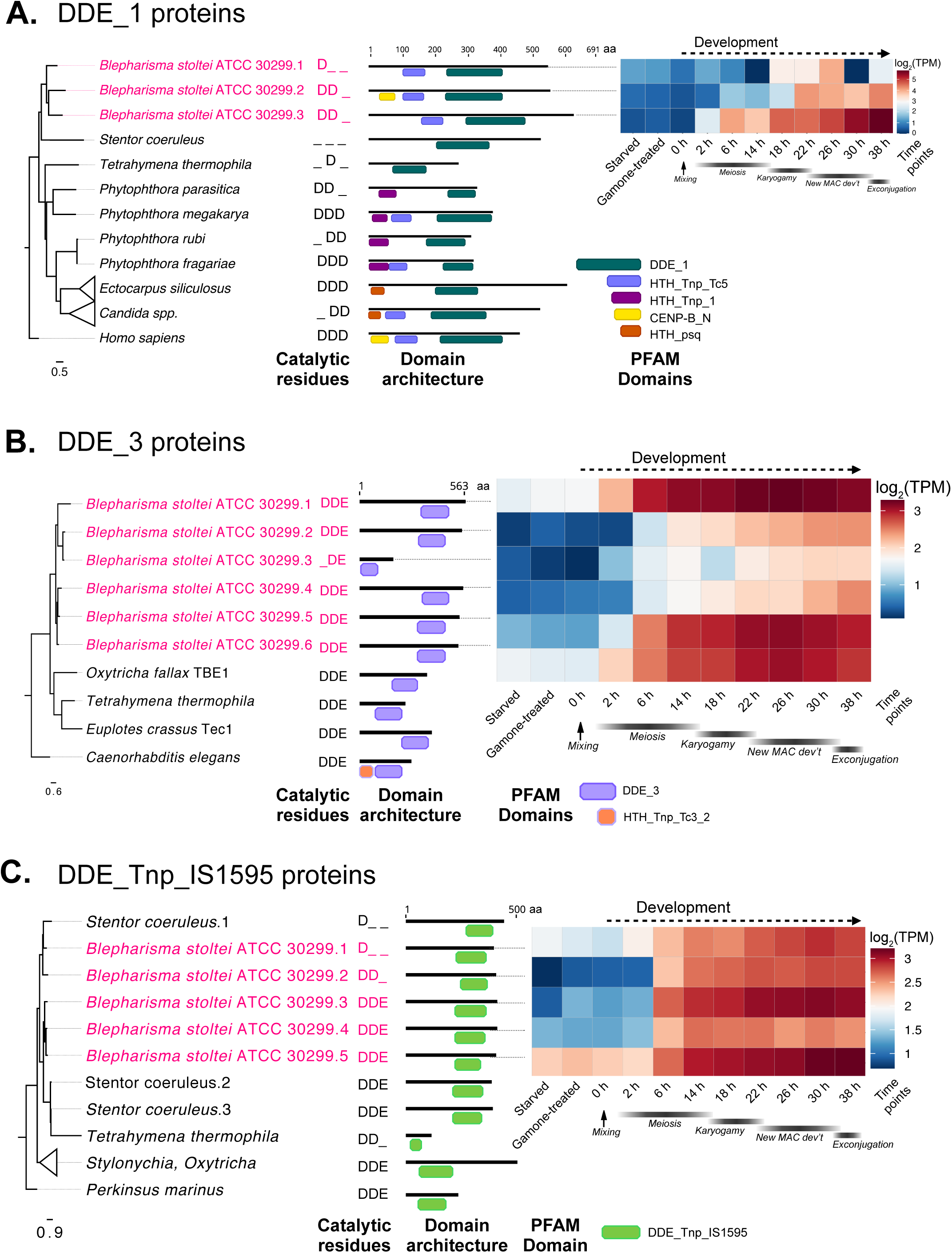
DDE_1, DDE_3 and DDE_Tnp_IS1595 domain-containing proteins in *Blepharisma*. **A.** DDE_1 domain phylogeny with PFAM domain architecture and gene expression heatmap for *Blepharisma*. **B.** DDE_3 domain phylogeny with PFAM domain architecture and gene expression heatmap for *Blepharisma*. **C.** DDE_Tnp_IS1595 domain phylogeny with PFAM domain architecture and gene expression heatmap for *Blepharisma*. See also Figure S7.

Six MAC-encoded transposases containing the DDE_3 domain (PF13358) are present in *Blepharisma*, all of which are substantially upregulated in MAC development and five of which possess the complete DDE catalytic triad (Figure 7B). The DDE_3 domain is characteristic of DDE transposases encoded by the Telomere-Bearing Element transposons (TBEs) of *Oxytricha trifallax* (Williams et al., 1993; Witherspoon et al., 1997), which, despite being MIC genome-limited, are proposed to be involved in IES excision (Nowacki et al., 2009). DDE_3-containing transposons, called Tec elements, are found in another spirotrichous ciliate, *Euplotes crassus,* but no role in genome editing has been established for these (Jahn et al., 1993). TBEs and Tec elements do not share obvious features, other than both possessing an encoded protein belonging to the IS630-Tc1 transposase (super)-family (Doak et al., 1994). All six *Blepharisma* DDE_3 genes have at least 150× HiFi read coverage, consistent with their presence in *bona fide* MAC DNA.

As judged by BLASTP searches in which most of the top hundred best matches are classified are “IS630 family” transposases, *Blepharisma* MAC-encoded DDE_3 domain transposases are more closely related to the IS630 transposase family than to *Oxytricha* TBE transposases and *Euplotes* Tec transposases. One of the BLAST top hits is a MIC genome-encoded protein in *Oxytricha trifallax* with a DDE_3 domain which is not a TBE transposase (GenBank accession: KEJ83017.1). IS630 transposases diverge considerably from Tc1-Mariner transposases, and hence are considered an outgroup to them (Dupeyron et al., 2020). IS630-related transposases encoded by Anchois transposons have also been detected in the *Paramecium tetraurelia* MIC genome (Arnaiz et al., 2012). Given that all but one of the *B. stoltei* paralogs appear to possess a complete catalytic triad, there is a possibility that they may be involved in some IES exicison.

Among other ciliates with draft MAC genomes we examined, the IS1595- and MULE transposase-like domains (PFAM PF12762 and PF10551) have so far only been observed in the spirotrichs *Oxytricha* and *Stylonychia* (Aeschlimann et al., 2014; Swart et al., 2013). DDE_Tnp_IS1595 domains are characteristic of the Merlin transposon superfamily and MULE is part of the Mutator transposon superfamily (Yuan and Wessler, 2011). Currently no particular functions have been demonstrated for these proteins in these ciliates, but their genes were substantially upregulated during their development (Chen et al., 2014; Swart et al., 2013). Both transposase-like domains are found in MAC-encoded proteins in *Blepharisma* and their underlying genes are upregulated during MAC development (Figure 7C, Figure S7). Consistent with the notion of transposase domestication, the genes encoding DDE_Tnp_IS1595 and MULE proteins appear to lack flanking transposon terminal inverted repeats. Members of both IS1595 and MULE transposases also appear to have complete catalytic triads.

In addition to cut-and-paste transposases, we detected a family (> 30 copies) of APE-type non-LTR retrotransposase genes encoding proteins with two characteristic domains, i.e., an APE endonuclease domain (PFAM “exo_endo_phos_2”; PF14529) and a reverse transcriptase domain (PFAM “RVT_1”; PF00078) present on adjacent genes. Unlike the conventional transposase-derived genes in *B. stoltei*, the expression of all these genes throughout the conditions we examined is negligible, and some also appear to be truncated pseudogenes (Data S3; workbook “RVT1 + exo_endo_phos_2”). Since it is necessary to understand the relationship of these sequences with respect to IESs, and that they are not due to residual MIC DNA contamination, their analysis is reported in the context of the *Blepharisma stoltei* MIC genome (Seah et al. 2022).

## Discussion

The genus *Blepharisma* represents one of the earliest diverging ciliate lineages, the heterotrichs, forming an outgroup to the best-studied and deeply divergent oligohymenophorean and spirotrich ciliates (Lynn, 2010). *Blepharisma* species thus provide a vantage point to compare unique processes that have accompanied the evolution of nuclear and genomic dimorphism in ciliates, particularly the extensive genomic editing occurring during MAC development. The annotated draft *B. stoltei* ATCC 30299 MAC genome and associated transcriptomic data provide the basis for comparative studies of genome editing.

### *Blepharisma* PiggyMac is the primary candidate IES excisase

A considerable body of evidence implicates PiggyBac homologs in IES excision of the oligohymenophorean ciliates *Tetrahymena* and *Paramecium* (Arnaiz et al., 2012; Baudry et al., 2009; Bischerour et al., 2018; Cheng et al., 2010; Feng et al., 2017). The responsible IES excisases in the less-studied spirotrichs, *Oxytricha*, *Stylonychia* and *Euplotes*, are not as evident. *Oxytricha*’s TBE transposases are considered to be involved in IES excision, but are encoded by full-length germline-limited transposons and are absent from the MAC (Nowacki et al., 2009), unlike the primary, MAC genome-encoded IES excisase (Tpb2) in *Tetrahymena* and the *Paramecium* PiggyMacs and PiggyMac-likes. The pronounced developmental upregulation of numerous additional MAC- and MIC-encoded transposases in *Oxytricha* raises the possibility that transposases other than those of TBEs could also be involved in IES excision (Chen et al., 2014; Swart et al., 2013). Knowledge of IESs in other ciliates is sparse, primarily confined to the phyllopharyngean *Chilodonella uncinata* (Zufall and Katz, 2007; Zufall et al., 2012). As far as we are aware, no specific IES excisases have been proposed for them.

In current models of IES excision, MIC-limited sequence demarcation by deposition of methylation marks on histones occurs in an sRNA-dependent process (Chalker et al., 2013). These sequences are recognized by domesticated transposases whose excision is supported by additional proteins that somehow recognize these marks (Chalker et al., 2013). Together with MIC sequencing we observed abundant, development-specific sRNA production in *Blepharisma* resembling other model ciliates (Seah et al. 2022). Homologs of proteins implicated in ciliate genome editing were present among the genes most highly differentially upregulated during new MAC development, notably including Dicer-like and Piwi proteins which are candidate genes responsible for development-specific sRNA biogenesis (Figure S8).

Since the oligohymenophorean PiggyBac homologs are clear IES excisases, we sought and found eight homologs of these genes in the *Blepharisma* MAC genome and five in the IESs. *Blepharisma* is the first ciliate genus aside from *Tetrahymena* and *Paramecium* in which such proteins have been reported, and distantly related to both. Additional searches revealed clear PiggyBac homologs in *Condylostoma magnum*, and a weaker pair of matches in *Stentor coeruleus*, suggesting that these are a common feature of heterotrich ciliates. Reminiscent of *Paramecium tetraurelia*, in which just one of the nine PiggyBac homologs, PiggyMac, has a complete DDD catalytic triad (Bischerour et al., 2018), a single *Blepharisma* PiggyBac homolog has a complete canonical DDD catalytic triad, and its gene is highly upregulated during MAC development. As is characteristic of PiggyBac homologs, each of these three proteins also has a C-terminal, cysteine-rich, zinc finger domain. The organization of the heterotrich PiggyBac homolog zinc finger domains is more similar to comparable domains of *Homo sapiens* PGBD2 and PGBD3 homologs than the zinc finger domains in *Paramecium* and *Tetrahymena* PiggyBac homologs.

Since the discovery of multiple PiggyBac homologs (PiggyMac-likes) in *Paramecium* there have been questions about their role. Aside from PiggyMac, all PiggyMac-likes have incomplete catalytic triads, and are thus likely catalytically inactive, but nevertheless their gene knockdowns lead to pronounced IES retention (Bischerour et al., 2018). It has therefore been proposed that the PiggyMac-likes may function as heteromeric multi-subunit complexes in conjunction with PiggyMac during DNA excision (Bischerour et al., 2018). On the other hand, cryo-EM structures available for moth PiggyBac transposase support a model in which these proteins function as a homodimeric complex *in vitro* (Chen et al., 2020). Furthermore, the primary *Tetrahymena* PiggyBac, Tpb2, is able to perform cleavage *in vitro* alone (Cheng et al., 2010). In other eukaryotes, domesticated PiggyBacs without complete catalytic triads are thought to be retained due to co-option of their DNA-binding domains (Sarkar et al., 2003). One possibility for such purely DNA-binding transposase-derived proteins in ciliates could be in competitively regulating (taming) the excision of DNA by the catalytically active transposases. Future experimental analyses of the BPgm and the BPgm-likes could aid in resolving the conundrums and understanding of possible interactions between catalytically active and inactive transposases.

### *Blepharisma* has additional domesticated transposases whose roles await determination

In addition to the PiggyBac homologs, we found MAC genome-encoded transposases with the PFAM domains “DDE_1”, “DDE_3”, “DDE_Tnp_IS1595” and “MULE” in *Blepharisma*. All the genes encoding these proteins lack flanking terminal repeats characteristic of active transposons, suggesting they are further classes of domesticated transposases. In *Blepharisma* and numerous other organisms, the DDE_1 domains co-occur with CENPB domains. Two such proteins represent totally different proposed exaptations in mammals (centromere-binding protein) and fission yeast (regulatory protein) (Casola et al., 2008; Hohmann, 1993; Mojzita and Hohmann, 2006). Given the great evolutionary distances involved, there is no reason to expect that the *Blepharisma* homologs have either function. None of the three proteins with co-occurring DDE_1 and CENPB domains have a complete catalytic triad, making it unlikely that these are active transposases or IES excisases, though all three are noticeably upregulated during MAC development. Six proteins with the PFAM domain DDE_3 are also encoded by *Blepharisma* MAC genes, of which five possess a complete catalytic triad. DDE_3 domains are also characteristic of TBE transposases in *Oxytricha* and Tec transposases in *Euplotes*. All the “DDE_3” protein genes are upregulated during conjugation in *B. stoltei*, peaking during new MAC development. A number of DDE_Tnp_IS1595 and MULE domain-containing proteins have complete catalytic triads and also show pronounced upregulation during *Blepharisma* MAC development.

All ciliate species have MAC genome-encoded transposase families (Figure 5A). Though upregulation of some of these homologs in model ciliates has been noted (Chen et al., 2014; Swart et al., 2013; Vogt and Mochizuki, 2013), their roles remain to be determined. Aside from the timing of IES excisase expression to coincide with new MAC genome formation, the manner in which the excisases perform excision is also crucial. Upon excision, classical cut-and-paste transposases in eukaryotes typically leave behind additional bases, notably including the target-site duplication arising when they were inserted, forming a “footprint” (van Luenen et al., 1994). PiggyBac homologs are unique in performing precise, “seamless” excision in eukaryotes (Elick et al., 1996), conserving the number of bases at the site of transposon insertion after excision, a property that makes them popular for genetic engineering (Chen et al., 2020). *Tetrahymena* Tpb2 is the one exception among PiggyBac homologs associated with imprecise excision in this eukaryote (Cheng et al., 2010). Since intragenic IESs are abundant in *Blepharisma*, like *Paramecium* and unlike *Tetrahymena*, it is essential that these are excised precisely.

Though there are clearly numerous additional domesticated transposases with complete catalytic triads and whose genes are substantially upregulated during *Blepharisma* development, whether they are capable of excision, and if this is precise, needs to be established. *Tetrahymena* has distinct domesticated transposases that excise different subsets of IESs, namely those that are predominant, imprecisely excised and intergenic (by Tpb2) (Cheng et al., 2010), versus those that are rare, precisely excised and intragenic (by Tpb1 and Tpb6) (Cheng et al., 2016; Feng et al., 2017). We could envisage if the additional *Blepharisma* domesticated transposases are still capable of excision, but not a precise form, an involvement in excision of a subset of the numerous intergenic IESs.

### A single origin of PiggyBac homologs within ciliates is the most parsimonious scenario

Though phylogenetic analyses indicate *Tetrahymena*, *Paramecium* and *Blepharisma* PiggyBac homologs form a monophyletic clade the lack of PiggyBac homologs in some ciliate classes and potentially the oligohymenophorean *Ichthyophthirius multifiliis* raises the question whether PiggyBac IES excisases were lost or replaced in these lineages, or rather gained independently from the same source by heterotrichs and a subset of oligohymenophoreans. We think the former is more likely, and consistent with a long-standing hypothesis for ancestral IES excisase substitution in particular ciliate lineages (Klobutcher and Herrick, 1997). However, the alternative cannot be dismissed, because non-model ciliates, where the genome assembly quality allows reliable gene and domain annotations, have only been sparsely sampled.

### Future directions

The *B. stoltei* ATCC 30299 MAC genome together with the corresponding MIC genome (Seah et al., 2022) pave the way for future investigations of genome editing in the context of a peculiar, direct pathway to new MAC genome development skipping the upstream complexity of the standard pathway (Miyake et al., 1991). The pair of *B. stoltei* strains used are both now low frequency selfers, in which the conventional, indirect MAC development pathway dominates. Comparisons with fresh, high frequency *Blepharisma* selfers collected from the wild will facilitate comparative gene expression analyses with the direct MAC development pathway, which will assist in distinguishing expression upregulation due to meiotic and fertilization processes preceding indirect new MAC development.

## Methods

### Strains and localities

The strains used and their original isolation localities were: *Blepharisma stoltei* ATCC 30299, Lake Federsee, Germany (Repak, 1968); *Blepharisma stoltei* HT-IV, Aichi prefecture, Japan; *Blepharisma japonicum* R1072, from an isolate from Bangalore, India (Harumoto et al., 1998).

### Cell cultivation, harvesting and cleanup

For genomic DNA isolation *B. stoltei* ATCC 30299 and HT-IV cells were cultured in Synthetic Medium for *Blepharisma* (SMB) (Miyake and Beyer, 1973) at 27°C. Belpharismas were fed *Chlorogonium elongatum* grown in Tris-acetate phosphate (TAP) medium (Andersen, 2004) at room temperature. *Chlorogonium* cells were pelleted at 1500 g at room temperature for 3 minutes to remove most of the TAP medium, and resuspended in 50 mL SMB. 50 ml of dense *Chlorogonium* was used to feed 1 litre of *Blepharisma* culture once every three days.

*Blepharisma stoltei* ATCC 30299 and HT-IV cells used for RNA extraction were cultured in Lettuce medium inoculated with *Enterbacter aerogenes* and maintained at 25°C (Miyake et al., 1990).

*Blepharisma* cultures were concentrated by centrifugation in pear-shaped flasks at 100 g for 2 minutes using a Hettich Rotanta 460 centrifuge with swing out buckets. Pelleted cells were washed with SMB and centrifuged again at 100 g for 2 minutes. The washed pellet was then transferred to a cylindrical tube capped with a 100 µm-pore nylon membrane at the base and immersed in SMB to filter residual algal debris from the washed cells. The cells were allowed to diffuse through the membrane overnight into the surrounding medium. The next day, the cylinder with the membrane was carefully removed while attempting to minimize dislodging any debris collected on the membrane. Cell density after harvesting was determined by cell counting under the microscope.

### DNA isolation, library preparation and sequencing

*B. stoltei* macronuclei were isolated by sucrose gradient centrifugation (Lauth et al., 1976). DNA was isolated with a Qiagen 20/G genomic-tip kit according to the manufacturer’s instructions. Purified DNA from the isolated MACs was fragmented, size selected and used to prepare libraries according to standard PacBio HiFi SMRTbell protocols. The libraries were sequenced in circular consensus mode to generate HiFi reads.

Total genomic DNA from *B. stoltei* HT-IV and *B. stoltei* ATCC 30299 was isolated with the SigmaAldrich GenElute Mammalian genomic DNA kit. A sequencing library was prepared with a NEBnext FS DNA Library Prep Kit for Illumina and sequenced on an Illumina HiSeq 3000 sequencer, generating 150 bp paired-end reads.

Total genomic DNA from *B. japonicum* was isolated with the Qiagen MagAttract HMW DNA kit. A long-read PacBio sequencing library was prepared using the SMRTbell® Express Template Preparation Kit 2.0 according to the manufacturers’ instructions and sequenced on an PacBio Sequel platform with 1 SMRT cell. Independently, total genomic DNA form *B. japonicum* was isolated with the SigmaAldrich GenElute Mammalian genomic DNA kit and an sequencing library was prepared with the TruSeq Nano DNA Library Prep Kit (Illumina) and sequenced on an Illumina NovaSeq6000 to generate 150 bp paired-end reads.

### Gamone 1/ Cell-Free Fluid (CFF) isolation and conjugation activity assay

*B. stoltei* ATCC 30299 cells were cultured and harvested and concentrated to a density of 2000 cells/mL according to the procedure described in “Cell cultivation, Harvesting and Cleanup”. This concentrated cell culture was incubated overnight at 27°C. The next day, the cells were harvested, and the supernatant collected and preserved at 4°C at all times after extraction. The supernatant was then filtered through a 0.22 µm-pore filter. BSA (10 mg/mL) was added to produce the final CFF at a final BSA concentration of 0.01%.

To assess the activity of the CFF, serial dilutions of the CFF were made to obtain the gamone activity in terms of units (U) (Miyake, 1981).The activity of the isolated CFF was 2^10^ U.

### Conjugation time course and RNA isolation for high-throughput sequencing

*B. stoltei* cells for the complementary strains, ATCC 30299 and HT-IV, were cultivated and harvested by gentle centrifugation to achieve a final cell concentration of 2000 cells/ml for each strain. Non-gamone treated ATCC 30299 (A1) and HT-IV cells (H1) were collected (time point: - 3 hours). Strain ATCC 30299 cells were then treated with synthetic gamone 2 (final concentration 1.5 µg/mL) and strain HT-IV cells were treated with cell-free fluid with a gamone 1 activity of ∼2^10^ U/ml for three hours (Figure S6).

Homotypic pair formation in both cultures was checked after three hours. More than 75% of the cells in both cultures formed homotypic pairs. At this point the samples A2 (ATCC 30299) and H2 (HT-IV) were independently isolated for RNA extraction as gamone-treated control cells just before mixing. For the rest of the culture, homotypic pairs in both cultures were separated by pipetting them gently with a wide-bore pipette tip. Once all pairs had been separated, the two cultures were mixed together. This constitutes the experiment’s 0-h time point. The conjugating culture was observed and samples collected for RNA isolation or cell fixation at 2 h, 6 h, 14 h, 18 h, 22 h, 26 h, 30 h and 38 h (Figure S6). Further details of the sample staging approach are described in (Miyake et al., 1991) and (Sugiura et al., 2012). At each time point including samples A1, H1, A2 and H2, 7 mL of culture was harvested for RNA-extraction using Trizol. The total RNA obtained was then separated into a small RNA fraction < 200 nt and a fraction with RNA fragments > 200 nt using the Zymo RNA Clean and Concentrator-5 kit according to the manufacturer’s instructions. RNA-seq libraries were prepared by BGI according to their standard protocols and sequenced on a BGISeq 500 instrument.

Separate 2 mL aliquots of cells at each time point for which RNA was extracted were concentrated by centrifuging gently at 100 rcf. 50 µL of the concentrated cells were fixed with Carnoy’s fixative (ethanol:acetic acid, 6:1), stained with DAPI and imaged to determine the state of nuclear development (Miyake et al., 1991).

### Cell fixation and imaging

*B. stoltei* cells were harvested as above (“Cell cultivation”), and fixed with an equal volume of “ZFAE” fixative, containing zinc sulfate (0.25 M, Sigma Aldrich), formalin, glacial acetic acid and ethanol (Carl Roth), freshly prepared by mixing in a ratio of 10:2:2:5. Fixed cells were pelleted (1000 g; 1 min), resuspended in 1% TritonX-100 in PHEM buffer to permeabilize (5 min; room temperature), pelleted and resuspended in 2% (w/v) formaldehyde in PHEM buffer to fix further (10 min; room temp.), then pelleted and washed twice with 3% (w/v) BSA in TBSTEM buffer (∼10 min; room temp.). For indirect immunofluorescence, washed cells were incubated with primary antibody rat anti-alpha tubulin (Abcam, ab6161; 1:100 dilution in 3% w/v BSA/TBSTEM; 60 min; room temp.) then secondary antibody goat anti-rat IgG H&L labeled with AlexaFluor 488 (Abcam, ab150157, 1:500 dilution in 3% w/v BSA/TBSTEM; 20 min; room temp.). Nuclei were counterstained with DAPI (1 µg/mL) in 3% (w/v) BSA/TBSTEM. A z-stack of images was acquired using a confocal laser scanning microscope (Leica TCS SP8), equipped with a HC PL APO 40× 1.30 Oil CS2 objective and a 1 photomultiplier tube and 3 HyD detectors, for DAPI (405 nm excitation, 420-470 nm emission) and Alexa Fluor 488 (488 nm excitation, 510-530 nm emission). Scanning was performed in sequential exposure mode. Spatial sampling was achieved according to Nyquist criteria. ImageJ (Fiji) (Schindelin et al., 2012) was used to adjust image contrast and brightness and overlay the DAPI and AlexaFluor 488 channels. The z-stack was temporally color-coded.

For a nuclear 3D reconstruction (Figure 1B), cells were fixed in 1% (w/v) formaldehyde and 0.25% (w/v) glutaraldehyde. Nuclei were stained with Hoechst 33342 (Invitrogen) (5 µM in the culture media), and imaged with a confocal laser scanning microscope (Zeiss, LSM780) equipped with an LD C-Apochromat 40x/1,1 W Korr objective and a 32 channel GaAsP array detector, with 405 nm excitation and 420-470 nm emission. Spatial sampling was achieved according to Nyquist criteria. The IMARIS (Bitplane) software v8.0.2 was used for three-dimensional reconstructions and contrast adjustments.

### Genome assembly

Two MAC genome assemblies for *B. stoltei* ATCC 30299 (70× and 76× coverage) were produced with Flye (version 2.7-b1585) (Kolmogorov et al., 2019) for the two separate PacBio Sequel II libraries (independent replicates) using default parameters and the switches: --pacbio-hifi -g 45m. The approximate genome assembly size was chosen based on preliminary Illumina genome assemblies of approximately 40 Mb. Additional assemblies using the combined coverage (145×) of the two libraries were produced using either Flye version 2.7-b1585 or 2.8.1- b1676, and the same parameters. Two rounds of extension and merging were then used, first comparing the 70× and 76× assemblies to each other, then comparing the 145× assembly to the former merged assembly. Assembly graphs were all relatively simple, with few tangles to be resolved (Figure S5B). Minimap2 (Li, 2018) was used for pairwise comparison of the assemblies using the parameters: -x asm5 --frag=yes --secondary=no, and the resultant aligned sequences were visually inspected and manually merged or extended where possible using Geneious (version 2020.1.2) (Kearse et al., 2012).

Visual inspection of read mapping to the combined assembly was then used to trim off contig ends where there was little correspondence between the assembly consensus and the mapped reads - which we classify as “cruft”. Read mapping to cruft regions was often lower or uneven, suggestive of repeats. Alternatively, these features could be due to trace MIC sequences, or sites of alternative chromosome breakage during development which lead to sequences that are neither purely MAC nor MIC. A few contigs with similar dubious mapping of reads at internal locations, which were also clear sites of chromosome fragmentation (evident by abundant telomere-bearing reads in the vicinity) were split apart and trimmed back as for the contig ends. Telomere-bearing reads mapped to the non-trimmed region nearest to the trimmed site were then used to define contig ends, adding representative telomeric repeats from one of the underlying sequences mapped to each of the ends. The main genome assembly with gene predictions can be obtained from the European Nucleotide Archive (ENA) (PRJEB40285; accession GCA_905310155). “Cruft” sequences are also available from the same accession.

Two separate assemblies were generated for *Blepharisma japonicum.* A genome assembly for *Blepharisma japonicum* strain R1072 was generated from Illumina reads, using SPAdes genome assembler (v3.14.0) (Prjibelski et al., 2020). An assembly with PacBio Sequel long reads was produced with Ra (v0.2.1) (Vaser and Sikic, 2019), which uses the Overlap-Layout-Consensus paradigm. The assembly produced with Ra was more contiguous, with 268 contigs, in comparison to 1510 contigs in the SPAdes assembly, and was chosen as the reference assembly for *Blepharisma japonicum* (ENA accession: ERR6474383).

*Condylostoma magnum* genomic reads (study accession PRJEB9019) from a previous study (Swart et al., 2016) were reassembled to improve contiguity and remove bacterial contamination. Reads were trimmed with bbduk.sh from the BBmap package v38.22 (https://sourceforge.net/projects/bbmap/), using minimum PHRED quality score 2 (both ends) and k-mer trimming for Illumina adapters and Phi-X phage sequence (right end), retaining only reads ≥25 bp. Trimmed reads were error-corrected and reassembled with SPAdes v3.13.0 (Prjibelski et al., 2020) using k-mer values 21, 33, 55, 77, 99. To identify potential contaminants, the unassembled reads were screened with phyloFlash v3.3b1 (Gruber-Vodicka et al., 2020) against SILVA v132 (Quast et al., 2013); the coding density under the standard genetic code and prokaryotic gene model were also estimated using Prodigal v2.6.3 (Hyatt et al., 2010). Plotting the coverage vs. GC% of the initial assembly showed that most of the likely bacterial contigs (high prokaryotic coding density, lower coverage, presence of bacterial SSU rRNA sequences) had >=40% GC, so we retained only contigs with <40% GC as the final *C. magnum* genome bin. The final assembly is available from the ENA bioproject PRJEB48875 (accession GCA_920105805).

All assemblies were inspected with the quality assessment tool QUAST (Gurevich et al., 2013).

### Variant calling

Illumina total genomic DNA-seq libraries for *B. stoltei* strains ATCC 30299 (ENA accession: ERR6061285) and HT-IV (ERR6064674) were mapped to the ATCC 30299 reference assembly with bowtie2 v2.4.2 (Langmead and Salzberg, 2012). Alignments were tagged with the MC tag (CIGAR string for mate/next segment) using samtools (Danecek et al., 2021) fixmate. The BAM file was sorted and indexed, read groups were added with bamaddrg (commit 9baba65, https://github.com/ekg/bamaddrg), and duplicate reads were removed with Picard MarkDuplicates v2.25.1 (http://broadinstitute.github.io/picard/). Variants were called from the combined BAM file with freebayes v1.3.2 (Garrison and Marth, 2012) in diploid mode, with maximum coverage 1000 (option -g). The resultant VCF file was combined and indexed with bcftools v1.12 (Danecek et al., 2021), then filtered to retain only SNPs with quality score > 20, and at least one alternate allele.

### Annotation of alternative telomere addition sites

Alternative telomere addition sites (ATASs) were annotated by mapping PacBio HiFi reads to the curated reference MAC assembly described above, using minimap2 and the following flags: -x asm20 --secondary=no --MD. We expect reads representing alternative telomere additions to have one portion mapping to the assembly (excluding telomeric regions), with the other portion containing telomeric repeats being soft-clipped in the BAM record. For each mapped read with a soft-clipped segment, we extracted the clipped sequence, and the coordinates and orientation of the clip relative to the reference. We searched for ≥ 24 bp tandem direct repeats of the telomere unit (i.e., ≥3 repeats of the 8 bp unit) in the clipped segment with NCRF v1.01.02 (Harris et al., 2019), which can detect tandem repeats in the presence of noise, e.g., from sequencing error. The orientation of the telomere sequence, the distance from the end of the telomeric repeat to the clip junction (‘gap’), and the number of telomere-bearing reads vs. total mapped reads at each junction were also recorded. Junctions with zero gap between telomere repeat and clip junction were annotated as ATASs. The above procedure was implemented in the MILTEL module of the software package BleTIES v0.1.3 (Seah and Swart, 2021).

MILTEL output was processed with Python scripts depending on Biopython (Cock et al., 2009), pybedtools (Dale et al., 2011), Bedtools (Quinlan and Hall, 2010), and Matplotlib (Hunter, 2007), to summarize statistics of junction sequences and telomere permutations at ATAS junctions, and to extract genomic sequences flanking ATASs for sequence logos. Logos were drawn with Weblogo v3.7.5 (Crooks et al., 2004), with sequences oriented such that the telomere would be added on the 5’ end of the ATAS junctions.

To calculate the expected minichromosome length, we assumed that ATASs were independent and identically distributed in the genome following a Poisson distribution. About 47×10^3^ ATASs were annotated, supported on average by a single read. Given a genome of 42 Mbp at 145× coverage, the expected rate of encountering an ATAS is 47×10^3^ / (145 × 42 Mbp), so the distance between ATASs (i.e., the minichromosome length) is exponentially distributed with expectation (145 × 42 Mbp) / 47×10^3^ = 130 kbp.

### RNA-seq read mapping

To permit correct mapping of tiny introns RNA-seq data was mapped to the *B. stoltei* ATCC 30299 MAC genome using a version of HISAT2 (Kim et al., 2019) with modified source code, with the static variable minIntronLen in hisat2.cpp lowered to 9 from 20 (change available in the HISAT2 github fork: https://github.com/Swart-lab/hisat2/; commit hash 86527b9). HISAT2 was run with default parameters and parameters --min-intronlen 9 --max-intronlen 500. It should be noted that RNA-seq from timepoints in which *B. stoltei* ATCC 30299 and *B. stoltei* HT-IV cells were mixed together were only mapped to the former genome assembly, and so reads for up to three alleles may map to each of the genes in this assembly.

### Genetic code prediction

We used the program PORC (Prediction Of Reassigned Codons; available from https://github.com/Swart-lab/PORC) previously written to predict genetic codes in protist transcriptomes (Swart et al., 2016) to predict the *B. stoltei* genetic code. This program was used to translate the draft *B. stoltei* ATCC 30299 genome assembly in all six frames (with the standard genetic code). Like the program FACIL (Dutilh et al., 2011) that inspired PORC, the frequencies of amino acids in PFAM (version 34.0) protein domain profiles aligned to the six frame translation by HMMER 3.1b2 (Eddy, 2011) (default search parameters; domains used for prediction with conditional E-values < 1e-20), and correspondingly also to the underlying codon, are used to infer the most likely amino acid encoded by each codon (Figure S1B).

### Gene prediction

We created a wrapper program, Intronarrator, to predict genes in *Blepharisma* and other heterotrichs, accommodating their tiny introns. Intronarrator can be downloaded and installed together with dependencies via Conda from GitHub (https://github.com/Swart-lab/Intronarrator). Intronarrator directly infers introns from spliced RNA-seq reads mapped by HISAT2 from the entire developmental time course we generated. RNA-seq reads densely cover almost the entire *Blepharisma* MAC genome, aside from intergenic regions, and most potential protein-coding genes (Figure 4B). After predicting the introns and removing them to create an intron-minus genome, Intronarrator runs AUGUSTUS (version 3.3.3) using its intronless model. It then adds back the introns to the intronless gene predictions to produce the final gene predictions.

Introns are inferred from “CIGAR” string annotations in mapped RNA-seq BAM files, using the regular expression “[0-9]+M([0-9][0-9])N[0-9]+M” to select spliced reads. For intron inference we only used primary alignments with: MAPQ >= 10; just a single “N”, indicating one potential intron, per read; and at least 6 mapped bases flanking both the 5’ and 3’ intron boundaries (to limit spurious chance matches of a few bases that might otherwise lead to incorrect intron prediction). The most important parameters for Intronarrator are a cut-off of 0.2 for the fraction of spliced reads covering a potential intron, and a minimum of 10 or more spliced reads to call an intron. The splicing fraction cut-off was chosen based on the overall distribution of splicing (Figure S4A-C). From our visual examination of mapped RNA-seq reads and gene predictions, values less than this were typically “cryptic” excision events (Saudemont et al., 2017) which remove potentially essential protein-coding sequences, rather than genuine introns. Intronarrator classifies an intron as sense (7389 in total, excluding alternative splicing), when the majority of reads (irrespective of splicing) mapping to the intron are the same strand, and antisense (554 in total) when they are not. The most frequently spliced intron was chosen in rare cases of overlapping alternative intron splicing.

To eliminate spurious prediction of protein-coding genes overlapping ncRNA genes, we also incorporated ncRNA prediction in Intronarrator. Infernal (Nawrocki et al., 2009) (default parameters; e-value < 1e-6) was used to predict a restricted set of conserved ncRNAs models (i.e., tRNAs, rRNAs, SRP, and spliceosomal RNAs) from RFAM 14.0 (Kalvari et al., 2018). These ncRNAs were hard-masked (with “N” characters) before AUGUSTUS gene prediction. Both Infernal ncRNA predictions (excluding tRNAs) and tRNA-scan SE 2.0 (Chan et al., 2019) (default parameters) tRNA predictions are annotated in the *B. stoltei* ATCC 30299 assembly deposited in the European Nucleotide Archive.

Since we found that *Blepharisma stoltei*, like *Blepharisma japonicum* (Swart et al., 2016), uses a non-standard genetic code, with UGA codon translated as tryptophan, gene predictions use the “The Mold, Protozoan, and Coelenterate Mitochondrial Code and the Mycoplasma/Spiroplasma Code (transl_table=4)” from the NCBI genetic codes. The default AUGUSTUS gene prediction parameters override alternative (mitochondrial) start codons permitted by NCBI genetic code 4, other than ATG. So, all predicted *B. stoltei* gene coding sequences begin with ATG.

RNA-seq read mapping relative to gene predictions of Contig_1 of *B. stoltei* ATCC30299 was visualized with PyGenomeTracks (Lopez-Delisle et al., 2021).

### Assessment of genome completeness

A BUSCO (version 4.0.2) (Waterhouse et al., 2018) analysis of the assembled MAC genomes of *B. stoltei* and *B. japonicum* was performed on the set of predicted proteins (BUSCO mode -prot) using the BUSCO Alveolata database. The completeness of the *Blepharisma* genomes was compared to the protein-level BUSCO analysis of the published genome assemblies of ciliates *T. thermophila*, *P. tetraurelia*, *S. coeruleus* and *I. multifiliis* (Figure S1).

### Gene annotation

Pannzer2 (Törönen et al., 2018) (default parameters) and EggNog (version 2.0.1) (Huerta-Cepas et al., 2019) were used for gene annotation. Annotations were combined and are available from the Max Planck Society’s Open Research Repository, Edmond (https://dx.doi.org/10.17617/3.8c). Protein domain annotations were performed using hmmscan from HMMER3 (version 3.3, Nov 2019) (Eddy, 2011) vs. the PFAM database (Pfam-A.full, 33.0, retrieved on June 23, 2020) with default parameters.

### Gene expression analysis

Features from RNA-seq reads mapped to the *B. stoltei* ATCC 30299 MAC and MAC+IES genomes over the developmental time-course were extracted using featureCounts from the Subread package (Liao et al., 2014). Further analysis was performed using the R software environment. Genes with a total read count of less than 50, across all timepoints, were filtered out of the dataset. The remaining genes were passed as a DGElist object to edgeR (Robinson et al., 2010). Each time point, representing one library, was normalized for library size using the edgeR function calcNormFactors. The normalized read counts were transformed into TPM (transcripts per million) values (Li et al., 2010; Wagner et al., 2012). The TPM-values for different genes were compared across timepoints to examine changes in gene expression. Heatmaps showing log2(TPM) changes across timepoints were plotted using the tidyverse collection of R packages (https://www.tidyverse.org/) and RColorBrewer (https://rdrr.io/cran/RColorBrewer/). Tabulated gene expression estimates together with protein annotations are available from Edmond (https://dx.doi.org/10.17617/3.8c).

### Sequence visualization and analysis

Nucleotide and amino acid sequences were visualized using Geneious Prime (Biomatters Ltd.) (Kearse et al., 2012). Multiple sequence alignments were performed with MAFFT version 7.450 (Katoh and Standley, 2013; Katoh et al., 2002). Phylogenetic trees were constructed with PhyML version 3.3.20180621 (Guindon et al., 2010).

### Orthogroup inference and analysis of orthogroup clusters

OrthoFinder version 2.5.2 with default parameters (i.e., using Diamond for searching, MAFFT for multiple alignment and FastTree for phylogenies) was used to define orthogroups, i.e., sets of genes descended from the last common ancestor of the chosen species. Proteomes for the following ciliate species were used: *Tetrahymena thermophila*, *Oxytricha trifallax*, *Stentor coeruleus* (data from ciliate.org (Stover et al., 2012)); *Euplotes octocarinatus* (EOGD (Wang et al., 2018)); *Paramecium tetraurelia*, *Paramecium caudatum* (data from ParameciumDB (Arnaiz et al., 2020)); plus *Perkinsus marinus* ATCC 50983 (GenBank accession: AAXJ00000000) as a non-ciliate outgroup. Orthogroup clusters are available as Data S2, or from Edmond (https://dx.doi.org/10.17617/3.8c).

### Identification and correction of MIC-encoded PiggyBac homologs

We sought coding regions present within *Blepharisma* IESs to gauge the expression and type of MIC-limited genes (IES assembly and gene prediction described in Seah et al. 2022). After gene prediction within IESs with Intronarrator, predicted protein domains were annotated by HMMER (v3.3) (Eddy, 2011). Several transposase families were represented in protein domains identified with coding regions of IESs. However, gene prediction within IESs was hampered by the presence of intermittent A-residues in the consensus sequence which occur due to the inaccuracy inherent in long-reads, from which the IES regions were assembled. These errors cause IES gene-prediction to falter by generating inaccurate ORFs. To circumvent this, a six-frame translation of the MIC-limited genome regions was performed using a custom script, which was then used to detect PFAM domains, using HMMER and the Pfam-A database 32.0 (release 9) (Mistry et al., 2021). Domain annotations for diagrams were generated with the InterproScan 5.44-79.0 pipeline (Jones et al., 2014)

Four instances of the Pfam domain DDE_Tnp_1_7, characteristic of PiggyBac transposases, were detected in an initial gene prediction within *Blepharisma* IESs. The four genes corresponding to the DDE_Tnp_1_7 domain had high RNA-seq coverage of combined reads from all timepoints across development. The IESs with the PiggyBac domains on Contig 17 and Contig 39 each had two ORFs with a partial DDE_1_7 domain, separated by a few hundred bp. Alignment of short-read MIC-enriched DNA reads mapped to the IES regions containing the putative PiggyBac homologs indicated that several A-nucleotides in the assembled IESs were insertion errors in the IES assembly, which were corrected with the short-read alignment. Open reading frames of predicted genes in these corrected regions were adjusted accordingly. The prefix “cORF” (corrected ORFs) was used to indicate the short-read corrected sequences of the PiggyMics.

Short-read MIC-enriched DNA sequences were aligned to the IES regions containing putative PiggyBac homologs with Hisat2 (2.0.0-beta) with modified source code (described above). Indel errors in the IES assembly were corrected manually, then used to predict coding regions. Pfam domains were annotated on MIC PiggyBac homologs with corrected ORFs using the InterproScan (v. 1.1.4) (Quevillon et al., 2005) plugin in Geneious v11.1.5 (Biomatter Ltd.). DDE_Tnp_1_7 domains were detected in the corrected ORFs, which in some cases spanned IES regions lacking predicted genic regions before correction. A multiple sequence alignment of the correct MIC PiggyBac homologs with other ciliate PiggyBac-derived proteins (PGBDs) and eukaryotic PiggyBac-like elements (PBLEs) that contain the PiggyBac transposase domain DDE_Tnp_1_7 (PF13843) was performed with MAFFT (v4.1) via the Geneious plugin (algorithm L-INS-i, BLOSUM62 scoring matrix, gap open penalty 1.53, offset value 0.123). A phylogenetic tree was constructed using the FastTree (v 2.1.11) plugin for Geneious (Whelan-Goldman model).

### d_N_/d_S_ estimation

We generated pairwise coding sequence alignments of PiggyMac paralog nucleotide sequences from *P. tetraurelia* and *P. octaurelia* using MAFFT version 7.450 (Katoh and Standley, 2013) (Katoh et al., 2002) (algorithm: “auto”, scoring matrix: 200PAM/k=2, gap open penalty 1.53, offset value 0.123) using the “translation align” panel of Geneious Prime (version 2020.1.2) (Kearse et al., 2012). PAML version 4.9 (Yang, 2007) was used to estimate d_N_/d_S_ values in pairwise mode (runmode = -2, seqtype = 1, CodonFreq = 2). For *Blepharisma stoltei*, we generated pairwise coding sequence alignments of the *Blepharisma* PiggyMac homolog, BPgm (Contig_49.g1063; BSTOLATCC_MAC17466), with the *Blepharisma* Pgm-likes (BPgmLs) using Translation Align panel of Geneious v11.1.5 (Genetic code: *Blepharisma*, Protein alignment options: MAFFT alignment (v7.450) (Katoh and Standley, 2013), scoring matrix: BLOSUM62, Gap open penaly: 1.53, offset value: 0.1). PAML version 4.9 was used to estimate dN/dS values in pairwise mode (runmode = -2, seqtype = 1, CodonFreq = 2).

### Phylogenetic analysis

Protein sequences of PBLEs were obtained from Bouallègue et al (Bouallègue et al., 2017). Protein sequences of *Paramecium* and *Tetrahymena* Pgms and PgmLs were obtained from ParameciumDB (Arnaiz et al., 2020) (PGM, PGMLs1-5) and ciliate.org (Stover et al., 2012) (Tpb1, Tpb2, Tpb7, LIA5), respectively. *Condylostoma* and *Blepharisma* Pgms and PgmLs were obtained from genome assemblies (accessions GCA_920105805 and GCA_905310155, respectively). Sequence manipulation was done using Geneious (Biomatters Ltd.). The Geneious plug-in for InterProScan (Jones et al., 2014) was used to identify DDE_Tnp_1_7 domains using the PFAM-A database (Mistry et al., 2021). The DDE_Tnp_1_7 domain and regions adjacent to it were extracted and aligned using the MAFFT plug-in (v7.450) for Geneious (Katoh and Standley, 2013) (Algorithm: L-INS-i, Scoring matrix: BLOSUM62, Gap open penalty: 1.53, Offset value: 0.123). Phylogenetic trees using this alignment were generated with the FastTree2 (v2.2.11) Geneious plug-in using the Whelan-Goldman model. The phylogenetic trees were visualized with FigTree (v1.4.4) (Andrew Rambaut, http://tree.bio.ed.ac.uk/).

### Repeat annotation

Interspersed repeat element families were predicted with RepeatModeler v2.0.1 (default settings, random number seed 12345) with the following dependencies: rmblast v2.9.0+ (http://www.repeatmasker.org/RMBlast.html), TRF 4.09 (Benson, 1999), RECON (Bao and Eddy, 2002), RepeatScout 1.0.6 (Price et al., 2005), RepeatMasker v4.1.1 (http://www.repeatmasker.org/RMDownload.html). Repeat families were also classified in the pipeline by RepeatClassifier v2.0.1 through comparison against RepeatMasker’s repeat protein database and the Dfam database. Consensus sequences of the predicted repeat families, produced by RepeatModeler, were then used to annotate repeats with RepeatMasker, using rmblast as the search engine.

Terminal inverted repeats (TIRs) of selected repeat element families were identified by aligning the consensus sequence from RepeatModeler, and/or selected full-length elements, with their respective reverse complements using MAFFT (Katoh and Standley, 2013) (plugin version distributed with Geneious). TIRs from the Dfam DNA transposon termini signatures database (v1.1, https://www.dfam.org/releases/dna_termini_1.1/dna_termini_1.1.hmm.gz) (Storer et al., 2021) were searched with hmmsearch (HMMer v3.2.1) against the IES sequences, to identify matches to TIR signatures of major transposon subfamilies.

### Data and code availability

The draft *Blepharisma stoltei* ATCC 30299 MAC genome assembly is accessible from bleph.ciliate.org and from the European Nucleotide Archive (ENA) bioproject PRJEB40285 under the accession GCA_905310155. PacBio CCS reads (ERR5873783 and ERR5873334) and subreads (ERR5962314) used to assemble the genome are also available from ENA. Illumina DNA-seq data for the *B. stoltei* ATCC 30299 and HT-IV strains is available from accessions ERR6061285 and ERR6064674, respectively. The RNA-seq developmental time course is available from the bioproject PRJEB45374 (accessions ERR6049461-ERR6049485).

Illumina and PacBio Sequel sequencing data for *Blepharisma japonicum* strain R1702 is available from the ENA bioproject PRJEB46921 (Illumina accessions: ERR6473251, ERR6474356; PacBio accession: ERR6474383).

Code availability for software we generated or modified is indicated in place in Methods.

## Acknowledgements

This paper is dedicated in memory of Akio Miyake and his decades of inspirational *Blepharisma* research. We thank Federico Buonanno for the provision of *B. stoltei* ATCC 30299 cells and culturing advice, Christa Lanz and the MPI for Biology’s genome center, Sebastien Colin and the MPI for Biology’s Light Microscopy Facility for the 3D nuclear reconstruction, and Adrian Streit for discussion. Research reported in this publication was supported by the National Institutes of Health (award No. P40OD010964) to N.A.S and the Max Planck Society.

## Author contributions

Conceptualization, M.S.1., K.B.B.S., E.C.S.; Methodology, M.S.1., K.B.B.S., C.E., A.S., C.W., B.H., E.C.S.; Software, M.S.1., K.B.B.S., E.C.S.; Investigation, M.S.1., K.B.B.S., C.W., B.H., E.C.S.; Writing – Original Draft, M.S.1., K.B.B.S., A.S., C.W., B.H., E.C.S.; Writing – Review & Editing, M.S.1, K.B.B.S., A.S., E.C.S.; Funding Acquisition, E.C.S.; Resources, A.B. and N.A.S.; Supervision, M.S.2., T.H., E.C.S.

## Abbreviations

MIC: micronucleus
MAC: macronucleus
IES: internally eliminated sequence
MDS: macronuclear-destined sequence
PacBio: Pacific Biosciences
CLR: continuous long read (PacBio)
CCS: circular consensus sequence (PacBio)
HiFi: High-fidelity read (PacBio)
ATAS: alternative telomere addition site
PBLE: PiggyBac-like element
PGBD: PiggyBac element-derived
Pgm: PiggyMac
PgmL: PiggyMac-like

## Supplemental figure captions

**Figure S1.**
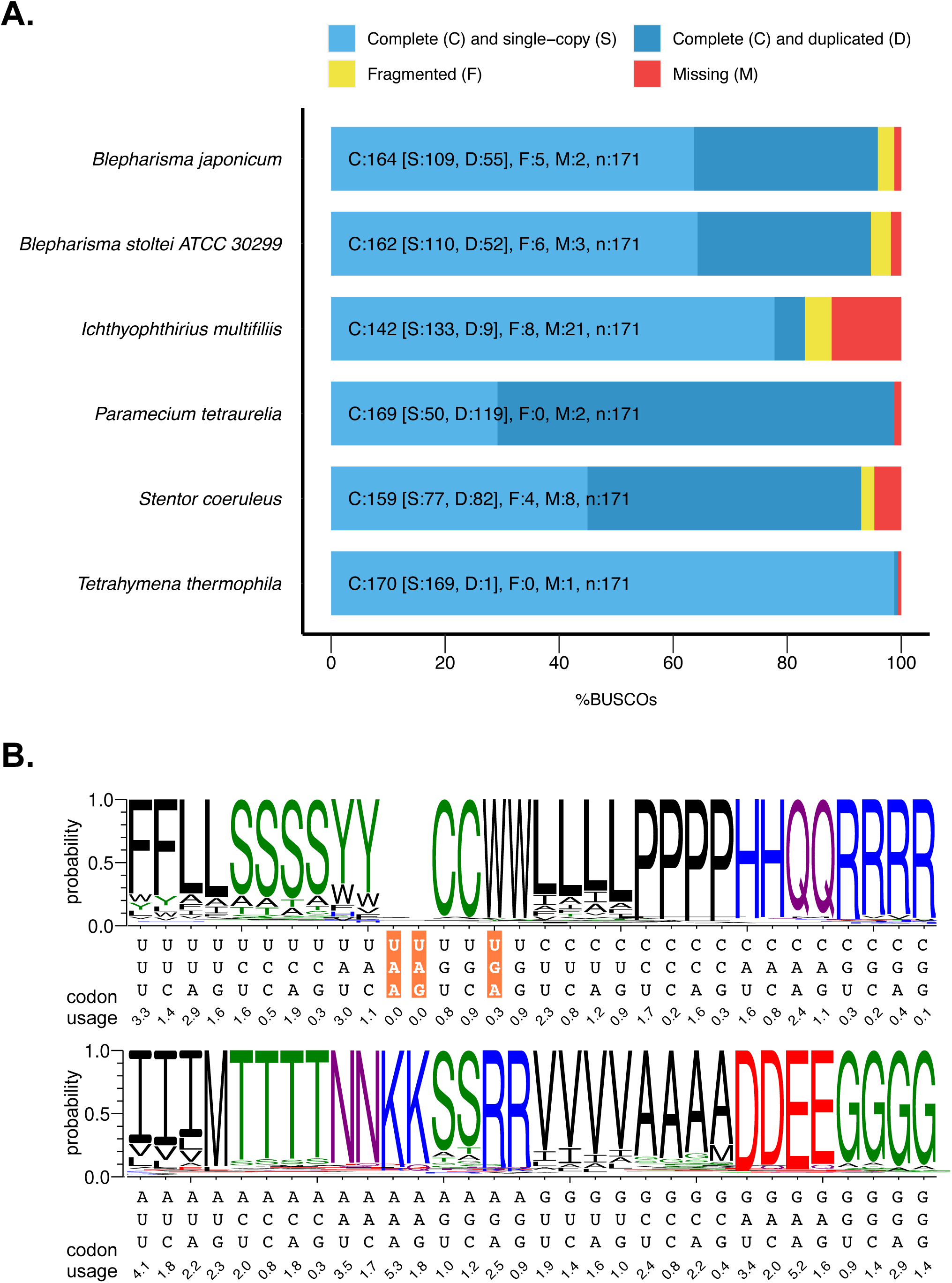
Analysis of assembly completeness and genetic code. **A.** Completeness of the *B. stoltei* ATCC 30299 MAC assembly was estimated by the percentage of BUSCOs found in the assembly with reference to the OrthoDB v10 alveolate database (Kriventseva et al., 2019). The nature of the ortholog-matches is indicated by characters followed by counts: C (complete orthologs) - light blue, D (duplicated orthologs) - dark blue, F (fragmented orthologs) - yellow and M (missing orthologs) - red. **B.** Prediction for *B. stoltei* ATCC 30299 MAC genome by PORC; codons that are stops in the standard genetic code are highlighted in orange.

**Figure S2.**
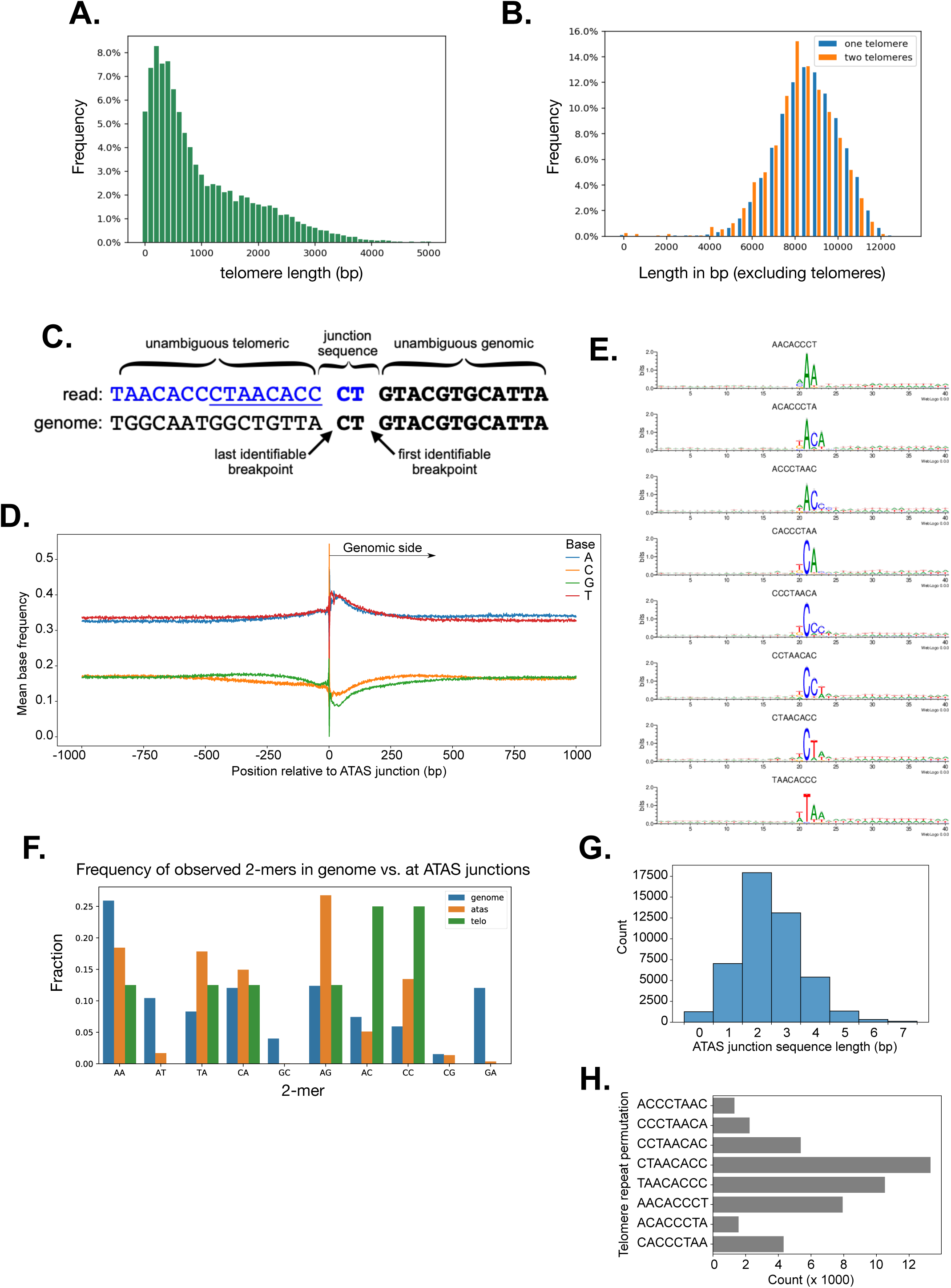
Properties of minichromosomes, telomeres, and alternative telomere addition sites. **A.** Length distribution of telomeres of telomere-bearing HiFi reads. **B.** Length distribution of HiFi reads delimited by telomeres. **C**. Diagram of a telomere-bearing read mapped onto genome reference at an ATAS. Sequence which is ambiguously chromosomal or telomeric is “junction sequence”; junction coordinate which maximizes telomere repeat length on the read is the “first identifiable breakpoint”; the coordinate maximizing alignment length to reference is the “last identifiable breakpoint”. The last telomeric unit permutation at the last identifiable breakpoint is underlined (length 8 bp). **D.** Mean base frequencies in +/- 1 kbp flanking ATAS junctions. **E.** Sequence logos of chromosomal sequence at ATAS junctions, sorted by which permutation of the telomeric repeat is present (plot labels). Logos are aligned to the “last identifiable breakpoint” between positions 20 and 21; telomeric repeats on telomere-bearing reads begin to the left of the breakpoint. **F**. Frequencies of 2-mers in whole genome (blue), in telomeres (green), and at ATAS junctions (chromosomal side after last identifiable breakpoint, orange). **G.** Histogram of junction sequence lengths for ATASs in *B. stoltei*. **H.** Counts of each telomere repeat permutation at ATAS junctions (last identifiable breakpoint).

**Figure S3.**
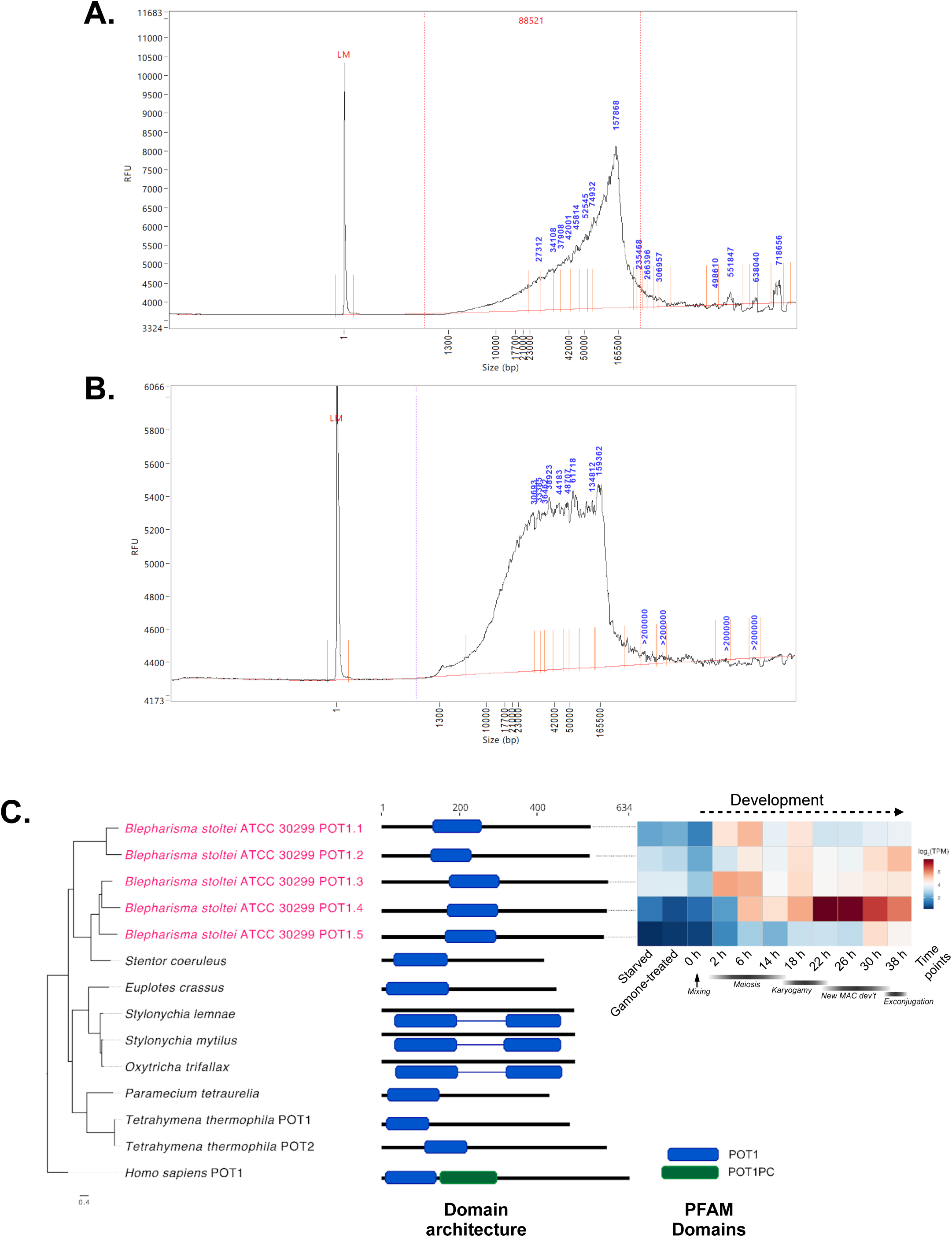
Femto Pulse analyses of *B. stoltei* MAC DNA and POT1 phylogeny. **A.** Mapping of PacBio CLR reads with 3 consecutive telomeric repeats to a representative *T. thermophila* MAC chromosome (Chr_001 from ciliate.org). **B.** Length distribution of input MAC DNA sizes prior to fragmentation and library preparation (Femto Pulse; LM = lower maker) - replicate 1. RFU=relative fluorescent units. **C.** Length distribution of input MAC DNA sizes prior to fragmentation and library preparation (Femto Pulse; LM = lower maker) - replicate 2. **C.** POT1 paralog phylogeny, PFAM domain architecture, and gene expression in *Blepharisma.* Diagram elements as described in Figure 5B.

**Figure S4.**
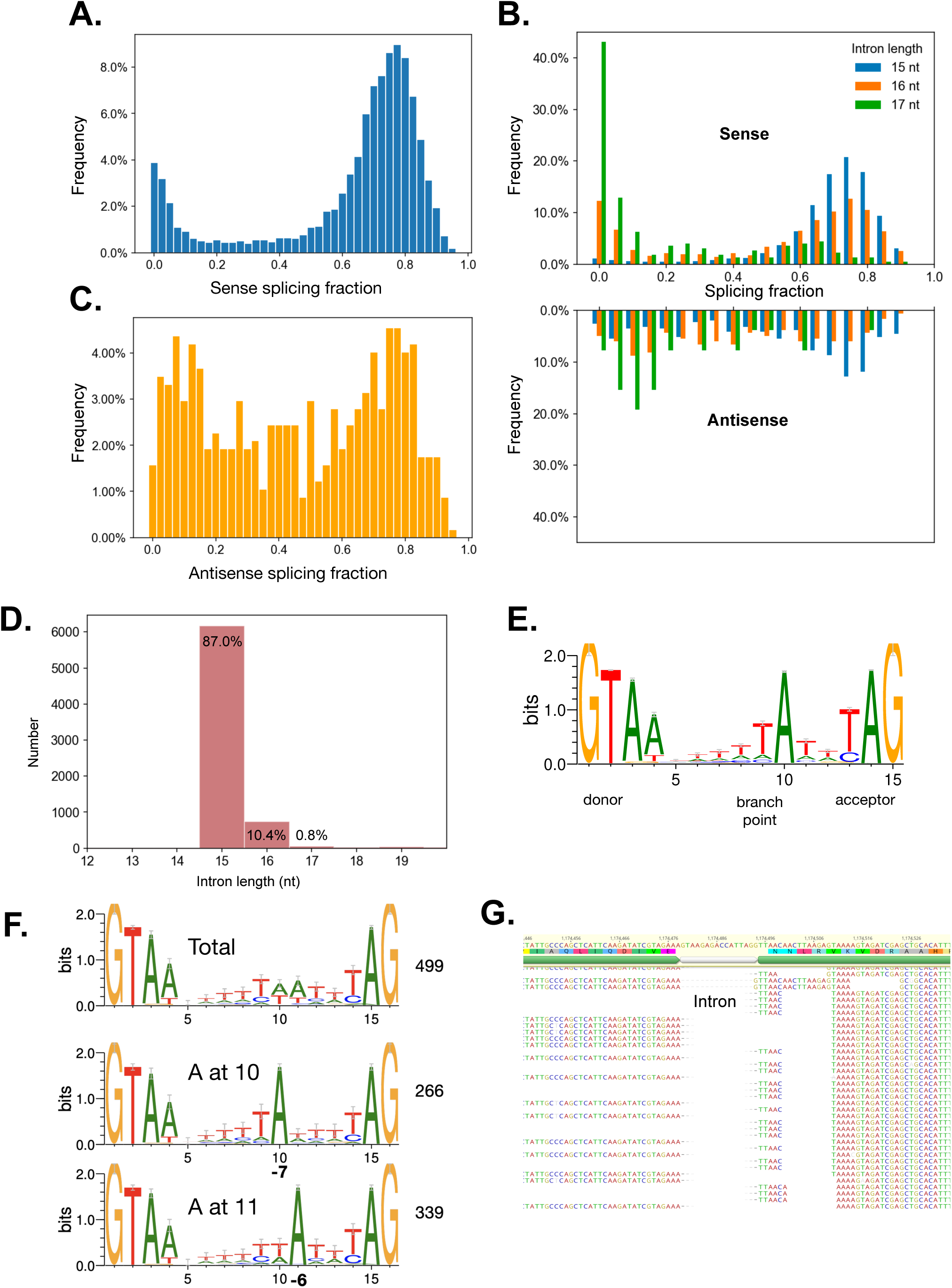
Intron splicing. **A.** Distribution of intron splicing fraction of candidate sense introns in the *B. stoltei* MAC genome. **B**. Distribution of intron splicing fractions of introns according to intron lengths. **C.** Distribution of intron splicing fraction of candidate antisense introns. **D.** Distribution of intron lengths from predicted genes. **E.** Sequence logos for 15 bp introns (splicing frequency > 0.5). **F.** Sequence logos for all predicted 16 nt introns, and 16 nt introns with “A” at either position -7 or -6 (counting from the 3’ end). The number of introns underlying the logos are indicated to the right. **G.** Distribution of intron splicing fractions of introns according to intron lengths. **H.** Sample of RNA-seq reads mapped to a GT-GG intron from gene BSTOLATCC_MAC21551 (Contig_57.g761). Translation in alternative reading frames downstream of the predicted intron leads to premature stop codons soon after the intron.

**Figure S5.**
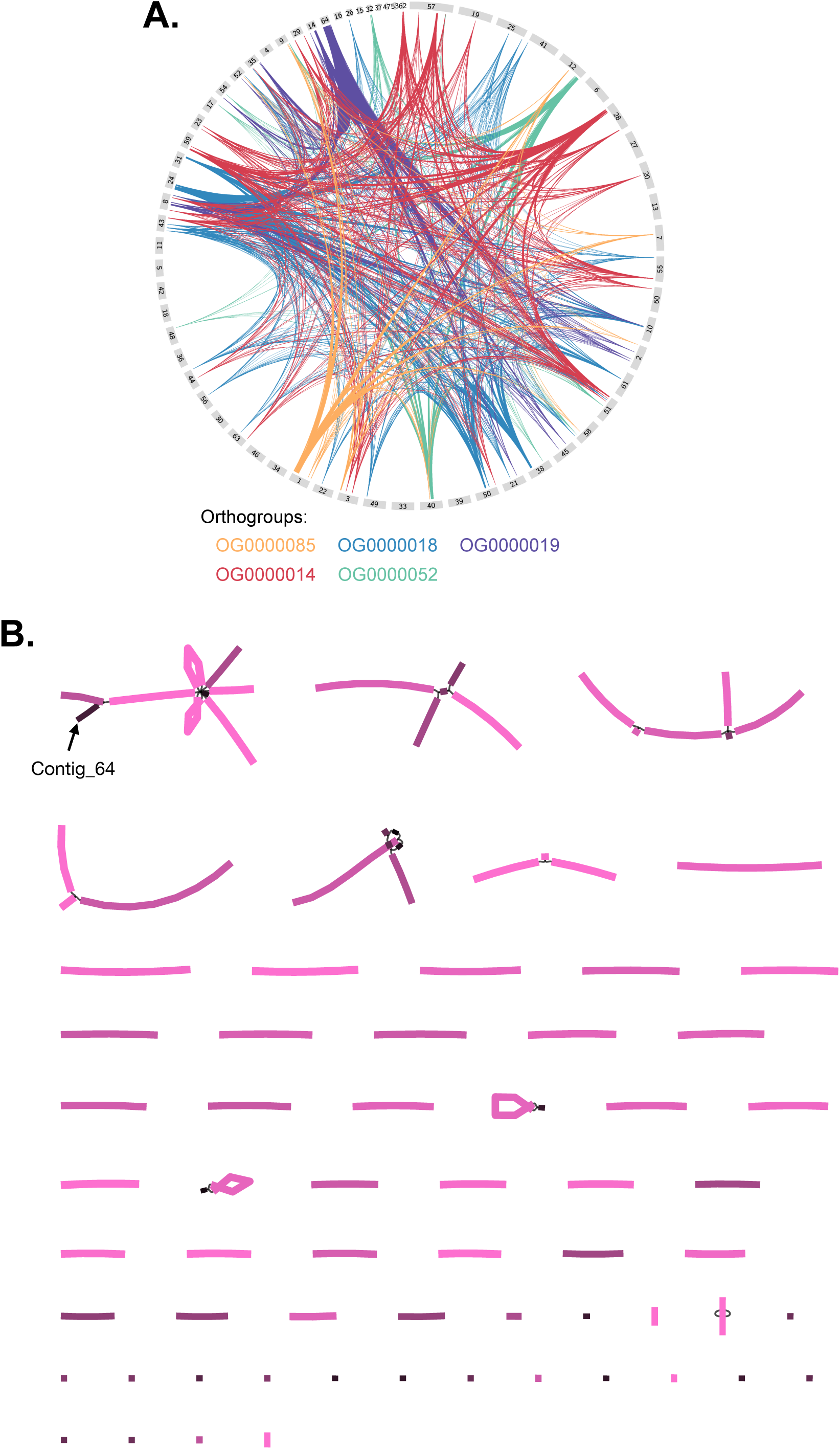
*B. stoltei* ATCC30299 MAC genome orthogroups and assembly graph. **A.** Clustered orthogroups (Data S2) in the *B. stoltei* MAC genome. **B.** Bandage (Wick et al., 2015) representation of Flye 2.8.1 assembly graph. Edges corresponding to contigs are colored by coverage (brightest pink = 160×, black=0×).

**Figure S6.**
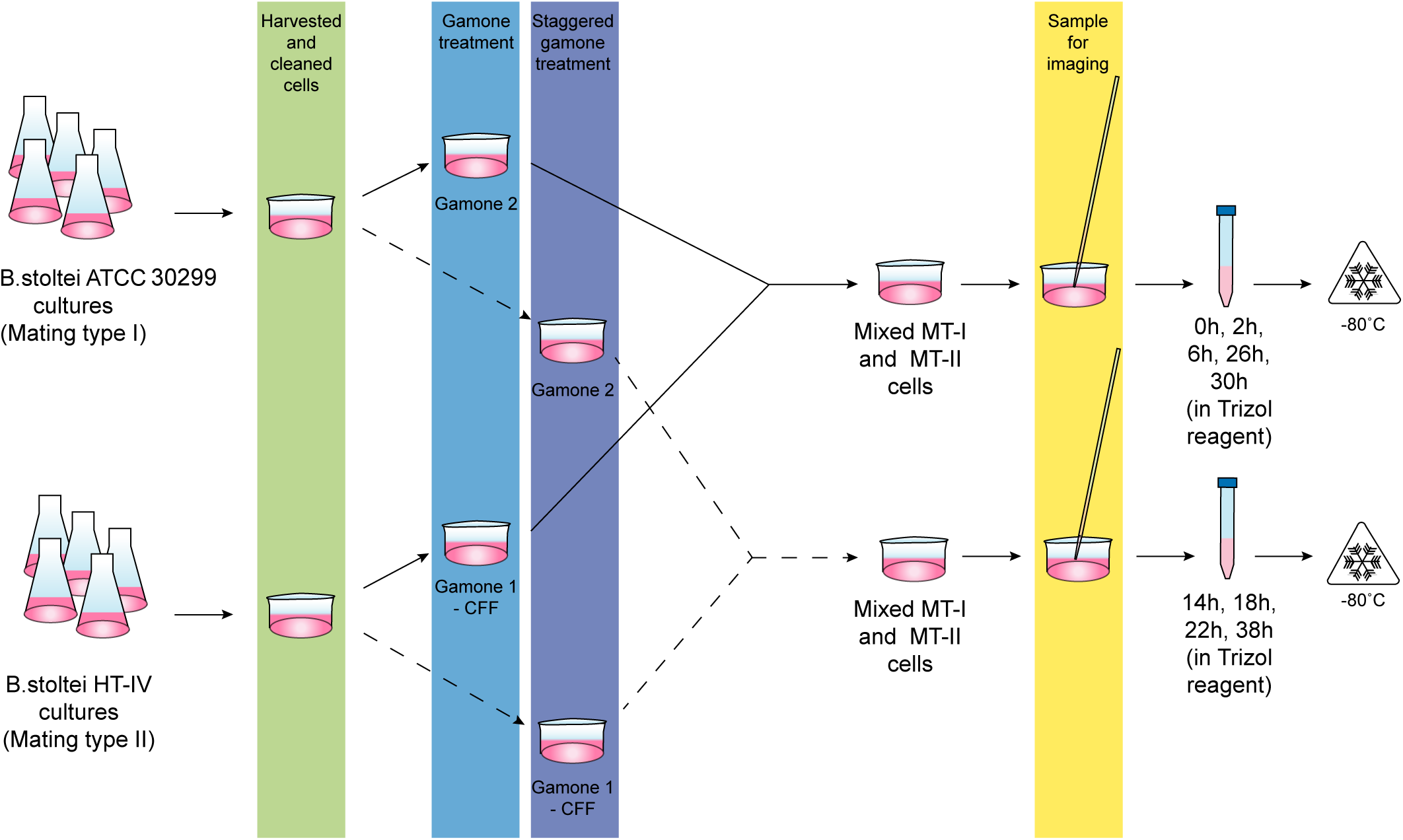
Experimental approach for conjugation RNA-seq time series. Complementary mating type strains of *Blepharisma stoltei* were harvested and cleaned by starving overnight. The cleaned cultures were treated in a time-staggered format, with gamones of the complementary mating type, where gamone 2 was a solution of the synthetic gamone 2 calcium salt and gamone 1 was provided as the cell-free fluid (CFF) harvested from mating-type I cells. Two sets of time-staggered gamone-treated cultures were used for the time series. Set I, indicated by the solid line, was mixed and used to observe and collect samples at 0 hours, 2 hours, 6 hours, 26 hours and 30 hours after mixing. Set II, indicated by the dashed lines, was mixed and used to observe and collect samples at 14 hours, 18 hours, 22 hours and 38 hours after mixing. Test tubes indicate Trizol samples prepared for RNA-extraction which were stored at -80 °C before processing. Cells collected for imaging were obtained shortly before the remainder were transferred into Trizol.

**Figure S7.**
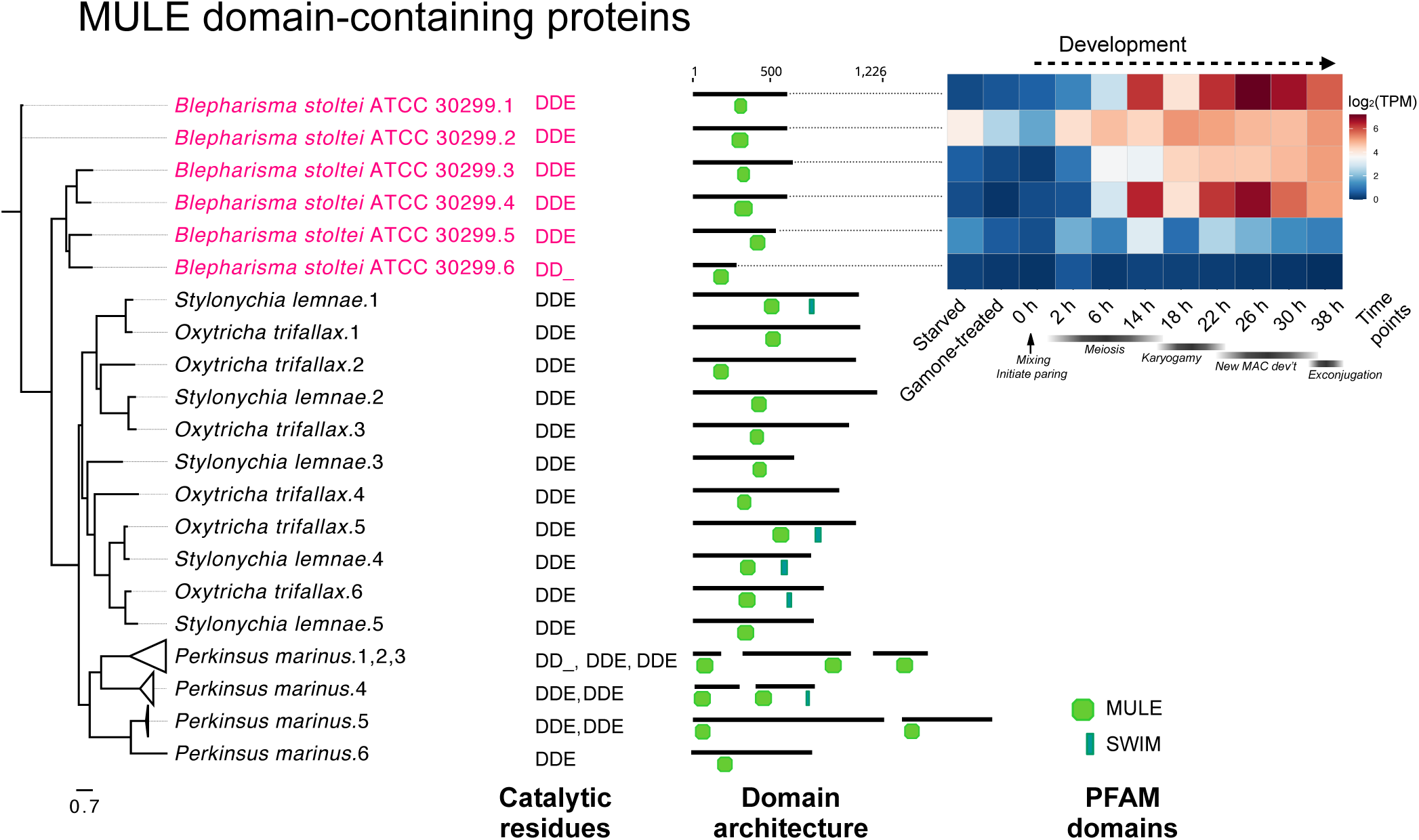
MULE domain transposases in *Blepharisma*. MULE domain phylogeny with PFAM domain architecture and gene expression heatmap for *Blepharisma*.

**Figure S8.**
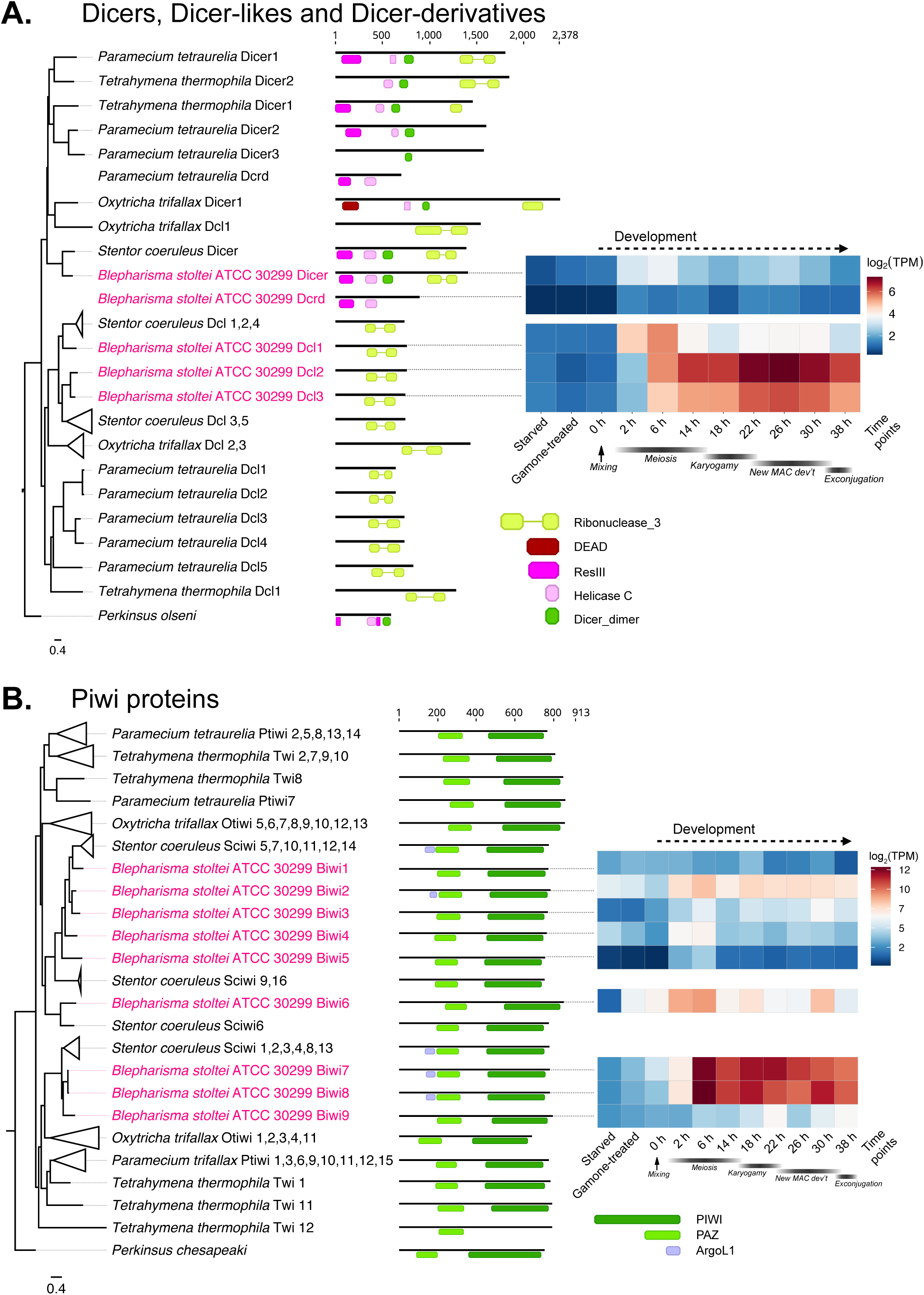
Small RNA-related proteins in *Blepharisma*. **A.** ResIII, Helicase_c and Ribonuclease_3 domain phylogeny with PFAM domain architecture and gene expression heatmap for *Blepharisma*. **B.** PIWI domain phylogeny with PFAM domain architecture and gene expression heatmap for *B. stoltei*.

**Figure S9.**
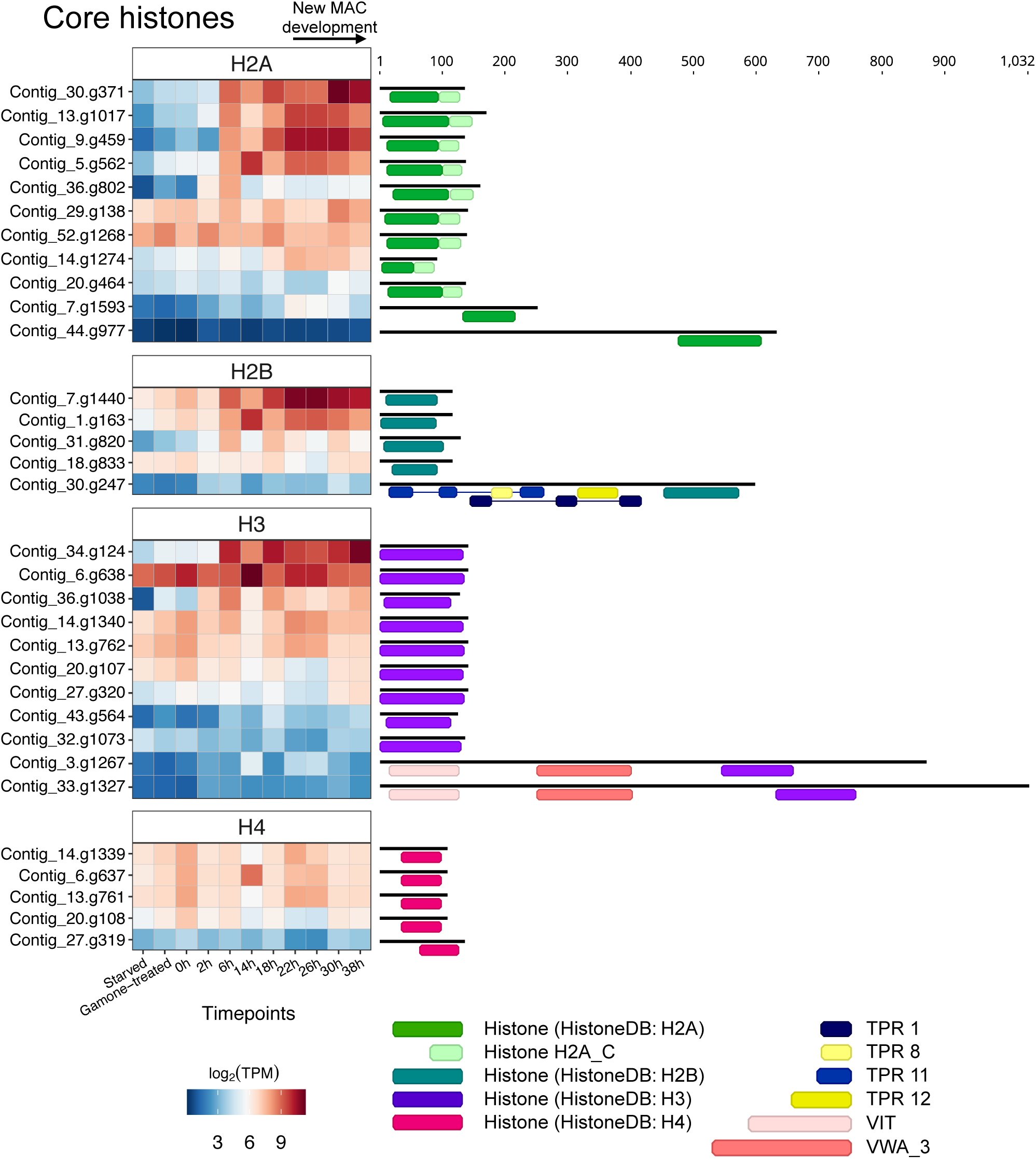
Histones and histone-domain-containing proteins in *Blepharisma*. Gene expression heatmaps are shown as in previous figures, are clustered according to major histone type as classified using HistoneDB domain models. Domains from PFAM and HistoneDB are shown to the right.

