## Supplementary_Information for "Genome editing excisase origins illuminated by somatic genome of *Blepharisma*"

### Supplemental information

#### Additional assembly considerations and inspection

Compared to assemblies of independent, replicate libraries with 70× or 76× coverage there was a modest improvement (e.g. from 86 and 89 contigs to 74) in assembly contiguity for the assemblies produced at 145× coverage (Table S1). With increasing sequence depth, reads of micronuclear origin could conceivably start linking MAC chromosomes and extending their ends, even though we depleted MIC DNA by sucrose gradient centrifugation of nuclei.

We chose to conservatively trim back the ends of most of the contigs in the assembly, and break apart a few contigs at internal sites. This was done either where the coverage locally decreased or increased and where there were extensive differences between the contig sequence and mapped reads. Some of these sequences could represent an intermediate state between fully retained and fully eliminated DNA in MACs. 1.3 Mb of such uncertain sequences (termed “cruft” in genome assembly terminology) were removed from the main assembly. No tRNA genes were predicted in the cruft sequences, nor did their removal reduce the BUSCO score for completeness of the MAC genome, which is comparable to or better than that of other ciliates (Figure S1). BUSCO analyses (Figure S1) also showed that gene duplication in *Blepharisma*, though common, is lower than in *Paramecium tetraurelia* and *Stentor coeruleus*.

#### Alternative telomere addition sites (ATASs)

Alternative telomere addition sites in the MAC genome tend to be intergenic in model ciliates like *Oxytricha trifallax* [(Swart et al., 2013)](https://sciwheel.com/work/citation?ids=2065272&pre=&suf=&sa=0). In *Blepharisma*, we found more intergenic ATASs (28309) than intragenic ones (18396). As intergenic regions only make up 10.1 Mb of the assembly, the intergenic frequency of ATASs is about five-fold higher (2.81 per 1 kb) than intragenic frequency (0.562 per 1 kb). The presence of intragenic ATASs raises the question how the cell tolerates or deals with mRNAs encoding partial proteins transcribed from 3’ truncated genes. Since the sequence data was from a clonal population, it is not possible to tell how much ATAS variability there is within individual cells. However, it is conceivable that their positional variation in single cells reflects that of the population. In this case, together with redundancy from massive DNA amplification there would likely be sufficient intact copies of every gene.

Beyond the first 2-5 bp corresponding to the junction sequences, the average base composition on the chromosome flanking ATAS junctions shows an asymmetrical bias (Figure S2D). From position +6 onwards there is an enrichment of T to about 40% and A to 35-39%, compared to the genome-wide frequencies of 33% each. At position +19 to +23, there is a slight decrease in T to 37-39%. AT values gradually decline back to about 35% each by position +150. Correspondingly, G and C are depleted downstream of ATAS junctions, dropping to a minimum of 8.6% and 11% respectively around position +37, compared to the genome-wide average of 17% each. AT enrichment and GC depletion upstream of ATAS junctions are less pronounced.

If breakage and chromosome healing were random, we would not expect such an asymmetry. This suggests that there is a nucleotide bias, whether in the initiation of breaks, telomere addition, or in the processing of breaks before telomere addition. However, we have not yet identified any conserved motif like the 15 bp chromosome breakage site (CBS) in *Tetrahymena* [(Yao et al., 1990)](https://sciwheel.com/work/citation?ids=10576221&pre=&suf=&sa=0) nor a short 10-bp sequence periodicity in base composition like in *Oxytricha trifallax* [(Cavalcanti et al., 2004)](https://sciwheel.com/work/citation?ids=10575970&pre=&suf=&sa=0). Therefore, telomere addition in *B. stoltei* appears to involve base-pairing of short segments of about 2 bp between the telomere and chromosome, with a bias centered on the “CT” in the telomere unit, and an asymmetrical preference for AT-rich sequences on the chromosomal side of the junction.

The position of an ATAS junction is potentially ambiguous because the last adjoining telomere repeat can potentially be extended into the chromosomal sequence, if the chromosomal sequence at the junction contains a partial match to the telomere (Figure S2C). The junction position that maximizes the length of the telomere sequence on a read has been termed as the “first identifiable breakpoint”, and that which maximizes the chromosomal sequence as the “last identifiable breakpoint” [(Putnam et al., 2004)](https://sciwheel.com/work/citation?ids=1682447&pre=&suf=&sa=0). The overlapping sequence, which could either be telomeric or chromosomal, is termed the “junction sequence”.

Most ATAS junctions in *B. stoltei* have an overlapping junction sequence, on average 2-3 bp long (Figure S2G). This can also be observed when separate sequence logos are drawn for each of the possible telomere repeat permutations observed at the ATAS junction (Figure S2E). Such a short overlap of a few base pairs between telomere repeat and chromosome sequence is similar to what has been observed in other organisms, such as 3-5 bp in yeast [(Putnam et al., 2004)](https://sciwheel.com/work/citation?ids=1682447&pre=&suf=&sa=0) and 2-4 bp in humans [(Morin, 1991)](https://sciwheel.com/work/citation?ids=10575969&pre=&suf=&sa=0). This is in contrast to *Tetrahymena* where telomeres are often added to sites that have no homology to the telomere sequence [(Wang and Blackburn, 1997)](https://sciwheel.com/work/citation?ids=1583815&pre=&suf=&sa=0).

We hypothesized that the location of ATAS junctions in the genome might be randomly distributed and simply reflect the baseline sequence composition of the genome and/or the telomeres. To test this, we counted the frequency of 2-mers in the MAC genome (excluding telomeric regions) and in the telomere repeats, and compared them to the 2-mer frequencies observed at ATAS junctions (2 bp on chromosomal side of last identifiable breakpoints, Figure S2F). Sequence composition of the telomeres does have a strong influence, as 2-mers that are not represented in the telomeres (AT, GC, CG, GA) are poorly represented at ATAS junctions even though they may be frequent in the genome, e.g. GA, 12.0% in genome vs. 0.36% at ATAS; AT, 10.4% vs. 1.7%. However, 2-mer frequencies at ATAS junctions do not match frequencies in the telomeres closely either. For example, the 2-mer AG is about twice as frequent at ATAS junctions as compared to telomeres, and as compared to the genome generally. Instead, the telomere permutations at ATAS junctions are not uniformly distributed; the permutation CTAACACC is the most common, followed by its adjacent permutations TAACACCC and AACACCCT (using last identifiable breakpoints, Figure S2H). These would account for the three most common 2-mers at ATAS junctions: AG (canonical form of CT), AA, and TA.

#### Telomere-binding protein paralogs

Despite the abundance of *Blepharisma* MAC genome telomeres, we did not detect a typical ncRNA gene corresponding to the telomerase RNA component (TERC) of the ribozyme responsible for telomere synthesis in the MAC genome. We suspect this is due to ncRNAs presenting a far greater challenge to detect than protein-coding genes and the presence of highly divergent ncRNA with insufficient similarity to the handful of taxonomically-restricted TERCs identified in oligohymenophorean and spirotrich ciliates and other eukaryotes so far.

Other than the components of telomerase, ciliate were among the first organisms where telomere-binding proteins were characterized [(Gray et al., 1991)](https://sciwheel.com/work/citation?ids=1552383&pre=&suf=&sa=0). Telomere-binding protein paralogs with distinctive patterns of gene expression during development are present in some ciliate species [(Cranert et al., 2014; Swart et al., 2013)](https://sciwheel.com/work/citation?ids=2065272,6345192&pre=&pre=&suf=&suf=&sa=0,0). *Tetrahymena* *thermophila* has two telomere-binding protein paralogs POT1 and POT2 [(Cranert et al., 2014)](https://sciwheel.com/work/citation?ids=6345192&pre=&suf=&sa=0). POT2 is upregulated during conjugation, accumulates in developing new macronuclei, and binds to chromosome breakage sites rather than telomeres [(Cranert et al., 2014)](https://sciwheel.com/work/citation?ids=6345192&pre=&suf=&sa=0). *Blepharisma stoltei* has five POT1 paralogs POT1.1-POT1.5 (Figure S3C). One *B. stoltei* POT1 paralog expressed at low levels in starved (0 h) cells, POT1.4, is sharply upregulated during development, peaking when new macronuclei are forming (22 h) (Figure S3C).

Since we were unable to identify a specific chromosome breakage signal like that of *Tetrahymena*, a future avenue to search for such a signal would be to assess the DNA-binding preferences of POT1.4 and the other *Blepharisma* POT1 paralogs. In any event, since this is one of the most highly upregulated genes in the 22-26 h time range compared to vegetative (0 h, and gamone treated cells; see “Results”, “Features of gene expression during new MAC development”), future investigation of its developmental role is warranted.

#### Tiny spliceosomal introns

Like *Stentor* [(Slabodnick et al., 2017)](https://sciwheel.com/work/citation?ids=5381445&pre=&suf=&sa=0), most (82%) *Blepharisma* genes have no introns. In line with genome compactness, during our inspections we also observed numerous overlapping poly(A)-tailed RNA-seq reads on opposite strands derived from convergently transcribed gene pairs. The correlation of the lengths of different noncoding region classes (intergenic regions, introns and UTRs) can be explained by them being subject to common, neutral evolutionary processes [(Lynch, 2006)](https://sciwheel.com/work/citation?ids=402819&pre=&suf=&sa=0).

*Blepharisma* introns are mostly (97%) 15 or 16 nucleotides (nt) long, like those of *Stentor* (Figure S4D). Though intron reduction (7389 introns predicted in the reference *B. stoltei* MAC genome, i.e., 0.29 introns per gene) is not as extreme as some other microbial eukaryotes, like *Giardia lamblia* [(Roy et al., 2012)](https://sciwheel.com/work/citation?ids=11964032&pre=&suf=&sa=0), where almost all have been lost, both *Blepharisma* and *Stentor* have much fewer introns relative to other ciliates (e.g., intron densities of 1.6, 2.3 and 4.8 introns per gene in *Paramecium*, *Oxytricha* and *Tetrahymena*, respectively [(Bondarenko and Gelfand, 2016)](https://sciwheel.com/work/citation?ids=5381339&pre=&suf=&sa=0)) and to the putative, relatively intron-rich eukaryotic common ancestor [(Csuros et al., 2011)](https://sciwheel.com/work/citation?ids=1210161&pre=&suf=&sa=0), along with their extreme length reduction.

*Blepharisma* 15 nt introns possess a characteristic branch-point “A”, as would be expected in classical models of lariat formation during mRNA splicing (Figure S4C). 16 nt introns almost invariably have an “A” at either 10 or 11 nt downstream of the donor site (i.e., only one of 499 does not, but has “A” at 9 nt), although this is not obvious in the consensus sequence logo because the position is variable (Figure S4D). Similarly, 17 nt introns all possess “A” at 10-12 nt downstream of the donor site. Only a few intron bases, 5-8 and 12, of *Blepharisma*’s 15 nt introns are relatively unconstrained (Figure S4C). This leaves little room for the presence of any additional regulatory elements in the mRNA or underlying DNA.

In the final gene predictions, just over 1% of predicted *Blepharisma* introns lack canonical GT-AG boundaries (62 out of 4670 introns). Just under half of these (30) are 15 or 16 bp long and predominantly appear to represent true spliceosomal introns. The boundaries of two predicted introns with CT-AC boundaries (14 and 15 nt in length) resulted from misalignment of nucleotides in the mapped spliced reads at conventional GT-AG junctions. We found no evidence of minor spliceosomal RNAs (U11, U12, U4atac, and U6atac) using Infernal searches (E-value < 10). Thus, *Blepharisma* appears to lack a minor spliceosome and minor spliceosomal introns. As far as we are aware no minor spliceosomal introns have been reported in any ciliates. Loss of minor spliceosomal machinery and introns, relative to the eukaryotic common ancestor, may be relatively common in alveolates including ciliates [(Russell et al., 2006)](https://sciwheel.com/work/citation?ids=456205&pre=&suf=&sa=0).

The most common 5’ boundaries for *Blepharisma* introns that possess a 3’-AG but lack 5’-GT are 5’-GC or 5’-GG (the latter are most often 5’-GGT; Table S4). Introns that possess a 5’-GT but lack 3’-AG typically have 3’-GG boundaries (most often 3’-AGG; Table S4). Visual inspection of the mapped RNA-seq data to the non-canonical *Blepharisma* introns and predicted coding sequences suggests that the GC-AG, GT-GG and GG-AG introns are correct, i.e., lead to prediction of complete coding sequences downstream of their locations. Lower frequency alternative splicing may occur in some cases (e.g. Figure S4G), but these generate prematurely terminated coding sequences.

GC-AG introns are the most common alternative major spliceosomal introns in multicellular organisms [(Sheth et al., 2006)](https://sciwheel.com/work/citation?ids=5407365&pre=&suf=&sa=0). In *Blepharisma* such introns are most frequently 15 bp long. In contrast to GC-AG introns and conventional *Blepharisma* GT-AG introns, GG-AG and GT-GG (or GGT-AG and GT-AGG) introns are 16 bp or longer (Table S4). This suggests most of these introns evolved from conventional 15 bp GT-AG introns. It is possible that splicing of the shorter internal GT-AG introns, instead of their longer non-canonical forms that give rise to full-length coding sequences, leads to NMD of some mRNAs, since these invariably have a premature in-frame stop codon downstream of the intron. Thus RNA-seq may underestimate the amount of splicing of the shorter forms.

#### Overcoming challenges in gene prediction due to tiny introns

As reported in *Stentor*, splicing frequency decreases as intron length increases in *Blepharisma* (Figure S4B). This trend is also evident in antisense introns, though weaker and more noisy due to their lower abundance (Figure S4B). Since antisense intron splicing would be free from selective constraints imposed by protein-coding sequence translation, we suggest that the intron length distribution primarily reflects the splicing length preferences of the spliceosome. The decreased efficiency of splicing of introns longer than 15 nt, and evident inability to splice introns shorter than this, means that most intron indels may be deleterious. We therefore suggest that, like its IESs which are skewed towards shorter lengths (Seah et al. 2022), *Blepharisma*’s introns can largely be thought of as parasitic elements which bear significant potential costs. This would also be consistent with the absence of introns in most heterotrich genes, and a pronounced decrease in intron density relative to model ciliates such as *Paramecium*, *Tetrahymena* and *Oxytricha* [(Slabodnick et al., 2017)](https://sciwheel.com/work/citation?ids=5381445&pre=&suf=&sa=0).

The tiny introns of *Stentor coeruleus* previously created significant challenges for gene prediction [(Slabodnick et al., 2017)](https://sciwheel.com/work/citation?ids=5381445&pre=&suf=&sa=0) using AUGUSTUS [(Stanke and Waack, 2003)](https://sciwheel.com/work/citation?ids=2757128&pre=&suf=&sa=0). In the *Stentor* study, some predicted genes were observed to be incorrectly joined, and so were split with a custom script. Furthermore, introns of lengths other than 15 or 16 bp were attributed to genome mis-assembly [(Slabodnick et al., 2017)](https://sciwheel.com/work/citation?ids=5381445&pre=&suf=&sa=0). In our study, after adjusting AUGUSTUS parameters for tiny introns as for *Stentor*, and training AUGUSTUS for gene prediction in *Blepharisma* from visual inspection of mapped RNA-seq reads, we saw that most predicted introns longer than 16 bp are incorrect. With the benefit of major technological advances in long read sequencing and considerably increased sequencing depth over the last years, the *Blepharisma* MAC genome assembly is not as prone to misassembly, and contiguity substantially improved compared to that of the draft *Stentor* *coeruleus* assembly. Consequently most of the incorrect introns predicted with AUGUSTUS in *Blepharisma* were errors in gene prediction rather than mis-assembly. Additionally, numerous introns, including some of length 15 or 16 nt, were predicted in regions deeply covered by RNA-seq with no mapped reads evidencing splicing. No matter what changes we attempted to the AUGUSTUS source code in attempts to more accurately predict introns, more were incorrectly predicted than not (e.g. Table S3).

Since we obtained extensive RNA-seq data across a developmental time course which appeared to cover most genes (e.g. Figure 3A), we chose to eliminate incorrect intron predictions, the major source of inaccuracy in *Blepharisma* gene predictions, by directly predicting introns using mapped reads. This approach, Intronarrator, runs AUGUSTUS in “intronless” prediction mode on a version of the genome with introns removed, before replacing the introns in the genes. Visual inspection of the predicted introns on Contig_1, showed there was a marked improvement in intron prediction sensitivity from 0.75 with AUGUSTUS to 0.97 with Intronarrator, while precision improved from 0.42 to 1.00 (Table S3). In general, there is consistency between the locations of the predicted genes and RNA-seq coverage, notably including genes with introns (Figure 3A).

#### Extensive duplications of transmembrane protein genes

A notable extended ~220 kb region encoding 53 genes belonging to a single orthologous group (orthogroup), OG0000085 is present on Contig_1 (Figure 3A). Four additional OG0000085 genes are present at the opposite end of Contig_1, and 24 copies are found on other contigs, often clustered together (Figure S5A). The DNA coverage across this region is lower (74×) than the rest of Contig_1 (185×). Though there is uncertainty in the exact extent, given the sheer volume of reads involved, the assembled sequences certainly correspond to highly repetitive regions of the MAC genome. At the junction between the lower and higher coverage regions more than 30 HiFi reads link the two regions of coverage, and a similar number of telomere-bearing reads are in close proximity. At the junction we also observe at least two potential locations of IESs, corresponding to regions that may be partially IES/partially MDS.

Large clusters of genes from particular orthogroups can be found on additional contigs (Figure S5). In total 551 (2%) of predicted *B. stoltei* genes belong to the orthogroups with the largest clusters per contig. Some of the largest contiguous clusters of genes from these orthogroups are situated at the ends of contigs, suggesting they may have caused assembly breaks beyond them. One contig, split off from other connected components in the assembly graph, predominantly encodes genes from a single orthogroup (contig_64, 43× coverage; Figure S5). Further increases in read length and accuracy may allow assemblers to fully resolve these in future. Curiously, all the orthogroups corresponding to the largest contiguous clusters of genes appear to be transmembrane proteins, or decayed remnants thereof. The nature of these proteins is described in Supplemental Text (“Properties of proteins encoded by extensive duplications”).

Full-length proteins from OG0000085 (81 proteins in total) and OG0000014 (143 proteins in total) both contain a central PFAM “ANF_receptor” domain (PF01094), annotated in the PFAM database with the description “This family includes extracellular ligand binding domains of a wide range of receptors”. Though distantly related, e.g. 32% amino acid identity of the consensus sequences (produced by the majority rule for each orthogroup), the full-length proteins are of similar length and align well, and thus are likely homologs. A clear C-terminal transmembrane domain region comprising seven to nine alpha helices is predicted for presumed full-length versions of proteins from both ortholog groups using TMHMM2 [(Krogh et al., 2001)](https://sciwheel.com/work/citation?ids=178741&pre=&suf=&sa=0). Queries of UniProt revealed that, though widely distributed among eukaryotes, among ciliates only *Stentor coeruleus* also possesses proteins with this domain classified (29 in total). BLAST searches versus the GenBank NR database detect a similar number of matches to *Stentor* *coeruleus* homologs (E-value < 1e-30) but none in any other ciliates. Ortholog groups OG0000018 and OG0000052 also appear to be homologous to one another (31% amino acid identity of the consensus sequences produced by the majority rule for each orthogroup). Full-length proteins from these ortholog groups possess a clear N-terminal transmembrane domain predicted by TMHMM2, composed of seven or more transmembrane helices. We also detected a central Pas domain (PF00989) in a couple of these proteins in InterProScan searches of PFAM. Ortholog group OG0000019 has a seven transmembrane C-terminal domain predicted by TMHMM2, a series of centrally located “Laminin_G_3” (PF13385) domains, and an N-terminal “Malectin” (PF11721) domain in some proteins.

Ciliates encode a moderately large number of protein-coding genes compared to other eukaryotes, often exceeding 25,000. Species like *Paramecium tetraurelia* which have undergone multiple whole genome duplications, may have more than 40,000 genes [(Aury et al., 2006)](https://sciwheel.com/work/citation?ids=149353&pre=&suf=&sa=0). In ciliate species like *Tetrahymena thermophila* (26,258 genes [(Sheng et al., 2020)](https://sciwheel.com/work/citation?ids=10022620&pre=&suf=&sa=0)), with no evidence of whole genome duplications, it has been a question as to why these species are so gene rich [(Eisen et al., 2006)](https://sciwheel.com/work/citation?ids=149350&pre=&suf=&sa=0).

Segmental duplications identified in the human genome are defined as duplications > 1 kb and > 90% sequence identity [(Bailey et al., 2001)](https://sciwheel.com/work/citation?ids=511303&pre=&suf=&sa=0). Little evidence for such duplications was found in the *Tetrahymena* MAC genome [(Eisen et al., 2006)](https://sciwheel.com/work/citation?ids=149350&pre=&suf=&sa=0). Since the divergences of the proteins within the large clustered *Blepharisma* orthogroups are typically moderately high (e.g. < 40% amino acid identity), the duplications that led to them represent older events. Nonetheless, given their extent, it is likely that many of the duplicated genes originated from segmental duplications. Recombination of clusters of some of these genes into other genomic regions may subsequently have spread them elsewhere, and led to gradual erosion of the original locus. The OrthoFinder algorithm is specifically designed to eliminate scoring biases against shorter sequences, a significant advance over older algorithms like OrthoMCL [(Emms and Kelly, 2015)](https://sciwheel.com/work/citation?ids=1098484&pre=&suf=&sa=0). In our inspections of multiple sequence alignments of the largest orthogroups we also detected numerous genes that are clearly related to, but significantly shorter than the typical gene length of each orthogroup, thus likely representing eroding pseudogenes. Though we focused on the largest and most notable clusters of genes from the orthogroups, numerous other genes may also have arisen out of such clustered duplications.

#### Development-specific upregulation of proteins associated with DNA repair and chromatin

A variety of different DNA repair protein genes are strongly upregulated at 26 hours (Table S6; Data S3), among them: a 5’ Apollo exonuclease protein (BSTOLATCC_MAC16643), whose homologs are involved in DNA repair and telomere protection [(Lenain et al., 2006)](https://sciwheel.com/work/citation?ids=1131409&pre=&suf=&sa=0) (a paralog of this gene is constitutively expressed at low levels: BSTOLATCC_MAC3725; 58.9% pairwise amino acid identity); STAG1/2 (BSTOLATCC_MAC22820) and Rad21 (BSTOLATCC_MAC1548) homologs, both cohesin complex components, and a Rad50 homolog (BSTOLATCC_MAC2159), all proteins involved in DNA double-strand break repair [(Mondal et al., 2019; Rojowska et al., 2014)](https://sciwheel.com/work/citation?ids=6790429,1262197&pre=&pre=&suf=&suf=&sa=0,0); a homolog of MUS81 (BSTOLATCC_MAC21072) a protein involved in meiotic double-strand break repair in *Tetrahymena* [(Lukaszewicz et al., 2013)](https://sciwheel.com/work/citation?ids=766860&pre=&suf=&sa=0); a homolog of PARP2 (Poly(ADP-ribose) polymerase-2) (BSTOLATCC_MAC1058), a protein involved in DNA single-strand nick repair [(Riccio et al., 2016)](https://sciwheel.com/work/citation?ids=8646007&pre=&suf=&sa=0); a homolog (BSTOLATCC_MAC1470) of the DNA clamp, PCNA, which is involved in DNA repair associated with DNA polymerases delta and epsilon [(Shivji et al., 1992; Travali et al., 1989)](https://sciwheel.com/work/citation?ids=2085540,2086223&pre=&pre=&suf=&suf=&sa=0,0); two homologs (BSTOLATCC_MAC23155 and BSTOLATCC_MAC23646) of exodeoxyribonuclease III, a protein involved in abasic DNA base repair [(Mol et al., 1995)](https://sciwheel.com/work/citation?ids=449666&pre=&suf=&sa=0).

A dozen chromatin-related proteins are among the top 100 most strongly upregulated proteins at 26 hours (Table S6). These include a homolog (BSTOLATCC_MAC17684) of ISWI, a core ATPase remodeler present in a range of different chromatin remodelling protein complexes in eukaryotes [(Corona et al., 1999)](https://sciwheel.com/work/citation?ids=2082703&pre=&suf=&sa=0). In *Paramecium* *tetraurelia* the strongest developmentally upregulated ISWI homolog plays a critical role in nucleosome positioning in new MACs during genome editing (Singh, et al., submitted). A few histone/histone-related proteins and HMG boxes are also strongly upregulated. Two JmjC (Jumonji C) domain-containing proteins are also highly upregulated (BSTOLATCC_MAC23590 and BSTOLATCC_MAC5044). Proteins with this domain are histone lysine demethylases in other eukaryotes [(Klose et al., 2006)](https://sciwheel.com/work/citation?ids=222796&pre=&suf=&sa=0). BSTOLATCC_MAC23590 is likely to be orthologous to JMJ1 of *Tetrahymena* *thermophila* (TTHERM_00185640): they are reciprocal best BLASTP hits (with next best hit e-values many orders of magnitude higher), the JmjC domain occurs in a similar relative location in the two proteins, and their lengths are similar (1082 aa and 1198 aa). JMJ1 is highly upregulated during *T.* *thermophila* conjugation, first localizing in old MACs and later in the new MACs [(Chung and Yao, 2012)](https://sciwheel.com/work/citation?ids=6344895&pre=&suf=&sa=0). This protein is required for H3K27me3 demethylation later in conjugation, where it is proposed to influence gene expression, including those expressed later and involved in genome editing processes, rather than heterochromatin associated with *Tetrahymena* IES excision per se [(Chung and Yao, 2012)](https://sciwheel.com/work/citation?ids=6344895&pre=&suf=&sa=0).

#### Development-specific upregulation of proteins associated with initiation of transcription and translation

While overarching coordination of gene regulation and protein translation are expected during ciliate development, it is not evident how this might be achieved. Among the most strongly upregulated genes at 26 hours are homologs of proteins involved in initiation of either transcription or translation, notably an eIF4E translation initiation factor homolog (BSTOLATCC_MAC5291) and a TATA-binding protein (SPT15; BSTOLATCC_MAC11469; Table S6). In other eukaryotes eIF4E proteins bind to m7G 5’ mRNA caps permitting protein translation [(Sonenberg and Hinnebusch, 2009)](https://sciwheel.com/work/citation?ids=181786&pre=&suf=&sa=0). *B. stoltei* has ten homologs of these proteins, nine of which are moderately stably expressed throughout the RNA-seq conditions examined (Data S3; workbook “eIF4e homologs”). eIF4E paralogs are also abundant in *S. coeruleus*, with thirteen homologs found by BLASTP. One of the *B. stoltei* eIF4E paralogs is more highly expressed than the rest (BSTOLATCC_MAC25346), however this is still roughly an order of magnitude less than the development-specific paralog in all times after 2 hours post cell mixing. The pronounced upregulation of a homolog of eIF4e would be consistent with the massive amount of protein translation necessary during development. We therefore propose that translation initiation plays a critical regulatory role in protein synthesis during *Blepharisma* development, all the way through genome editing.

Regarding transcription regulation, *B. stoltei* has a constitutively expressed TATA-binding protein (BSTOLATCC_MAC16553) which is 64.8% identical (at the amino acid level) to the developmentally upregulated paralog (BSTOLATCC_MAC11469). *Tetrahymena thermophila* also appears to have two TATA-binding protein homologs annotated (TBP1 and TBP2; 30.5% pairwise amino acid identity) both of which are modestly upregulated during development (<http://tfgd.ihb.ac.cn/search/detail/gene/TTHERM_00575350> and <http://tfgd.ihb.ac.cn/search/detail/gene/TTHERM_00082170>). In *B. stoltei* we speculate that the two TATA-binding proteins may recognize distinct TATA box motifs, and thus transcription of a large, development-specific subset of proteins might be controlled by a master regulator. A homolog of transcription initiation factor TFIID subunit 1 (TAF), the largest core component of the transcription initiation complex [(Wang et al., 2014)](https://sciwheel.com/work/citation?ids=11126123&pre=&suf=&sa=0) that interacts with TATA-binding proteins [(Dynlacht et al., 1991)](https://sciwheel.com/work/citation?ids=4996385&pre=&suf=&sa=0) is encoded by the sixth most strongly upregulated gene (388×) at 26 hours (BSTOLATCC_MAC12987). The only other TAF homolog we detected (BSTOLATCC_MAC10371) is more weakly upregulated (12×) at 26 hours.

In *B. stoltei* an additional homolog of a protein involved in transcription elongation (SPT5; BSTOLATCC_MAC7803) is among the most highly upregulated genes at 26 hours, and also has a constitutively expressed paralog (BSTOLATCC_MAC18233 78.4% pairwise amino acid identity). *Paramecium tetraurelia* and *Oxytricha trifallax* both have SPT5 paralogs that appear to have been generated in separate duplication events [(Gruchota et al., 2017)](https://sciwheel.com/work/citation?ids=3286967&pre=&suf=&sa=0), and our phylogenies suggest the *B. stoltei* paralogs duplicated independently of these two species. In *P. tetraurelia*, one of the two paralogs is specific to meiotic micronuclei, and has an expression profile that peaks earlier during development prior to meiosis and declines during new MAC formation [(Gruchota et al., 2017)](https://sciwheel.com/work/citation?ids=3286967&pre=&suf=&sa=0). In *Oxytricha* one SPT5 paralog (SPT5a) is constitutively expressed, whereas the other (SPT5b) peaks during meiosis [(Neeb et al., 2017)](https://sciwheel.com/work/citation?ids=6873261&pre=&suf=&sa=0). In *Tetrahymena* the single SPT5 gene is strongly upregulated during development, peaking during meiosis (<http://tfgd.ihb.ac.cn/search/detail/gene/TTHERM_00028580>).

#### Homologs of small RNA-related proteins involved in ciliate genome editing

Development-specific proteins responsible for small RNA (sRNA) generation and transport play an important role in ciliate genome editing [(Chalker et al., 2013)](https://sciwheel.com/work/citation?ids=5616076&pre=&suf=&sa=0). In ciliates such as *Paramecium* and *Tetrahymena* shorter Dicer-like proteins (Dcls) are distinguished from longer Dicer proteins (Dcrs) which possess additional N-terminal domains and produce small RNAs, notably siRNAs, involved in gene regulation [(Sandoval et al., 2014)](https://sciwheel.com/work/citation?ids=613403&pre=&suf=&sa=0). In the scanning model of MAC development in *Tetrahymena* and *Paramecium*, Dcls cooperate with Piwi proteins, converting long double-stranded RNA transcripts produced in the maternal MIC into “scan RNAs” (scnRNAs) [(Lepère et al., 2009; Mochizuki and Gorovsky, 2005; Mochizuki et al., 2002; Noto and Mochizuki, 2018; Sandoval et al., 2014; Schoeberl et al., 2012)](https://sciwheel.com/work/citation?ids=1181530,1181534,613403,4034485,2310080,6344761&pre=&pre=&pre=&pre=&pre=&pre=&suf=&suf=&suf=&suf=&suf=&suf=&sa=0,0,0,0,0,0). Piwi-bound scnRNAs are transported to the maternal MAC where a subtractive process takes place, leaving only scnRNAs complementary to the MIC-limited genome. The remaining scnRNAs are transported to the new, developing MAC, where they target MIC-limited regions for excision [(Lepère et al., 2009; Mochizuki and Gorovsky, 2005; Mochizuki et al., 2002; Noto and Mochizuki, 2018; Sandoval et al., 2014; Schoeberl et al., 2012)](https://sciwheel.com/work/citation?ids=1181530,1181534,613403,4034485,2310080,6344761&pre=&pre=&pre=&pre=&pre=&pre=&suf=&suf=&suf=&suf=&suf=&suf=&sa=0,0,0,0,0,0).

We found putative Dicer, Dicer-like and Piwi proteins encoded by the *B. stoltei* MAC genome (Figure S8)*.* The single *B. stoltei* Dicer (Dcr) protein has the characteristic N-terminal Dicer domains followed by a pair of RNase III domains (PFAM domain Ribonuclease_3; PF00636) whereas RNase III domains alone were detected in three Dicer-like proteins (Dcl1-3). Dcl1 expression is upregulated shortly after conjugation begins and before meiosis begins; Dcl2 and Dcl3 are upregulated from meiosis onwards, peaking during anlagen formation. In *Paramecium* two Dcl’s are coexpressed and cooperate to produce scnRNAs [(Sandoval et al., 2014)](https://sciwheel.com/work/citation?ids=613403&pre=&suf=&sa=0), and so we predict that, as for *Paramecium*, *Blepharisma* Dcl2 and Dcl3 may cooperate.

*B. stoltei* also appears to have an additional truncated Dcr homolog (881 aa), Dicer-derived protein (Dcrd), lacking the RNase III domain portion (BSTOLATCC_MAC8391) of the complete Dicer (Figure S8A). A short protein (690 aa) with a similar domain structure is found in *Paramecium* *tetraurelia* (Genbank accession: XP_001459306.1), and, as judged from gene expression data in ParameciumDB [(Arnaiz et al., 2020)](https://sciwheel.com/work/citation?ids=10253652&pre=&suf=&sa=0), is substantially upregulated during new MAC development. The observation of these proteins suggests that it might be possible for Dicer helicase and cleavage activities to be encoded on separate molecules. The splitting of the helicase and RNAse domains is the converse of the common eukaryotic origin of Dicer from separate archaeal (helicase) and bacterial (RNase) domains [(Shabalina and Koonin, 2008)](https://sciwheel.com/work/citation?ids=1240975&pre=&suf=&sa=0). It is also conceivable, once they have split, that alternative helicases may substitute the original ones of helicase-less Dicer-like proteins.

In ciliates some Piwi proteins play a role in gene regulation in vegetative cells [(Götz et al., 2016)](https://sciwheel.com/work/citation?ids=6951533&pre=&suf=&sa=0) while others are involved in genome editing [(Bouhouche et al., 2011; Fang et al., 2012; Mochizuki et al., 2002)](https://sciwheel.com/work/citation?ids=4034482,1627833,1181530&pre=&pre=&pre=&suf=&suf=&suf=&sa=0,0,0). In *Stylonychia lemnae*, the massive upregulation of a Piwi homolog involved in genome editing allowed it to be identified by subtractive hybridization of RNA [(Fetzer et al., 2002)](https://sciwheel.com/work/citation?ids=6344913&pre=&suf=&sa=0). The ortholog of this gene in *Stylonychia* *lemnae*’s close relative, *Oxytricha* *trifallax*, is also one of the most highly transcribed and upregulated genes [(Fang et al., 2012)](https://sciwheel.com/work/citation?ids=1627833&pre=&suf=&sa=0). We found nine proteins with Piwi and PAZ domains (five of which also have ArgoL domains) in the *B. stoltei* ATCC 30299 MAC genome. Two closely related *Blepharisma* Piwi paralogs are highly upregulated during meiosis and throughout subsequent development (Figure S8B). These two genes are both among the most highly expressed genes at 26 h (12 and 154) while the new MAC is forming.

In *Tetrahymena* and *Paramecium* massive production of scnRNAs, using the Dcls and highly upregulated Piwis, initiates from meiotic nuclei. We observe a similar pattern of massive production of development-specific sRNAs during development, whose detailed analysis will be reported in conjunction with the draft *B. stoltei* ATCC 30299 MIC genome (Seah, et. al, 2022). Since *Blepharisma* species are distantly related to other ciliates whose sRNAs have been characterized, this suggests that an ancient, development-specific sRNA gene expression program may have been established in the ciliate common ancestor.

#### Development-specific histone variant upregulation

Access to DNA in eukaryotes is mediated by nucleosomes and nucleosome regulation plays a central role in DNA replication, repair, transcription and recombination [(Zentner and Henikoff, 2013)](https://sciwheel.com/work/citation?ids=93375&pre=&suf=&sa=0). The nucleosome is composed of 4 core histone proteins, H2A, H2B, H3 and H4, and is held together by electrostatic interactions between the negative charge of the phosphate backbone of DNA and the positively charged surface of the histones [(Bilokapic et al., 2018)](https://sciwheel.com/work/citation?ids=6823814&pre=&suf=&sa=0). The modification of core histones by acetylation and methylation is involved in allowing or repressing access to the DNA. Genome rearrangement in ciliates is influenced by processes that modify and regulate nucleosomes and consequently mediate the ability of IES-excision machinery to access the underlying DNA. In *Tetrahymena*, IESs, which frequently contain transposons or are derived from them, are targeted for removal by sRNA machinery that is involved in depositing methylation marks on Histone 3 Lysine 9 (H3K9) and Histone 3 Lysine 27 (H3K27), in a process akin to heterochromatin formation, except that the marked regions are excised entirely [(Chalker, 2008; Liu et al., 2007)](https://sciwheel.com/work/citation?ids=4034494,6344886&pre=&pre=&suf=&suf=&sa=0,0). In *Paramecium*, a mechanism reflecting the ancient ancestral eukaryotic origins of transposon silencing by heterochromatin formation, involving H3K9- and H3K27-trimethylation (H3K27me3), represseses MIC genome-encoded transposable element gene expression and experimental elimination of these marks leads to low efficiency of IES excision and lethal outcomes when new MAC genomes are produced [(Frapporti et al., 2019)](https://sciwheel.com/work/citation?ids=7102633&pre=&suf=&sa=0). A particular histone variant (H3.4) present in polytene DNA, was proposed to be the target of trimethylation, facilitating heterochromatinization and excision of IESs not protected by 27 nt macRNAs in the ciliate *Stylonychia* [(Postberg et al., 2018)](https://sciwheel.com/work/citation?ids=5413647&pre=&suf=&sa=0).

We annotated the four core histones, H2A, H2B, H3 and H4, in *B. stoltei* using the domain models from Histone DB (v2.0) (Figure S9). We found eleven putative H2A, five H2B, eleven H3 and five H4 histone proteins. Histone H2A forms dimers with histone H2B and histone H3 forms dimers with histone H4 [(Malik and Henikoff, 2003)](https://sciwheel.com/work/citation?ids=655987&pre=&suf=&sa=0). The H2B and H4 histones are known to be more conserved in comparison to H2A and H3 across several eukaryotic lineages [(Malik and Henikoff, 2003)](https://sciwheel.com/work/citation?ids=655987&pre=&suf=&sa=0). The trend of greater diversity in homologs of H2A and H3 in other eukaryotic lineages is also preserved in *Blepharisma*, where there are twice as many H2A homologs as those of H2B and almost twice as many H3 homologs as those of H4.

Unlike *Paramecium tetraurelia* and *Tetrahymena thermophila*, which have longer, divergent histone H3’s proposed to be centromeric [(Cervantes et al., 2006)](https://sciwheel.com/work/citation?ids=6345182&pre=&suf=&sa=0), i.e, CenH3, we did not observe such histones in *Blepharisma*. Both of the longer *Blepharisma* histone H3’s have unusual N-terminal domains (VIT and VWA_3, PF08487 and PF13768, respectively). In the PFAM database (34.0) the pairing of these two domains exclusively without any other domains represents the most common domain architectures for both. Searches of UniProt reveal that this domain pair is common in eukaryotes and bacteria, but it is not known what role they play in combination [(Whittaker and Hynes, 2002)](https://sciwheel.com/work/citation?ids=2171886&pre=&suf=&sa=0). Furthermore, the pair of proteins with the VIT-VWA_3 domain pair represent the most weakly expressed histone H3 domain-containing proteins in *Blepharisma*.

Since substantial upregulation of certain histone variants occurs during development in both *Oxytricha* and *Stylonychia*, including during the period of genome editing [(Aeschlimann et al., 2014; Forcob et al., 2014; Postberg et al., 2018)](https://sciwheel.com/work/citation?ids=6874065,11774648,5413647&pre=&pre=&pre=&suf=&suf=&suf=&sa=0,0,0), we examined the patterns of expression during *Blepharisma* development. Among the *Blepharisma* histones, certain candidates of three of the core histones H2A, H2B and H3 are constitutively expressed at similar levels throughout the cycle of sexual reproduction, while others are upregulated at timepoints corresponding to different stages of meiosis (6h and 14h timepoints) and also subsequently during new MAC development. The patterns of expression observed suggest that even the *Blepharisma* genome encodes variants that are likely to have a range of different functions, including in genome editing and likely also during DNA amplification in the developing new MAC. Histone H4, in contrast, appears to be expressed at relatively similar levels throughout conjugation. This constitutive expression of histone H4 is a characteristic shared among eukaryotes, which lack functional variants due their highly conserved constitution, a trait suggested to be favored by the greater necessity of this histone to maintain several protein-protein contacts with the other three histones [(Malik and Henikoff, 2003)](https://sciwheel.com/work/citation?ids=655987&pre=&suf=&sa=0).

#### PiggyBac homologs in other heterotrichs, but not the oligohymenophorean, *Ichthyophthirius multifiliis*

PiggyMac homologs are also present in other heterotrich ciliates but have not yet been described because of genome assembly or annotation challenges. Using BPgm as a query sequence, we found convincing homologs containing the conserved catalytic DDD-motif in a genome assembly of the heterotrichous ciliate *Condylostoma magnum* (TBLASTN e-value 2e-24 to 2e-37). All the *C. magnum* PiggyMac homologs have a complete DDD-catalytic triad. While we failed to detect the DDE_Tnp_1_7 domain in predicted genes of the heterotrich *Stentor* *coeruleus*, we detected relatively weak adjacent TBLASTN matches split across two frames in its draft MAC genome (e-value 7e-15; SteCoe_contig_741 positions 6558-5475). After joining ORFs corresponding to this region and translating them, we obtained a more convincing DDE_Tnp_1_7 match with HMMER3 (e-value 2e-24). This either corresponds to a pseudogene or a poorly assembled genomic region.

In addition, we searched for PiggyMac homologs in the MAC genome of the pathogenic oligohymenophorean ciliate *Ichthyophthirius multifiliis* [(Coyne et al., 2011)](https://sciwheel.com/work/citation?ids=149401&pre=&suf=&sa=0). TBLASTN searches using the *T. thermophila* Tpb2 as a query returned no hits. A HMMER search using hmmscan with a six-frame translation of the *I. multifiliis* MAC genome against the PFAM-A database also did not return any matches with independent E-values (i-E-value) less than 1. We note that based on BUSCO analyses the *I. multifiliis* genome appears to be less complete than other ciliates we examined (Figure S1) . So, a better genome assembly will be needed to investigate the possibility that PiggyBac homologs are encoded elsewhere in this MAC genome.
