## Supplementary material for "Genome editing excisase origins illuminated by somatic genome of *Blepharisma*": Table S1

**Table S1.** Comparison of MAC genome assemblies.

| **Assembly** | **Flye (v2.7) Replicate 1** | **Flye (v2.7) Replicate 2** | **Flye (v2.7) Combined** | **Flye (v2.8) Combined** | **Final assembly** |
|---|---|---|---|---|---|
| Contigs | 89 | 86 | 74 | 72 | 64 (excluding mitogenome) |
| Mean coverage (from flye.log) | 76 | 70 | 145 | 145 | NA |
| %GC | 33.3 | 33.3 | 33.4 | 32.9 | 33.6 |
| Longest contig (bp) | 2036921 | 1188116 | 1541963 | 1608201 | 1514878 |
| Assembly size (bp) | 42701284 | 43066385 | 43062848 | 42982242 | 41464486 |
| N50 | 738771 | 757357 | 799426 | 817639 | 795340 |
| Two telomeres | 38 | 37 | 36 | 16 | 64 |
| One telomere | 36 | 36 | 25 | 32 | 0 |
| Zero telomeres | 15 | 13 | 13 | 24 | 0 |
