## Supplementary material for "Genome editing excisase origins illuminated by somatic genome of *Blepharisma*": Table S2

**Table S2.** Citations for genome properties from Figure 1C.

| Species | Genome size (Mb) | Genome architecture | Genes (zygosity) | Codon reassignments |
|---|---|---|---|---|
| *Blepharisma stoltei* | 41 | Minichromosomes | 25726 (n) | UGA -> W |
| *Stentor coeruleus* | 77[2] | Not determined | 31426[2] (n) | Standard genetic code[2] |
| *Paramecium tetraurelia* | 72[3] | Chromosomes[3,4] | 39642[3] (n) | UAA, UAG -> Q[1] |
| *Tetrahymena thermophila* | 103[5] | Chromosomes[6] | 26258[5] (n) | UAA, UAG -> Q[1] |
| *Euplotes octocarinatus* | 88[7] | Nanochromosomes[8] | 29076[7] (n) | UGA -> C[9] |
| *Stylonychia lemnae* | 52[10] | Nanochromosomes[10] | 15102 (n)[10] | UAA, UAG -> Q [1] |
| *Oxytricha trifallax* | 50[11] | Nanochromosomes[11] | 18400 (n)[11] | UAA, UAG -> Q[1] |
| *Perkinsus olseni* | 63[12] | Chromosomes[12] | 17342 (4n)[12] | Standard genetic code[12] |
