## Supplementary material for "Genome editing excisase origins illuminated by somatic genome of *Blepharisma*": Table S3

**Table S3.** Comparison of *B. stoltei* intron prediction performance.

| | **AUGUSTUS*** | **Intronarrator**** |
|---|---|---|
| **True positives (TP) (real introns)** | 45 | 61 |
| **False positives (FP) (fake introns)** | 62 | 0 |
| **False negatives (FN) (missed introns)** | 15 | 2 |
| **Sensitivity: TP/(TP+FN)** | 0.75 | 0.97 |
| **Precision: TP/(TP+FP)** | 0.42 | 1.00 |

* Parameters/source code adjusted as for *Stentor* (Slabodnick et al. 2017).
** AUGUSTUS changes/parameters as in (Slabodnick et al. 2017).
