## Supplementary material for "Genome editing excisase origins illuminated by somatic genome of *Blepharisma*": Table S4

**Table S4.** Noncanonical introns (15 or 16 bp).

| Intron with 3 bp exon flanks | Length (bp) | Intronarrator/ AUGUSTUS gene ID | ENA gene accession | Spliced fraction |
|---|---|---|---|---|
| AAGgcaaattttatttagATT | 15 | Contig_10.g615 | BSTOLATCC_MAC443 | 0.827 |
| AAGgcaactataatttagAGC | 15 | Contig_11.g1292 | BSTOLATCC_MAC1123 | 0.73 |
| CGAtatgagtttacaaatTTA | 15 | Contig_36.g880 | BSTOLATCC_MAC12062 | 0.623 |
| AAGgcaaaatttaaatagAGC | 15 | Contig_43.g513 | BSTOLATCC_MAC15266 | 0.205 |
| CAGgcaatttttatttagAAG | 15 | Contig_46.g452 | BSTOLATCC_MAC16844 | 0.335 |
| ATGgcaagctctatatagAAT | 15 | Contig_49.g1050 | BSTOLATCC_MAC17453 | 0.797 |
| TTActtctataaatacacCAA | 15 | Contig_54.g273 | BSTOLATCC_MAC19826 | 0.537 |
| AAGgcaaaaaatatatagGTT | 15 | Contig_58.g1437 | BSTOLATCC_MAC22241 | 0.843 |
| GAGgcaatttttacgtagATT | 15 | Contig_59.g298 | BSTOLATCC_MAC22578 | 0.691 |
| TGAggtaaattataactagGGT | 16 | Contig_2.g441 | BSTOLATCC_MAC4446 | 0.54 |
| AAGgtaatttcccagcaggAAT | 16 | Contig_3.g1280 | BSTOLATCC_MAC9737 | 0.441 |
| CCCttgctcccctcagtagTTA | 16 | Contig_6.g757 | BSTOLATCC_MAC22759 | 0.477 |
| ATGgtaactcacaattaggCTT | 16 | Contig_7.g1329 | BSTOLATCC_MAC24915 | 0.283 |
| CACgtaaaatacaattaggAGT | 16 | Contig_12.g347 | BSTOLATCC_MAC1650 | 0.419 |
| TATggtaatttgttatcagGGA | 16 | Contig_13.g1129 | BSTOLATCC_MAC2441 | 0.326 |
| ATGtaatttaccaatagggCTA | 16 | Contig_19.g1089 | BSTOLATCC_MAC3792 | 0.867 |
| ACAgtaagatttaattaggCCT | 16 | Contig_19.g1253 | BSTOLATCC_MAC3956 | 0.525 |
| TGAgtaagatacaagtaggAGG | 16 | Contig_21.g736 | BSTOLATCC_MAC5692 | 0.644 |
| AGGtaattggcaaataggATA | 16 | Contig_24.g464 | BSTOLATCC_MAC7001 | 0.415 |
| AAGgtaaattacaagcaggAAA | 16 | Contig_25.g797 | BSTOLATCC_MAC7334 | 0.764 |
| CAAgtaattttcgaataggAAC | 16 | Contig_38.g1424 | BSTOLATCC_MAC12612 | 0.78 |
| AAGgtaatctctattaaggACA | 16 | Contig_42.g2 | BSTOLATCC_MAC14754 | 0.602 |
| AAGgcaattctctaggtagGAG | 16 | Contig_46.g332 | BSTOLATCC_MAC16724 | 0.286 |
| AGAggtaatgcataactagGGT | 16 | Contig_47.g486 | BSTOLATCC_MAC16883 | 0.648 |
| CCAgtaagtttctatttatGTC | 16 | Contig_55.g452 | BSTOLATCC_MAC20010 | 0.715 |
| AACgtaatttgtaactaggGGT | 16 | Contig_57.g559 | BSTOLATCC_MAC21349 | 0.7 |
| AAAgtaagagaccattaggTTA | 16 | Contig_57.g761 | BSTOLATCC_MAC21551 | 0.726 |
| ATTggtataggataattagGAA | 16 | Contig_60.g487 | BSTOLATCC_MAC23405 | 0.376 |
| TATacatgttttaaataatTGC | 16 | Contig_61.g1057 | BSTOLATCC_MAC23980 | 0.278 |
| AGAgtattttacaaataggCTA | 16 | Contig_63.g352 | BSTOLATCC_MAC24681 | 0.585 |

Predicted introns are in lower case; flanking exons are in upper case. Different possible donor and acceptor site pairs of bases are coloured. “Spliced fraction” indicates the efficiency of splicing calculated from Intronarrator.
