## Supplementary material for "Genome editing excisase origins illuminated by somatic genome of *Blepharisma*": Table S5

***B. stoltei* ATCC 30299 MAC gene expression** (only genes with PFAM annotations; domain multiplicitly indicated by "x" and a number; blue = 0-1 RPKM; cyan = 1-10 RPKM; 10-100 RPKM = yellow; 100-1000 RPKM = orange; 1000-10000 RPKM = red )

| | ENA accession | Gene ID | Length (CDS in bp) | Starved | Gamone-treated | 0h | 2h | 6h | 14h | 18h | 22h | 26h | 30h | 38h | 26h/average(starved +gamone+0h) | Transposase | DNA repair | Chromatin | Transcription | Small RNAs | Translation | PFAM Domains | eggNOG gene name | eggNOG description | Pannzer gene name | Pannzer description | Pannzer PPV |
|---|---|---|---|---|---|---|---|---|---|---|---|---|---|---|---|---|---|---|---|---|---|---|---|---|---|---|---|
| 1 | BSTOLATCC_MAC13665 | Contig_40.g581 | 1362 | 7712.4 | 10287.9 | 9655.0 | 9848.3 | 7732.0 | 9765.3 | 8089.2 | 8084.9 | 10403.2 | 8266.2 | 14239.1 | 1.1 | | | | | | | GTP_EFTU x1 GTP_EFTU_D3 x1 ( | TEF1 | regulation of response to stimulus | NA | Elongation factor 1-alpha | 0.46 |
| 2 | BSTOLATCC_MAC22028 | Contig_58.g1224 | 1167 | 3975.0 | 4211.8 | 4389.2 | 6931.6 | 5651.7 | 6554.9 | 3683.2 | 6009.0 | 6740.9 | 3880.1 | 5676.0 | 1.6 | | | | | | | ubiquitin x5 Rad60-SLD x5 | | ubiquitin | NA | Polyubiquitin | 0.62 |
| 3 | BSTOLATCC_MAC7206 | Contig_25.g669 | 2508 | 6164.1 | 6433.6 | 5759.2 | 5345.9 | 3753.0 | 4862.5 | 4374.3 | 3794.7 | 4592.3 | 3577.8 | 5261.9 | 0.8 | | | | | | | GTP_EFTU x1 EFG_IV x1 EFG_C | | Elongation factor G, domain IV fa | NA | Elongation factor 2 | 0.59 |
| 4 | BSTOLATCC_MAC8818 | Contig_28.g906 | 1335 | 7231.4 | 8506.6 | 10323.3 | 8424.5 | 5922.1 | 4983.5 | 3383.7 | 3704.3 | 4375.8 | 3077.5 | 3404.1 | 0.5 | | | | | | | Tubulin x1 Tubulin_C x1 Misat_Tub | | Tubulin is the major constituent of | NA | Tubulin beta chain | 0.51 |
| 5 | BSTOLATCC_MAC9322 | Contig_29.g167 | 975 | 2630.7 | 2828.7 | 2256.3 | 2164.1 | 1992.5 | 2427.7 | 2399.3 | 2720.4 | 4361.1 | 2932.7 | 3084.9 | 1.7 | | | | | | | Ldh_1_C x1 Ldh_1_N x1 | | Malate dehydrogenase | NA | Malate dehydrogenase | 0.44 |
| 6 | BSTOLATCC_MAC11989 | Contig_36.g807 | 2115 | 215.8 | 242.1 | 751.0 | 2684.2 | 2380.5 | 1191.9 | 775.8 | 1694.3 | 3605.0 | 1583.4 | 901.0 | 8.9 | | | | | | | HSP90 x1 HATPase_c x1 HATPase | HSP90 | Heat shock protein | HSP90 | Heat shock protein 90 | 0.56 |
| 7 | BSTOLATCC_MAC235 | Contig_1.g235 | 510 | 5822.9 | 4983.4 | 5291.7 | 4067.5 | 4404.2 | 3923.0 | 5056.5 | 4119.8 | 3537.3 | 4512.3 | 5802.0 | 0.7 | | | | | | | zf-C2H2 x1 zf-H2C2_2 x1 zf-Di19 x1 | | Zinc finger, C2H2 type | NA | NA | NA |
| 8 | BSTOLATCC_MAC19444 | Contig_52.g1287 | 1131 | 3118.4 | 4433.0 | 4259.3 | 3163.3 | 2784.2 | 2687.8 | 2623.6 | 2450.0 | 3346.1 | 2845.8 | 3395.0 | 0.8 | | | | | | | Actin x1Actin x1Actin x1Actin x1 | | Actin | NA | Actin | 0.6 |
| 9 | BSTOLATCC_MAC20266 | Contig_55.g708 | 1059 | 5877.8 | 8250.6 | 4377.8 | 3918.4 | 2937.8 | 3491.6 | 3508.9 | 2749.1 | 3051.9 | 3228.4 | 3229.2 | 0.5 | | | | | | | Peptidase_C1 x1 Inhibitor_I29 x1 F | | cysteine-type endopeptidase activ | NA | Cysteine protease-1 | 0.67 |
| 10 | BSTOLATCC_MAC7218 | Contig_25.g681 | 2007 | 378.4 | 539.3 | 1538.5 | 2037.1 | 3363.6 | 1283.8 | 1388.8 | 1690.3 | 2970.0 | 2671.0 | 1983.5 | 3.6 | | | | | | | HSP70 x1 MreB_Mbl x1 | HSP70 | heat shock protein 70 | HSP70 | Heat shock protein 70 | 0.55 |
| 11 | BSTOLATCC_MAC20471 | Contig_56.g905 | 1431 | 5214.7 | 5813.6 | 3727.6 | 3824.5 | 2628.6 | 3179.9 | 2384.1 | 2675.5 | 2917.4 | 2717.3 | 3373.1 | 0.6 | | | | | | | AdoHcyase x1 AdoHcyase_NAD x1 | AHCYL1 | adenosylhomocysteinase activity | NA | Adenosylhomocysteinase | 0.53 |
| 12 | BSTOLATCC_MAC19601 | Contig_54.g48 | 516 | 3534.6 | 4007.6 | 4767.6 | 2916.7 | 2869.5 | 2742.2 | 3109.5 | 2614.4 | 2814.3 | 3242.1 | 3465.8 | 0.7 | | | | | | | TCTP x1TCTP x1TCTP x1TCTP x1 | TPT1 | Translationally controlled tumour p | NA | TCTP domain-containing protein (F | 0.51 |
| 13 | BSTOLATCC_MAC17155 | Contig_48.g755 | 1896 | 7.1 | 12.3 | 9.7 | 60.2 | 3458.6 | 4149.6 | 2813.3 | 2326.9 | 2799.4 | 2446.1 | 1305.3 | 288.1 | | | | | | | DEAD x1 zf-CCHC x4 Helicase_C x | DBP2 | Belongs to the DEAD box helicase | DDX43 | RNA helicase | 0.41 |
| 14 | BSTOLATCC_MAC25659 | Contig_8.g281 | 1032 | 4697.5 | 4933.8 | 8139.4 | 4222.1 | 3275.1 | 4421.9 | 3634.2 | 2659.0 | 2744.1 | 2451.2 | 2259.2 | 0.5 | | | | | | | Gp_dh_C x1 Gp_dh_N x1 | | Glyceraldehyde-3-phosphate deh | NA | Glyceraldehyde-3-phosphate dehy | 0.45 |
| 15 | BSTOLATCC_MAC5403 | Contig_20.g450 | 513 | 3405.3 | 4181.7 | 3231.0 | 2230.0 | 3065.7 | 3322.0 | 3606.4 | 3107.9 | 2744.0 | 3524.0 | 4678.4 | 0.8 | | | | | | | Pro_isomerase x1 | -CYP 1.00 | PPIases accelerate the folding of | NA | Peptidyl-prolyl cis-trans isomerase | 0.43 |
| 16 | BSTOLATCC_MAC4350 | Contig_19.g1647 | 1056 | 4114.4 | 4806.8 | 3543.2 | 3053.1 | 2561.0 | 2509.8 | 2578.3 | 2527.8 | 2590.2 | 2066.0 | 2990.5 | 0.6 | | | | | | | Peptidase_C1 x1 Inhibitor_I29 x1 F | | cysteine-type endopeptidase activ | NA | Fruit bromelain | 0.44 |
| 17 | BSTOLATCC_MAC23581 | Contig_60.g663 | 1536 | 3206.6 | 3985.2 | 2509.7 | 2399.0 | 2001.0 | 2388.3 | 2276.9 | 2296.2 | 2485.5 | 2239.1 | 3020.2 | 0.8 | | | | | | | Aldedh x1Aldedh x1Aldedh x1 | ALDH7A1 | Belongs to the aldehyde dehydro | NA | Aldedh domain-containing protein | 0.43 |
| 18 | BSTOLATCC_MAC21846 | Contig_58.g1042 | 357 | 5819.0 | 4540.0 | 5027.4 | 2697.9 | 2582.4 | 3060.3 | 3909.7 | 2801.6 | 2388.8 | 3025.0 | 5348.3 | 0.5 | | | | | | | Bac_globin x1Bac_globin x1 | glbN | Bacterial-like globin | NA | NA | NA |
| 19 | BSTOLATCC_MAC23653 | Contig_60.g735 | 714 | 3374.9 | 4042.0 | 3121.1 | 2392.3 | 2075.6 | 2330.7 | 2216.5 | 2130.3 | 2384.1 | 2240.2 | 2798.2 | 0.7 | | | | | | | EF1_GNE x1 GST_C_2 x1 GST_C | CARS | EF-1 guanine nucleotide exchang | NA | Elongation factor 1-beta | 0.3 |
| 20 | BSTOLATCC_MAC5406 | Contig_20.g453 | 2340 | 4.5 | 9.5 | 10.1 | 150.6 | 6059.2 | 4134.1 | 3321.1 | 2959.9 | 2282.3 | 1941.8 | 1102.0 | 283.3 | | | | | ◉ | | Piwi x1 PAZ x1 ArgoL1 x1 | | Piwi | NA | NA | NA |
| 21 | BSTOLATCC_MAC1451 | Contig_12.g148 | 549 | 2.6 | 144.1 | 108.6 | 1282.1 | 6916.5 | 2971.2 | 2324.0 | 2534.4 | 2277.1 | 3006.5 | 1805.7 | 26.8 | | | | | | | HMG_box x1 HMG_box_2 x1 | NA | NA | NA | NA | NA |
| 22 | BSTOLATCC_MAC8375 | Contig_27.g470 | 780 | 2039.5 | 2458.6 | 2955.8 | 2100.5 | 1981.4 | 1761.8 | 1555.8 | 1534.1 | 2248.0 | 1885.6 | 1392.6 | 0.9 | | | | | | | Ribosomal_S3Ae x1 | RPS3A | Belongs to the eukaryotic ribosom | RPS1 | 40S ribosomal protein S1 | 0.5 |
| 23 | BSTOLATCC_MAC5529 | Contig_20.g576 | 1032 | 2287.3 | 2280.0 | 2299.5 | 1384.4 | 1432.1 | 2541.4 | 2859.1 | 2191.6 | 2222.4 | 2210.9 | 2857.6 | 1.0 | | | | | | | Gp_dh_C x1 Gp_dh_N x1 | | Glyceraldehyde-3-phosphate deh | NA | Glyceraldehyde-3-phosphate dehy | 0.46 |
| 24 | BSTOLATCC_MAC15966 | Contig_45.g1208 | 636 | 3350.5 | 3972.7 | 1980.9 | 1900.7 | 1449.2 | 1978.8 | 2620.6 | 2282.5 | 2210.5 | 1920.2 | 2184.0 | 0.7 | | | | | | | GST_C_3 x1 GST_C x1 GST_N x1 | | Glutathione S-transferase, N-term | NA | Glutathione S-transferase | 0.34 |
| 25 | BSTOLATCC_MAC6598 | Contig_23.g65 | 603 | 1847.7 | 1840.8 | 1671.4 | 2258.1 | 1863.9 | 1849.0 | 1386.2 | 1801.7 | 2179.3 | 1605.9 | 1528.1 | 1.2 | | | | | | | Ribosomal_S8e x1 | | Ribosomal protein S8e | NSA2 | Ribosome biogenesis protein NSA | 0.73 |
| 26 | BSTOLATCC_MAC13450 | Contig_4.g162 | 636 | 1683.7 | 2134.8 | 1907.3 | 1483.4 | 1631.6 | 1818.6 | 2362.4 | 1813.8 | 2127.3 | 2423.0 | 2620.5 | 1.1 | | | | | | | AhpC-TSA x1 Redoxin x1 1-cysPrx_ | | Tsa family | NA | peroxiredoxin-2 | 0.41 |
| 27 | BSTOLATCC_MAC13519 | Contig_4.g231 | 519 | 2440.5 | 2027.7 | 2274.5 | 1940.0 | 2106.9 | 2047.4 | 2374.6 | 2204.6 | 2058.3 | 2441.9 | 3145.9 | 0.9 | | | | | | | zf-C2H2 x1 zf-Di19 x1 | | nucleic acid binding | NA | NA | NA |
| 28 | BSTOLATCC_MAC23948 | Contig_61.g1025 | 906 | 5517.7 | 6461.1 | 3489.9 | 3076.5 | 2617.4 | 3574.9 | 4928.6 | 3425.4 | 2050.1 | 2395.9 | 5042.1 | 0.4 | | | | | | | Peptidase_C1 x1Peptidase_C1 x1 | | Belongs to the peptidase C1 fami | NA | Cathepsin B | 0.5 |
| 29 | BSTOLATCC_MAC13751 | Contig_40.g667 | 648 | 2414.2 | 2557.0 | 1914.2 | 1585.5 | 1315.9 | 1266.1 | 2249.7 | 1899.4 | 2046.8 | 2184.6 | 4272.1 | 0.9 | | | | | | | Sod_Fe_C x1 Sod_Fe_N x1 | | oxidoreductase activity, acting on | NA | Superoxide dismutase | 0.44 |
| 30 | BSTOLATCC_MAC23158 | Contig_6.g1156 | 1191 | 1923.5 | 2440.4 | 2780.1 | 2108.5 | 1717.2 | 1700.1 | 1194.0 | 1270.5 | 1982.4 | 1626.2 | 1135.3 | 0.8 | | | | | | | Ribosomal_L3 x1Ribosomal_L3 x1 | RPL3 | 60S ribosomal protein L3. Source | NA | 60S ribosomal protein L3 | 0.56 |
| 31 | BSTOLATCC_MAC7493 | Contig_25.g956 | 1362 | 4720.6 | 5233.4 | 3802.4 | 3394.9 | 2205.8 | 2827.2 | 2306.9 | 2192.2 | 1798.9 | 1545.4 | 2192.1 | 0.4 | | | | | | | Tubulin x1 Tubulin_C x1 | | Tubulin is the major constituent of | NA | Tubulin alpha chain | 0.54 |
| 32 | BSTOLATCC_MAC7844 | Contig_25.g1307 | 1137 | 883.7 | 867.1 | 1279.8 | 1696.4 | 1262.5 | 1326.6 | 759.9 | 1171.8 | 1783.6 | 998.4 | 675.1 | 1.8 | | | | | | | Alba x2Alba x2Alba x2Alba x2 | | Alba | NA | Alba domain-containing protein | 0.37 |
| 33 | BSTOLATCC_MAC6369 | Contig_22.g1410 | 1047 | 1680.0 | 2138.4 | 1923.7 | 1804.7 | 1383.5 | 1303.4 | 966.8 | 1170.8 | 1756.3 | 1291.7 | 1170.4 | 0.9 | | | | | | | Ribosomal_L4 x1 Ribos_L4_asso_ | | Ribosomal protein L4 | NA | 60S ribosomal protein L4 | 0.7 |
| 34 | BSTOLATCC_MAC22535 | Contig_59.g254 | 276 | 1560.1 | 1455.2 | 3302.0 | 1940.2 | 2831.7 | 1813.0 | 1763.9 | 1590.8 | 1752.8 | 2169.4 | 1685.2 | 0.8 | | | | | | | EF-hand_1 x2 EF-hand_5 x2 EF-ha | NA | NA | NA | Calmodulin | 0.5 |
| 35 | BSTOLATCC_MAC642 | Contig_10.g814 | 1080 | 2315.2 | 2854.8 | 2198.4 | 1762.5 | 1691.9 | 2057.9 | 2051.2 | 1826.7 | 1729.8 | 1733.5 | 1759.9 | 0.7 | | | | | | | Glycolytic x1Glycolytic x1 | | fructose-bisphosphate aldolase | NA | Fructose-bisphosphate aldolase | 0.5 |
| 36 | BSTOLATCC_MAC21506 | Contig_57.g716 | 672 | 3025.4 | 3615.0 | 2467.2 | 1640.4 | 1818.7 | 2011.7 | 2485.1 | 1920.8 | 1724.9 | 2053.6 | 3295.7 | 0.6 | | | | | | | Ribosomal_S3_C x1 KH_2 x1 | RpS3 | positive regulation of DNA N-glyco | NA | KH type-2 domain-containing prote | 0.53 |
| 37 | BSTOLATCC_MAC21995 | Contig_58.g1191 | 414 | 9.2 | 38.2 | 33.0 | 61.5 | 906.8 | 918.9 | 1414.2 | 1735.4 | 1722.8 | 2325.4 | 2117.3 | 64.3 | | | | | | | Histone x1Histone x1Histone x1 | | protein heterodimerization activity | NA | Histone domain-containing protein | 0.51 |
| 38 | BSTOLATCC_MAC250 | Contig_1.g250 | 798 | 3077.2 | 3710.4 | 2591.3 | 1810.0 | 1723.2 | 1976.1 | 1944.8 | 1793.8 | 1695.0 | 1746.4 | 2403.0 | 0.5 | | | | | | | Ribosomal_S5 x1 Ribosomal_S5_C | RPS2 | translation | NA | Small subunit ribosomal protein S2 | 0.56 |
| 39 | BSTOLATCC_MAC8624 | Contig_28.g712 | 1407 | 2824.7 | 3304.2 | 2192.3 | 2128.8 | 1631.8 | 1887.4 | 2089.1 | 1575.1 | 1657.4 | 1609.9 | 1823.8 | 0.6 | | | | | | | EF1G x1 GST_C x1 GST_C_2 x1 ( | EEF1G | translation elongation factor activi | NA | NA | NA |
| 40 | BSTOLATCC_MAC7002 | Contig_24.g465 | 357 | 2129.9 | 2285.3 | 3106.6 | 1620.7 | 1509.6 | 1530.1 | 1803.2 | 1319.1 | 1580.3 | 1737.1 | 1821.8 | 0.6 | | | | | | | SCP2 x1 Alkyl_sulf_C x1 | SCP2D1 | SCP-2 sterol transfer family | NA | SCP2 domain-containing protein | 0.49 |
| 41 | BSTOLATCC_MAC13544 | Contig_4.g256 | 510 | 3181.1 | 2961.3 | 1910.5 | 1409.6 | 1399.5 | 2159.9 | 2191.9 | 2262.9 | 1547.7 | 1816.7 | 4250.7 | 0.6 | | | | | | | zf-C2H2 x2 zf-Di19 x1 | NA | NA | NA | NA | NA |
| 42 | BSTOLATCC_MAC3386 | Contig_18.g685 | 747 | 2018.5 | 2245.7 | 1701.1 | 1532.1 | 1145.4 | 1435.8 | 1217.2 | 1278.2 | 1525.8 | 1168.3 | 1243.6 | 0.8 | | | | | | | Ribosomal_L7Ae x1 | | Ribosomal protein L7Ae/L30e/S12 | NA | 60S ribosomal protein L8 | 0.39 |
| 43 | BSTOLATCC_MAC14030 | Contig_40.g946 | 546 | 0.4 | 33.4 | 17.0 | 721.8 | 4701.6 | 2241.2 | 3048.8 | 3099.9 | 1508.7 | 1231.4 | 861.7 | 89.1 | | | ◉ | | | | HMG_box x1 HMG_box_2 x1 | NA | NA | NA | NA | NA |
| 44 | BSTOLATCC_MAC23079 | Contig_6.g1077 | 792 | 2398.2 | 3165.3 | 1882.2 | 1579.9 | 1418.9 | 1584.3 | 1508.9 | 1358.0 | 1504.5 | 1421.4 | 1653.2 | 0.6 | | | | | | | Ribosomal_S4e x1 40S_S4_C x1 F | RPS4 | Belongs to the eukaryotic ribosom | NA | Ribosomal protein S4, putative | 0.81 |
| 45 | BSTOLATCC_MAC9923 | Contig_30.g352 | 1101 | 3564.2 | 3111.2 | 1592.2 | 1555.3 | 654.7 | 2955.4 | 1652.6 | 2141.4 | 1502.8 | 1145.0 | 2306.9 | 0.5 | | | | | | | Fungal_lectin x2Fungal_lectin x2 | NA | NA | NA | NA | NA |
| 46 | BSTOLATCC_MAC7895 | Contig_25.g1358 | 948 | 1844.3 | 2096.4 | 1300.4 | 1761.4 | 1069.4 | 1445.3 | 969.1 | 1202.4 | 1486.7 | 924.8 | 817.0 | 0.9 | | | | | | | Ribosomal_L5e x1 Ribosomal_L18 | RPL5 | ribosomal protein | NA | 60S ribosomal protein L5 | 0.53 |
| 47 | BSTOLATCC_MAC23089 | Contig_6.g1087 | 1002 | 1565.5 | 1864.4 | 1670.3 | 1286.9 | 968.8 | 1118.8 | 1086.9 | 1002.8 | 1483.3 | 1096.9 | 1158.8 | 0.9 | | | | | | | RL10P_insert x1 Ribosomal_L10 x1 | RPP0 | Component of the ribosome, a lar | NA | 60S acidic ribosomal protein P0 | 0.7 |
| 48 | BSTOLATCC_MAC5313 | Contig_20.g360 | 1104 | 2156.7 | 2673.6 | 2735.9 | 1671.6 | 1442.9 | 1401.4 | 1535.3 | 941.5 | 1431.6 | 1633.4 | 1382.2 | 0.6 | | | | | | | Asp x1 TAXi_N x1 | | Belongs to the peptidase A1 fami | NA | Cathepsin D | 0.8 |
| 49 | BSTOLATCC_MAC3156 | Contig_17.g456 | 1062 | 2342.1 | 2915.2 | 2999.7 | 2129.8 | 1494.8 | 2019.0 | 1901.5 | 1543.0 | 1419.3 | 1218.7 | 1719.6 | 0.5 | | | | | | | Peptidase_C1 x1 Inhibitor_I29 x1 | | Belongs to the peptidase C1 fami | NA | Senescence-specific cysteine prot | 0.69 |
| 50 | BSTOLATCC_MAC3991 | Contig_19.g1288 | 2433 | 285.7 | 337.2 | 384.1 | 1419.2 | 825.1 | 921.9 | 446.4 | 931.2 | 1398.0 | 549.7 | 417.5 | 4.2 | | | | | | | Iso_dh x1Iso_dh x1Iso_dh x1 | IDH1 | Belongs to the isocitrate and isop | NA | NA | NA |
| 51 | BSTOLATCC_MAC4112 | Contig_19.g1409 | 1050 | 3000.8 | 3630.0 | 2142.1 | 1687.7 | 1443.7 | 1812.2 | 2101.1 | 1644.2 | 1389.6 | 1558.3 | 2231.4 | 0.5 | | | | | | | Peptidase_C1 x1 Inhibitor_I29 x1 | | cysteine-type endopeptidase activ | NA | Cathepsin L | 0.44 |
| 52 | BSTOLATCC_MAC1449 | Contig_12.g146 | 631 | 961.1 | 1107.5 | 1458.1 | 1298.5 | 1391.3 | 1074.9 | 1015.1 | 1091.6 | 1386.6 | 1257.9 | 1058.8 | 1.2 | | | | | | | Ras x1 Roc x1 Arf x1 MMR_HSR1 x | | Rab subfamily of small GTPases | NA | Rab protein 7 | 0.88 |
| 53 | BSTOLATCC_MAC11656 | Contig_35.g476 | 1362 | 3507.0 | 3286.3 | 4024.4 | 3353.9 | 2466.6 | 2646.2 | 1939.1 | 1490.4 | 1386.0 | 1488.1 | 1261.2 | 0.4 | | | | | | | Tubulin x1 Tubulin_C x1 | | Tubulin is the major constituent of | NA | Tubulin alpha chain | 0.54 |
| 54 | BSTOLATCC_MAC11524 | Contig_34.g345 | 777 | 1844.2 | 2422.6 | 1886.1 | 1488.6 | 1476.7 | 1255.0 | 1273.3 | 1168.4 | 1382.3 | 1419.4 | 1484.9 | 0.7 | | | | | | | Ribosomal_L2_C x1 Ribosomal_L2 | RPL8 | structural constituent of ribosome | NA | 60S ribosomal protein L8 | 0.54 |
| 55 | BSTOLATCC_MAC4721 | Contig_2.g717 | 777 | 1794.4 | 2257.6 | 1936.5 | 1248.0 | 1249.2 | 1228.7 | 1412.6 | 1143.3 | 1380.8 | 1501.9 | 1774.6 | 0.7 | | | | | | | Ribosomal_L2_C x1 Ribosomal_L2 | RPL8 | structural constituent of ribosome | NA | 60S ribosomal protein L8 | 0.54 |
| 56 | BSTOLATCC_MAC23186 | Contig_6.g1184 | 780 | 1097.4 | 1118.3 | 1361.4 | 1077.0 | 953.5 | 1019.5 | 766.3 | 930.5 | 1376.6 | 990.8 | 729.5 | 1.2 | | | | | | | Ribosomal_S3Ae x1 | RPS3A | Belongs to the eukaryotic ribosom | RPS1 | 40S ribosomal protein S1 | 0.49 |
| 57 | BSTOLATCC_MAC15760 | Contig_44.g1003 | 576 | 7.8 | 29.4 | 16.7 | 102.1 | 822.0 | 697.4 | 840.6 | 1266.0 | 1370.2 | 1903.7 | 1632.8 | 76.3 | | | | | | | SAM_1 x1 SAM_2 x1 | NA | NA | NA | NA | NA |
| 58 | BSTOLATCC_MAC18158 | Contig_50.g10 | 906 | 2616.2 | 3282.9 | 2264.8 | 1798.8 | 1648.1 | 1530.6 | 2119.5 | 1524.9 | 1335.9 | 1538.9 | 2767.7 | 0.5 | | | | | | | Peptidase_C1 x1Peptidase_C1 x1 | | Belongs to the peptidase C1 fami | NA | Cathepsin B | 0.59 |
| 59 | BSTOLATCC_MAC15584 | Contig_44.g827 | 990 | 2439.4 | 2946.9 | 2070.6 | 1392.1 | 1137.2 | 1258.7 | 1472.9 | 1188.8 | 1262.4 | 1376.8 | 1628.3 | 0.5 | | | | | | | WD40 x5WD40 x5WD40 x5 | RACK1A | component of 40S small ribosoma | NA | Guanine nucleotide-binding protei | 0.58 |
| 60 | BSTOLATCC_MAC5726 | Contig_21.g770 | 939 | 3.7 | 6.5 | 3365.7 | 4971.6 | 3160.3 | 230.6 | 453.9 | 498.2 | 1260.5 | 3683.6 | 240.5 | 1.1 | | | | | | | Pkinase x1 PK_Tyr_Ser-Thr x1 | CDC28 | cyclin-dependent protein serine/th | NA | Cell division control protein 2 | 0.62 |
| 61 | BSTOLATCC_MAC8364 | Contig_27.g459 | 402 | 1604.4 | 1635.8 | 1722.8 | 1189.5 | 1176.4 | 1195.1 | 1252.2 | 1062.9 | 1238.6 | 1224.6 | 1189.2 | 0.7 | | | | | | | Ribosomal_L32e x1 | | 60S ribosomal protein L32-1-like | RPL32 | 60S ribosomal protein L32 | 0.72 |
| 62 | BSTOLATCC_MAC19969 | Contig_55.g411 | 747 | 1195.7 | 1078.4 | 1612.8 | 1057.7 | 1097.7 | 1075.9 | 1071.1 | 1062.7 | 1236.9 | 1240.9 | 1290.5 | 1.0 | | | | | | | 14-3-3 x114-3-3 x114-3-3 x1 | | Belongs to the 14-3-3 family | NA | NA | NA |
| 63 | BSTOLATCC_MAC11303 | Contig_34.g124 | 414 | 14.7 | 38.3 | 16.2 | 45.6 | 888.7 | 732.1 | 1263.0 | 1417.3 | 1216.9 | 1734.7 | 1857.8 | 52.8 | | | | | | | Histone x1Histone x1Histone x1 | | protein heterodimerization activity | NA | Histone domain-containing protein | 0.5 |
| 64 | BSTOLATCC_MAC4978 | Contig_20.g25 | 510 | 1164.2 | 1736.1 | 1016.9 | 741.3 | 986.2 | 1378.1 | 1588.1 | 1396.5 | 1207.1 | 1275.1 | 1750.5 | 0.9 | | | | | | | zf-C2H2 x1 zf-H2C2_2 x1 zf-Di19 x1 | | Zinc-finger of C2H2 type | NA | NA | NA |
| 65 | BSTOLATCC_MAC21189 | Contig_57.g399 | 351 | 6.0 | 92.0 | 46.6 | 260.6 | 1419.6 | 786.0 | 1028.8 | 1533.8 | 1189.8 | 1752.0 | 1529.1 | 24.7 | | | | | | | Histone x1Histone x1Histone x1 | | Core component of nucleosome. I | NA | Histone domain-containing protein | 0.1 |
| 66 | BSTOLATCC_MAC23535 | Contig_60.g617 | 426 | 1951.9 | 2072.1 | 1989.8 | 1278.6 | 1307.4 | 1198.7 | 1424.3 | 1064.3 | 1184.8 | 1381.2 | 1544.7 | 0.6 | | | | | | | Ribosomal_L27A x1 | | Belongs to the universal ribosoma | NA | RL17A, ribosomal protein 27A 60S | 0.58 |
| 67 | BSTOLATCC_MAC10015 | Contig_30.g444 | 1335 | 2522.3 | 2394.9 | 2575.5 | 2051.9 | 1457.5 | 1712.8 | 1396.8 | 1166.4 | 1183.3 | 991.8 | 1034.9 | 0.5 | | | | | | | Tubulin x1 Tubulin_C x1 Misat_Tub | | Tubulin is the major constituent of | NA | Tubulin beta chain | 0.51 |
| 68 | BSTOLATCC_MAC3117 | Contig_17.g417 | 462 | 2418.9 | 2259.4 | 2668.4 | 1571.9 | 1684.5 | 1703.9 | 1962.8 | 1612.6 | 1168.6 | 1373.3 | 1986.8 | 0.5 | | | | | | | NDK x1NDK x1NDK x1NDK x1 | awd | nucleoside diphosphate kinase ac | NDK | Nucleoside diphosphate kinase | 0.45 |
| 69 | BSTOLATCC_MAC5291 | Contig_20.g338 | 543 | 1.7 | 5.4 | 3.1 | 102.8 | 2131.6 | 1550.9 | 2462.4 | 1970.2 | 1161.6 | 1535.0 | 1168.4 | 339.5 | | | | | | ◉ | IF4E x1IF4E x1IF4E x1IF4E x1 | | translation initiation factor activity | NA | Eukaryotic translation initiation fac | 0.22 |
| 70 | BSTOLATCC_MAC11000 | Contig_33.g1425 | 621 | 1997.7 | 2645.5 | 1598.4 | 1364.8 | 1148.5 | 1193.7 | 1304.5 | 1285.3 | 1156.0 | 1221.4 | 1526.2 | 0.6 | | | | | | | Ribosomal_S8e x1 | RPS8 | Belongs to the eukaryotic ribosom | NA | 40S ribosomal protein S8 | 0.67 |
| 71 | BSTOLATCC_MAC22403 | Contig_59.g122 | 615 | 1526.4 | 1602.6 | 1772.2 | 1039.2 | 959.5 | 1080.1 | 885.5 | 937.0 | 1142.5 | 1068.6 | 998.2 | 0.7 | | | | | | | Ribosomal_L15e x1 | | Belongs to the eukaryotic ribosom | RPL15 | Ribosomal protein L15 | 0.67 |
| 72 | BSTOLATCC_MAC15091 | Contig_42.g339 | 276 | 2724.6 | 2192.3 | 1422.6 | 1265.5 | 1448.1 | 1946.0 | 2100.0 | 1627.9 | 1134.6 | 1499.9 | 2463.1 | 0.5 | | | | | | | EF-hand_1 x2 EF-hand_5 x2 EF-ha | NA | NA | NA | Calmodulin (Fragment) | 0.4 |
| 73 | BSTOLATCC_MAC12326 | Contig_38.g1138 | 396 | 1058.5 | 1184.9 | 1492.7 | 1083.0 | 1179.6 | 968.5 | 1037.7 | 998.0 | 1122.5 | 1214.4 | 1317.0 | 0.9 | | | | | | | DUF2423 x1DUF2423 x1 | NA | NA | NA | NA | NA |
| 74 | BSTOLATCC_MAC24658 | Contig_63.g328 | 366 | 1594.3 | 1614.5 | 2024.6 | 1060.8 | 1040.7 | 993.6 | 1058.6 | 719.6 | 1103.0 | 1121.5 | 1116.7 | 0.6 | | | | | | | Ribosomal_S25 x1 | RPS25B | 40S ribosomal protein S25 | NA | 40S ribosomal protein S25 | 0.6 |
| 75 | BSTOLATCC_MAC12160 | Contig_36.g978 | 1527 | 1277.2 | 1446.8 | 1101.9 | 1336.4 | 756.7 | 1065.1 | 693.0 | 982.8 | 1092.5 | 710.8 | 1020.7 | 0.9 | | | | | | | Aldedh x1Aldedh x1Aldedh x1 | | Aldehyde dehydrogenase family | MMSA | Methylmalonate-semialdehyde deh | 0.49 |
| 76 | BSTOLATCC_MAC19478 | Contig_52.g1321 | 792 | 1211.9 | 1404.5 | 1089.6 | 954.8 | 775.5 | 970.4 | 858.7 | 904.3 | 1087.0 | 885.2 | 1073.8 | 0.9 | | | | | | | Ribosomal_S4e x1 40S_S4_C x1 F | RPS4 | Belongs to the eukaryotic ribosom | NA | Ribosomal protein S4, putative | 0.8 |
| 77 | BSTOLATCC_MAC4515 | Contig_2.g510 | 606 | 1624.3 | 1956.7 | 1671.7 | 1201.9 | 1101.8 | 1059.1 | 1169.7 | 970.9 | 1080.6 | 1159.7 | 1193.4 | 0.6 | | | | | | | Ribosomal_L13 x1 | RPL13A | Belongs to the universal ribosoma | NA | Large subunit ribosomal protein L1 | 0.72 |
| 78 | BSTOLATCC_MAC3660 | Contig_19.g957 | 504 | 1428.6 | 1486.1 | 1293.0 | 899.3 | 1007.0 | 1029.3 | 1226.0 | 984.1 | 1080.6 | 1295.7 | 1451.0 | 0.8 | | | | | | | Profilin x1Profilin x1Profilin x1 | | Profilin | NA | Profilin | 0.56 |
| 79 | BSTOLATCC_MAC13936 | Contig_40.g852 | 1467 | 963.8 | 953.4 | 1158.0 | 971.2 | 585.4 | 717.1 | 559.4 | 704.9 | 1078.0 | 713.0 | 863.6 | 1.1 | | | | | | | ELFV_dehydrog x1 ELFV_dehydro | gdhA | Belongs to the Glu Leu Phe Val d | GDHA | Glutamate dehydrogenase | 0.45 |
| 80 | BSTOLATCC_MAC12384 | Contig_38.g1196 | 1617 | 1153.0 | 1085.3 | 803.8 | 1091.8 | 808.6 | 1031.9 | 760.0 | 932.5 | 1061.7 | 780.6 | 885.4 | 1.0 | | | | | | | DEAD x1 Helicase_C x1 ResIII x1 E | DBP2 | Belongs to the DEAD box helicase | NA | Similar to Saccharomyces cerevisia | 0.42 |
| 81 | BSTOLATCC_MAC18506 | Contig_50.g358 | 390 | 1652.8 | 1546.8 | 1276.8 | 990.1 | 876.0 | 1007.2 | 1112.3 | 1004.6 | 1060.2 | 1029.5 | 1357.6 | 0.7 | | | | | | | ubiquitin x1 Ribosomal_L40e x1 Ra | | Ribosomal L40e family | NA | Ubiquitin | 0.59 |
| 82 | BSTOLATCC_MAC9163 | Contig_29.g8 | 1146 | 322.1 | 764.9 | 981.8 | 1642.7 | 1150.3 | 782.0 | 837.5 | 854.3 | 1051.2 | 930.4 | 537.8 | 1.5 | | | | | | | ubiquitin x5 Rad60-SLD x5 | | ubiquitin | NA | Polyubiquitin | 0.64 |

| | | | | | | | | | | | | | | | | | | | | | | | | | | | |
|---|---|---|---|---|---|---|---|---|---|---|---|---|---|---|---|---|---|---|---|---|---|---|---|---|---|---|---|
| 83 | BSTOLATCC_MAC15740 | Contig_44.g983 | 531 | 2170.2 | 2017.5 | 1904.8 | 1409.7 | 1295.7 | 1480.2 | 1567.8 | 1326.1 | 1044.9 | 1023.8 | 1088.3 | 0.5 | | | | | | | EF-hand_7 x2 EF-hand_6 x1 EF-ha | CEN2 | EF-hand domain | NA | Centrin-2 | 0.69 |
| 84 | BSTOLATCC_MAC23416 | Contig_60.g498 | 1221 | 986.0 | 1020.5 | 1400.6 | 1276.0 | 1013.4 | 1143.0 | 690.6 | 858.4 | 1042.8 | 789.1 | 697.3 | 0.9 | | | | | | | 2-Hacid_dh_C x1 2-Hacid_dh x1 N/ | | D-isomer specific 2-hydroxyacid de | NA | 2-oxoglutarate reductase | 0.5 |
| 85 | BSTOLATCC_MAC11902 | Contig_36.g720 | 639 | 1054.1 | 1503.7 | 1370.5 | 1010.2 | 1100.5 | 849.7 | 864.3 | 868.5 | 1039.6 | 1045.7 | 1039.4 | 0.8 | | | | | | | Ribosomal_L16 x1 | | Ribosomal protein L16p/L10e | RL10 | 60S ribosomal protein L10 | 0.58 |
| 86 | BSTOLATCC_MAC22770 | Contig_6.g768 | 1812 | 172.7 | 257.9 | 299.9 | 500.3 | 1129.4 | 554.5 | 785.2 | 740.6 | 1037.4 | 1180.0 | 773.3 | 4.3 | | | | | | | RRM_1 x4 PABP x1 | | Binds the poly(A) tail of mRNA | NA | NA | NA |
| 87 | BSTOLATCC_MAC23362 | Contig_60.g444 | 462 | 833.8 | 1204.5 | 931.9 | 637.9 | 843.8 | 1215.5 | 1237.9 | 1213.2 | 1030.3 | 991.6 | 2040.4 | 1.0 | | | | | | | zf-CCCH x1 zf_CCCH_4 x1 zf-CCCH | | 3'-UTR-mediated mRNA destabiliz | NA | NA | NA |
| 88 | BSTOLATCC_MAC3493 | Contig_18.g792 | 1422 | 507.9 | 748.5 | 581.7 | 616.4 | 666.4 | 642.3 | 726.2 | 680.8 | 1011.5 | 890.7 | 877.7 | 1.7 | | | | | | | Peptidase_M20 x1 M20_dimer x1 | dapE1 | peptidase M20 | NA | Acetylornithine deacetylase/Succir | 0.39 |
| 89 | BSTOLATCC_MAC9037 | Contig_28.g1125 | 456 | 2296.6 | 2347.8 | 1657.8 | 1186.2 | 1135.5 | 1352.8 | 1538.3 | 1347.4 | 1011.3 | 1230.3 | 1995.7 | 0.5 | | | | | | | Ribosomal_S13_N x1 Ribosomal_S | RPS13 | Belongs to the universal ribosoma | NA | 40S ribosomal protein S13 | 0.58 |
| 90 | BSTOLATCC_MAC1654 | Contig_12.g351 | 444 | 2219.0 | 2381.2 | 1516.5 | 1045.8 | 967.4 | 1218.8 | 1727.8 | 1256.8 | 1010.2 | 1014.9 | 1699.8 | 0.5 | | | | | | | UQ_con x1UQ_con x1UQ_con x1 | ubcB | Ubiquitin-conjugating enzyme E2, | NA | Ubiquitin conjugating enzyme, put | 0.49 |
| 91 | BSTOLATCC_MAC12990 | Contig_39.g185 | 462 | 483.2 | 603.4 | 1159.0 | 1623.8 | 1354.6 | 648.2 | 793.6 | 672.7 | 997.6 | 1151.5 | 736.1 | 1.3 | | | | | | | ubiquitin x2 Rad60-SLD x2 Ubiquiti | UBI4 | ubiquitin | NA | Polyubiquitin | 0.62 |
| 92 | BSTOLATCC_MAC1604 | Contig_12.g301 | 1467 | 1818.6 | 2022.6 | 1007.9 | 960.0 | 762.9 | 1194.4 | 970.5 | 1230.5 | 995.6 | 940.3 | 1481.2 | 0.6 | | | | | | | PALP x1 CBS x1PALP x1 CBS x1 | CYS4 | Belongs to the cysteine synthase | NA | Cystathionine beta-synthase | 0.59 |
| 93 | BSTOLATCC_MAC21712 | Contig_57.g922 | 453 | 1816.4 | 1609.7 | 1416.0 | 1063.6 | 881.1 | 1190.5 | 1002.8 | 1092.3 | 994.9 | 1049.0 | 1240.8 | 0.6 | | | | | | | Ribosomal_L24e x1 | RPL24 | 60S ribosomal protein L24. Sourc | NA | 60S ribosomal protein L24 | 0.45 |
| 94 | BSTOLATCC_MAC25165 | Contig_7.g1579 | 930 | 1404.3 | 1352.5 | 1995.0 | 1104.5 | 1041.2 | 1080.8 | 1375.4 | 862.6 | 986.0 | 1148.2 | 1049.7 | 0.6 | | | | | | | Mito_carr x3Mito_carr x3 | SLC25A4 | Belongs to the mitochondrial carri | NA | ADP/ATP translocase 1 | 0.56 |
| 95 | BSTOLATCC_MAC22656 | Contig_6.g654 | 672 | 2862.0 | 3056.3 | 1808.2 | 1534.0 | 950.2 | 2245.8 | 1981.5 | 1819.9 | 981.6 | 902.6 | 1872.9 | 0.4 | | | | | | | START x1START x1START x1 | NA | NA | NA | NA | NA |
| 96 | BSTOLATCC_MAC4293 | Contig_19.g1590 | 422 | 1357.8 | 1061.5 | 1511.0 | 1205.4 | 980.8 | 1080.7 | 976.6 | 841.7 | 973.5 | 987.7 | 870.8 | 0.7 | | | | | | | Ribosomal_L14e x1 | RPL14 | ribosomal protein | RPL14 | 60S ribosomal protein L14 | 0.85 |
| 97 | BSTOLATCC_MAC9061 | Contig_28.g1149 | 798 | 1905.3 | 2230.7 | 1628.3 | 1114.6 | 1099.9 | 1207.2 | 1409.5 | 998.4 | 968.7 | 1074.9 | 1726.6 | 0.5 | | | | | | | C2 x1C2 x1C2 x1C2 x1C2 x1C2 x1 | | Protein kinase C conserved regior | NA | C2 domain-containing protein | 0.46 |
| 98 | BSTOLATCC_MAC11139 | Contig_33.g1564 | 639 | 2449.1 | 3034.7 | 1652.6 | 1289.7 | 1166.2 | 1313.1 | 1439.4 | 1227.6 | 958.0 | 1050.9 | 1427.8 | 0.4 | | | | | | | Ribosomal_L16 x1 | | Ribosomal protein L16p/L10e | RL10 | 60S ribosomal protein L10 | 0.58 |
| 99 | BSTOLATCC_MAC16146 | Contig_45.g1388 | 432 | 1454.3 | 1422.0 | 1644.9 | 963.9 | 964.2 | 935.9 | 1173.3 | 912.6 | 936.7 | 1049.9 | 1090.1 | 0.6 | | | | | | | Ribosom_S12_S23 x1 | | 40s ribosomal protein | NA | 40S ribosomal protein S23 | 0.61 |
| ## | BSTOLATCC_MAC23848 | Contig_61.g925 | 876 | 1170.4 | 1162.8 | 1047.6 | 864.8 | 711.1 | 1028.9 | 898.6 | 854.4 | 936.4 | 739.8 | 749.5 | 0.8 | | | | | | | Porin_3 x1Porin_3 x1Porin_3 x1 | NA | NA | NA | NA | NA |
