## Supplementary material for "Genome editing excisase origins illuminated by somatic genome of *Blepharisma*": Table S6

***B. stoltei* ATCC 30299 MAC gene expression** (only genes with PFAM annotations; domain multiplicitly indicated by "x" and a number; blue = 0-1 RPKM; cyan = 1-10 RPKM; 10-100 RPKM = yellow; 100-1000 RPKM = orange; 1000-10000 RPKM = red)

| | ENA accession | Gene ID | Length (CDS in bp) | Starved | Gamone-treated | 0h | 2h | 6h | 14h | 18h | 22h | 26h | 30h | 38h | 26h/average(starved +gamone+0h) | Transposase | DNA repair | Chromatin | Transcription | Small RNAs | Translation | PFAM Domains | eggNOG gene name | eggNOG description | Pannzer gene name | Pannzer description | Pannzer PPV |
|---|---|---|---|---|---|---|---|---|---|---|---|---|---|---|---|---|---|---|---|---|---|---|---|---|---|---|---|
| 1 | BSTOLATCC_MAC22820 | Contig_6.g818 | 3780 | 0.3 | 0.5 | 0.2 | 1.4 | 51.7 | 101.1 | 105.4 | 209.8 | 278.1 | 118.7 | 65.5 | 824.9 | | ⊙ | | | | | STAG x1 | SA | STAG domain | NA | SCD domain-containing protein | 0.56 |
| 2 | BSTOLATCC_MAC12076 | Contig_36.g894 | 927 | 0.1 | 0.2 | 0.1 | 0.2 | 1.7 | 6.4 | 5.7 | 17.2 | 50.9 | 89.7 | 64.2 | 514.0 | | | | | | | CENP-B_N x1 DUF2969 x1 | NA | NA | NA | NA | NA |
| 3 | BSTOLATCC_MAC15299 | Contig_43.g546 | 1335 | 0.0 | 0.1 | 0.1 | 0.5 | 22.9 | 25.7 | 24.7 | 53.7 | 48.5 | 35.7 | 23.4 | 507.9 | | ⊙ | | | | | BRCT_2 x1 | NA | NA | NA | NA | NA |
| 4 | BSTOLATCC_MAC7513 | Contig_25.g976 | 348 | 0.0 | 0.0 | 0.4 | 37.4 | 77.2 | 34.8 | 29.4 | 59.7 | 68.3 | 40.6 | 26.3 | 506.7 | | | | | | ⊙ | HSP90 x1 | HSP90 | Heat shock protein | NA | Heat shock protein 90 | 0.39 |
| 5 | BSTOLATCC_MAC7514 | Contig_25.g977 | 1638 | 0.0 | 0.0 | 0.5 | 48.0 | 101.3 | 44.7 | 36.4 | 71.8 | 84.9 | 54.3 | 32.2 | 478.6 | | | | | | ⊙ | HSP90 x1 HATPase_c x1 HATPa | NA | heat shock protein | NA | Heat shock protein 90 | 0.59 |
| 6 | BSTOLATCC_MAC12987 | Contig_39.g182 | 3876 | 0.1 | 0.3 | 0.2 | 5.9 | 23.3 | 32.3 | 23.6 | 68.7 | 74.7 | 24.4 | 18.0 | 387.5 | | | | ⊙ | | | DUF3591 x1 Bromodomain x1 | Taf1 | binding. It is involved in the biol | NA | NA | NA |
| 7 | BSTOLATCC_MAC5291 | Contig_20.g338 | 543 | 1.7 | 5.4 | 3.1 | 102.8 | 2131.6 | 1550.9 | 2462.4 | 1970.2 | 1161.6 | 1535.0 | 1168.4 | 339.5 | | | | | | ⊙ | IF4E x1 | NA | translation initiation factor activit | NA | Eukaryotic translation initiation f | 0.22 |
| 8 | BSTOLATCC_MAC10792 | Contig_33.g1217 | 669 | 0.1 | 0.4 | 0.1 | 2.3 | 188.0 | 152.0 | 220.0 | 156.4 | 69.5 | 79.9 | 70.7 | 338.0 | | | | | | | 14-3-3 x1 | NA | 14-3-3 homologues | NA | NA | NA |
| 9 | BSTOLATCC_MAC24980 | Contig_7.g1394 | 396 | 0.1 | 0.5 | 0.1 | 1.2 | 72.4 | 57.5 | 99.4 | 107.9 | 70.8 | 97.4 | 79.5 | 297.5 | | | | | | | zf-C2H2 x2 zf-C2H2_4 x2 zf-H2C | NA | NA | NA | NA | NA |
| 10 | BSTOLATCC_MAC2188 | Contig_13.g876 | 1590 | 0.1 | 0.1 | 0.1 | 0.3 | 3.7 | 7.9 | 10.7 | 19.1 | 34.3 | 47.2 | 37.2 | 290.3 | ⊙ | | | | | | DDE_Tnp_1_7 x1 | NA | Transposase IS4 | NA | NA | NA |
| 11 | BSTOLATCC_MAC17155 | Contig_48.g755 | 1896 | 7.1 | 12.3 | 9.7 | 60.2 | 3458.6 | 4149.6 | 2813.3 | 2326.9 | 2799.4 | 2446.1 | 1305.3 | 288.1 | | | | | | | DEAD x1 zf-CCHC x4 Helicase_C | DBP2 | Belongs to the DEAD box helica | DDX43 | RNA helicase | 0.41 |
| 12 | BSTOLATCC_MAC5406 | Contig_20.g453 | 2340 | 4.5 | 9.5 | 10.1 | 150.6 | 6059.2 | 4134.1 | 3321.1 | 2959.9 | 2282.3 | 1941.8 | 1102.0 | 283.3 | | | | | ⊙ | | Piwi x1 PAZ x1 ArgoL1 x1 | NA | Piwi | NA | NA | NA |
| 13 | BSTOLATCC_MAC711 | Contig_10.g883 | 717 | 0.6 | 1.4 | 0.5 | 2.1 | 51.5 | 90.8 | 97.2 | 190.8 | 235.5 | 342.6 | 203.8 | 280.7 | | | | ⊙ | | | CID x1 | NA | NA | NA | CID domain-containing protein | 0.56 |
| 14 | BSTOLATCC_MAC4276 | Contig_19.g1573 | 879 | 0.5 | 0.6 | 1.0 | 13.6 | 198.9 | 163.8 | 191.7 | 266.7 | 200.5 | 255.2 | 175.1 | 279.5 | | | | | | | zf-B_box x1 | NA | NA | NA | NA | NA |
| 15 | BSTOLATCC_MAC23590 | Contig_60.g672 | 3246 | 0.5 | 0.7 | 0.3 | 9.4 | 45.7 | 59.4 | 40.9 | 128.3 | 146.3 | 47.3 | 35.1 | 269.7 | | | ⊙ | | | | JmjC x1 | NA | A domain family that is part of th | NA | JmjC domain-containing protein | 0.19 |
| 16 | BSTOLATCC_MAC6526 | Contig_22.g1567 | 1923 | 0.0 | 0.0 | 0.0 | 0.2 | 0.0 | 0.6 | 1.8 | 0.1 | 1.8 | 0.1 | 0.1 | 258.8 | | | | | | | UCH x1 | NA | ubiquitin carboxyl-terminal hydro | NA | NA | NA |
| 17 | BSTOLATCC_MAC6719 | Contig_23.g186 | 633 | 0.6 | 0.5 | 0.4 | 1.9 | 51.8 | 80.2 | 56.0 | 165.0 | 125.6 | 183.5 | 93.9 | 254.7 | | | | | | | CENP-B_N x1 ubiquitin x1 HTH_ | NA | NA | NA | NA | NA |
| 18 | BSTOLATCC_MAC11436 | Contig_34.g257 | 525 | 1.0 | 1.9 | 0.7 | 1.5 | 67.0 | 116.3 | 139.2 | 262.7 | 279.8 | 374.8 | 321.0 | 235.3 | | | ⊙ | | | | Chromo x1 | NA | Chromo (CHRromatin Organisati | NA | NA | NA |
| 19 | BSTOLATCC_MAC22509 | Contig_59.g228 | 1614 | 0.0 | 0.0 | 0.0 | 0.1 | 1.0 | 2.7 | 2.2 | 5.3 | 6.5 | 8.5 | 5.9 | 230.3 | | | | | | | Kelch_6 x1 | NA | NA | NA | NA | NA |
| 20 | BSTOLATCC_MAC12942 | Contig_39.g137 | 393 | 0.3 | 0.1 | 0.0 | 0.2 | 35.0 | 27.6 | 40.6 | 37.8 | 33.3 | 50.0 | 41.4 | 230.1 | | | | | | | zf-C2H2_4 x2 zf-C2H2 x2 zf-H2C | NA | NA | NA | NA | NA |
| 21 | BSTOLATCC_MAC12676 | Contig_38.g1488 | 258 | 0.5 | 1.8 | 0.9 | 6.6 | 317.8 | 232.6 | 546.2 | 439.4 | 241.7 | 273.4 | 232.3 | 223.0 | | | | ⊙ | | ⊙ | LSM14 x1 SM-ATX x1 | NA | NA | NA | NA | NA |
| 22 | BSTOLATCC_MAC5920 | Contig_21.g964 | 540 | 0.0 | 0.0 | 0.2 | 0.8 | 10.3 | 6.2 | 5.1 | 9.6 | 12.6 | 8.2 | 3.7 | 217.7 | | | | | | ⊙ | HSP70 x1 | HSP70 | heat shock protein 70 | NA | Heat shock 70 kDa protein (Frag | 0.64 |
| 23 | BSTOLATCC_MAC5919 | Contig_21.g963 | 765 | 0.0 | 0.0 | 0.1 | 0.7 | 8.0 | 5.4 | 4.0 | 7.3 | 10.6 | 5.5 | 3.1 | 216.5 | | | | | | ⊙ | HSP70 x1 MreB_Mbl x1 | NA | Belongs to the heat shock prote | HSP70 | Heat shock protein 70 (Fragmen | 0.46 |
| 24 | BSTOLATCC_MAC14490 | Contig_41.g1404 | 1590 | 1.8 | 1.4 | 1.8 | 4.6 | 40.7 | 117.4 | 139.0 | 257.0 | 342.5 | 267.0 | 177.2 | 205.5 | ⊙ | | | | | | DDE_3 x1 | NA | DDE superfamily endonuclease | NA | NA | NA |
| 25 | BSTOLATCC_MAC23800 | Contig_60.g882 | 2058 | 0.6 | 0.5 | 0.6 | 1.7 | 39.8 | 49.7 | 54.1 | 107.0 | 111.4 | 51.2 | 30.2 | 201.5 | | | | | | | DEAD x1 Helicase_C x1 ResIII x | DDX25 | RNA helicase activity | NA | NA | NA |
| 26 | BSTOLATCC_MAC18054 | Contig_5.g529 | 1812 | 0.3 | 0.1 | 0.3 | 0.4 | 5.3 | 15.3 | 16.6 | 30.4 | 40.5 | 33.5 | 25.6 | 195.6 | ⊙ | | | | | | MULE x1 Transposase_mut x1 | NA | Protein FAR1-RELATED SEQUE | NA | NA | NA |
| 27 | BSTOLATCC_MAC1548 | Contig_12.g245 | 1554 | 0.4 | 0.9 | 2.4 | 1.6 | 33.3 | 93.7 | 112.5 | 153.6 | 232.4 | 254.3 | 168.5 | 191.1 | | ⊙ | | | | | Rad21_Rec8_N x1 | RAD21 | positive regulation of sister chro | NA | NA | NA |
| 28 | BSTOLATCC_MAC13455 | Contig_4.g167 | 1164 | 0.1 | 0.2 | 0.0 | 0.1 | 8.0 | 14.1 | 18.9 | 22.6 | 19.3 | 21.7 | 12.7 | 190.2 | | | | | | | NTP_transf_2 x1 PAP_assoc x1 | NA | RNA uridylyltransferase activity | NA | Non-canonical poly(A) RNA poly | 0.12 |
| 29 | BSTOLATCC_MAC2214 | Contig_13.g902 | 891 | 0.2 | 0.1 | 0.4 | 0.3 | 13.1 | 19.2 | 23.2 | 35.5 | 40.2 | 49.6 | 24.1 | 187.0 | | | | | | | zf-RING_UBOX x1 zf-RING_5 x1 | NA | regulation of erythrocyte enucle | NA | NA | NA |
| 30 | BSTOLATCC_MAC5044 | Contig_20.g91 | 2136 | 2.2 | 2.3 | 2.9 | 4.8 | 80.8 | 193.3 | 212.6 | 370.6 | 431.5 | 397.0 | 250.7 | 174.1 | | | ⊙ | | | | JmjC x1 | NA | A domain family that is part of th | NA | NA | NA |
| 31 | BSTOLATCC_MAC6514 | Contig_22.g1555 | 1497 | 0.0 | 0.0 | 0.0 | 0.1 | 0.3 | 0.2 | 1.1 | 0.4 | 0.7 | 0.0 | 0.4 | 167.7 | | | | | | | UCH x1 | NA | ubiquitin carboxyl-terminal hydro | NA | NA | NA |
| 32 | BSTOLATCC_MAC17684 | Contig_49.g1283 | 2916 | 0.3 | 0.7 | 0.2 | 17.5 | 48.4 | 47.8 | 31.6 | 69.7 | 68.5 | 28.6 | 19.9 | 167.5 | | | ⊙ | | | | SNF2_N x1 SLIDE x1 Helicase_ | NA | helicase superfamily c-terminal | NA | Transcription activator snf2l1 | 0.73 |
| 33 | BSTOLATCC_MAC16927 | Contig_48.g527 | 960 | 1.0 | 2.9 | 1.2 | 11.7 | 86.5 | 151.9 | 155.9 | 297.4 | 282.5 | 227.8 | 153.1 | 166.9 | | | ⊙ | | | | Chromo x1 RNA_pol_Rpb2_6 x1 | NA | Chromatin organization modifier | NA | NA | NA |
| 34 | BSTOLATCC_MAC16237 | Contig_45.g1479 | 561 | 1.8 | 1.2 | 1.3 | 2.3 | 176.8 | 204.1 | 283.6 | 321.3 | 234.9 | 287.4 | 180.3 | 164.4 | | | | | | | zf-RanBP x2 | NA | Zinc finger domain | NA | NA | NA |
| 35 | BSTOLATCC_MAC16168 | Contig_45.g1410 | 1944 | 0.3 | 0.2 | 1.2 | 2.6 | 52.0 | 39.3 | 36.6 | 62.0 | 88.5 | 62.7 | 31.9 | 157.8 | | | | | | ⊙ | HSP70 x1 MreB_Mbl x1 | NA | heat shock protein 70 | NA | Cytosol-type hsp70 | 0.58 |
| 36 | BSTOLATCC_MAC15955 | Contig_45.g1197 | 456 | 1.3 | 0.8 | 1.2 | 18.5 | 184.8 | 129.7 | 182.8 | 216.3 | 163.8 | 187.7 | 97.6 | 147.6 | | | | | | | TCR x2 | NA | Tesmin/TSO1-like CXC domain | NA | NA | NA |
| 37 | BSTOLATCC_MAC18055 | Contig_5.g530 | 1812 | 0.3 | 0.4 | 0.4 | 1.3 | 4.7 | 13.3 | 16.3 | 30.1 | 54.6 | 64.9 | 43.0 | 145.2 | ⊙ | | | | | | MULE x1 | NA | FAR1 DNA-binding domain | NA | NA | NA |
| 38 | BSTOLATCC_MAC15249 | Contig_43.g496 | 1902 | 0.5 | 0.3 | 0.2 | 1.0 | 8.6 | 18.8 | 20.4 | 44.5 | 46.0 | 36.7 | 26.3 | 142.2 | ⊙ | | | | | | MULE x1 | NA | MULE transposase domain | NA | NA | NA |
| 39 | BSTOLATCC_MAC17007 | Contig_48.g607 | 1218 | 0.1 | 0.1 | 0.4 | 1.4 | 3.2 | 2.9 | 3.2 | 7.3 | 28.9 | 23.3 | 12.0 | 141.3 | | | | | | | Bromodomain x1 BET x1 | BRDT | bromo domain | NA | NA | NA |
| 40 | BSTOLATCC_MAC12545 | Contig_38.g1357 | 1542 | 0.3 | 0.2 | 0.1 | 2.6 | 11.3 | 12.3 | 12.3 | 25.5 | 26.8 | 14.6 | 9.2 | 140.3 | | ⊙ | | | | | BRCT_2 x1 | NA | NA | NA | NA | NA |
| 41 | BSTOLATCC_MAC1138 | Contig_11.g1307 | 2256 | 1.6 | 0.8 | 1.1 | 4.3 | 43.2 | 83.3 | 80.7 | 142.1 | 158.8 | 123.8 | 75.5 | 136.5 | | | | | ⊙ | | Ribonuclease_3 x2 Ribonucleas_ | dcl1 | Mortierella verticillata NRRL 633 | NA | NA | NA |
| 42 | BSTOLATCC_MAC10037 | Contig_30.g466 | 2151 | 1.2 | 1.2 | 1.0 | 7.3 | 66.6 | 81.7 | 65.9 | 152.1 | 149.7 | 94.6 | 66.4 | 133.3 | | | | ⊙ | | | NIF x1 | fcp1 | catalytic domain of ctd-like phos | NA | NA | NA |
| 43 | BSTOLATCC_MAC9080 | Contig_28.g1168 | 1581 | 0.6 | 0.5 | 0.9 | 2.4 | 16.1 | 33.9 | 31.2 | 65.7 | 86.8 | 91.4 | 60.5 | 130.0 | | | | | | | PB1 x1 | NA | NA | NA | NA | NA |
| 44 | BSTOLATCC_MAC1071 | Contig_11.g1240 | 1992 | 2.0 | 1.7 | 1.3 | 8.2 | 91.8 | 113.6 | 108.7 | 219.2 | 209.8 | 146.6 | 111.5 | 128.8 | | | | | | | Dynamin_M x1 Dynamin_N x1 G | NA | Belongs to the TRAFAC class d | NA | Dynamin-related protein 3A | 0.76 |
| 45 | BSTOLATCC_MAC25739 | Contig_9.g358 | 1014 | 1.2 | 1.0 | 0.7 | 2.6 | 44.6 | 71.5 | 76.7 | 150.0 | 120.8 | 130.3 | 80.3 | 126.5 | | | | | | | zf-B_box x1 zf-RING_5 x1 zf-RIN | NA | zinc ion binding | NA | NA | NA |
| 46 | BSTOLATCC_MAC7356 | Contig_25.g819 | 2061 | 0.4 | 0.3 | 0.6 | 6.7 | 56.7 | 49.9 | 36.9 | 56.0 | 51.7 | 36.6 | 24.6 | 124.4 | | | | | | | FH2 x1 | GRID2IP | Glutamate receptor, ionotropic, | NA | NA | NA |
| 47 | BSTOLATCC_MAC1470 | Contig_12.g167 | 783 | 1.4 | 1.3 | 1.0 | 1.2 | 29.0 | 65.8 | 97.1 | 126.6 | 142.6 | 218.1 | 186.1 | 116.0 | | ⊙ | | | | | PCNA_N x1 PCNA_C x1 Rad1 x | NA | This protein is an auxiliary prote | NA | Proliferating cell nuclear antigen | 0.42 |
| 48 | BSTOLATCC_MAC11469 | Contig_34.g290 | 651 | 0.8 | 1.4 | 2.5 | 3.9 | 73.4 | 103.7 | 138.6 | 173.1 | 175.8 | 278.8 | 208.9 | 114.4 | | | | ⊙ | | | TBP x2 | SPT15 | Transcription factor TFIID (or TA | NA | Tata-binding general transcriptic | 0.56 |
| 49 | BSTOLATCC_MAC20942 | Contig_57.g152 | 435 | 2.7 | 10.7 | 3.1 | 9.0 | 579.9 | 497.9 | 943.6 | 834.7 | 622.9 | 982.3 | 1137.5 | 113.2 | | | | | | | zf-H2C2_2 x1 | NA | NA | NA | NA | NA |
| 50 | BSTOLATCC_MAC13720 | Contig_40.g636 | 4278 | 1.2 | 2.6 | 1.1 | 37.1 | 75.0 | 83.3 | 67.8 | 144.7 | 182.9 | 83.7 | 59.0 | 111.7 | | | | | | | Nipped-B_C x1 Cnd1 x1 PHD x1 | NIPBL | Sister chromatid cohesion C-ter | NA | NA | NA |
| 51 | BSTOLATCC_MAC23155 | Contig_6.g1153 | 1380 | 0.2 | 0.7 | 0.6 | 0.3 | 6.1 | 14.0 | 20.5 | 27.1 | 59.9 | 115.1 | 79.7 | 110.3 | | ⊙ | | | | | zf-GRF x2 Exo_endo_phos x1 | APEX2 | double-stranded DNA 3'-5' exod | NA | DNA-(apurinic or apyrimidinic site | 0.12 |
| 52 | BSTOLATCC_MAC1521 | Contig_12.g218 | 567 | 1.3 | 6.5 | 2.8 | 5.7 | 175.3 | 170.8 | 196.2 | 315.0 | 387.9 | 516.2 | 444.0 | 108.8 | | | | | | | SAM_1 x1 DUF1805 x1 | NA | NA | NA | NA | NA |
| 53 | BSTOLATCC_MAC25840 | Contig_9.g459 | 402 | 1.6 | 5.1 | 7.6 | 4.9 | 233.0 | 328.6 | 626.3 | 576.4 | 517.1 | 851.2 | 682.6 | 108.8 | | | ⊙ | | | | Histone x1 Histone_H2A_C x1 C | NA | Histone H2AX | NA | NA | NA |
| 54 | BSTOLATCC_MAC16265 | Contig_45.g1507 | 2055 | 0.0 | 0.0 | 0.0 | 1.0 | 2.7 | 4.1 | 1.7 | 2.5 | 1.7 | 1.3 | 0.9 | 104.5 | | | | | | | AAA-ATPase_like x1 | NA | Psort location Cytoplasmic, scor | NA | NA | NA |
| 55 | BSTOLATCC_MAC23523 | Contig_60.g605 | 1536 | 1.7 | 1.3 | 1.6 | 26.6 | 335.8 | 220.4 | 195.8 | 240.1 | 162.0 | 99.3 | 60.6 | 104.2 | | | | ⊙ | | | RRM_1 x4 PABP x1 | NA | Binds the poly(A) tail of mRNA | NA | NA | NA |
| 56 | BSTOLATCC_MAC21072 | Contig_57.g282 | 1215 | 0.3 | 0.2 | 0.2 | 0.9 | 3.9 | 6.6 | 7.3 | 14.4 | 23.5 | 25.5 | 16.7 | 104.0 | | | | | | | ERCC4 x1 | mus81 | It is involved in the biological pr | NA | NA | NA |
| 57 | BSTOLATCC_MAC20619 | Contig_56.g1053 | 3813 | 0.6 | 1.8 | 0.6 | 41.0 | 56.3 | 42.8 | 35.6 | 81.9 | 104.8 | 40.3 | 25.3 | 103.3 | | | | | | | DNA_topoisoIV x1 TOPRIM_C x | NA | Control of topological states of | NA | DNA topoisomerase 2 | 0.63 |
| 58 | BSTOLATCC_MAC20673 | Contig_56.g1107 | 1245 | 0.5 | 0.7 | 0.3 | 23.1 | 66.7 | 23.3 | 23.2 | 60.4 | 51.5 | 31.0 | 19.3 | 103.2 | | | | | | | TolA_bind_tri x1 | NA | NA | NA | NA | NA |
| 59 | BSTOLATCC_MAC4026 | Contig_19.g1323 | 828 | 0.0 | 0.0 | 0.1 | 0.2 | 1.0 | 3.4 | 3.7 | 5.0 | 3.6 | 4.2 | 4.3 | 101.9 | | | | | | | START x1 | NA | NA | NA | NA | NA |
| 60 | BSTOLATCC_MAC18052 | Contig_5.g527 | 1221 | 1.5 | 3.8 | 3.5 | 5.1 | 75.0 | 141.0 | 160.6 | 252.5 | 291.8 | 348.1 | 289.2 | 100.1 | ⊙ | | | | | | DDE_Tnp_IS1595 x1 | NA | ISXO2-like transposase domain | NA | NA | NA |
| 61 | BSTOLATCC_MAC1058 | Contig_11.g1227 | 2148 | 0.6 | 1.0 | 1.5 | 8.2 | 42.7 | 55.3 | 50.1 | 93.6 | 101.8 | 62.4 | 41.2 | 99.7 | | ⊙ | | | | | PARP x1 PARP_reg x1 WGR x1 | PARP2 | WGR domain containing protein | NA | Poly [ADP-ribose] polymerase | 0.47 |
| 62 | BSTOLATCC_MAC6181 | Contig_22.g1222 | 471 | 0.2 | 0.4 | 0.2 | 1.0 | 11.0 | 24.0 | 17.0 | 36.2 | 28.4 | 49.2 | 48.8 | 98.7 | | | | | | | RNA_GG_bind x1 | NA | NA | NA | NA | NA |
| 63 | BSTOLATCC_MAC13381 | Contig_4.g93 | 1680 | 0.5 | 0.3 | 0.9 | 15.3 | 38.1 | 32.0 | 28.7 | 64.7 | 58.1 | 32.7 | 21.2 | 98.5 | | | | | | | Kinesin x1 Microtub_bd x1 | NA | Belongs to the TRAFAC class m | KIFC1 | kinesin-like protein KIFC1 isofor | 0.59 |
| 64 | BSTOLATCC_MAC13030 | Contig_39.g225 | 1101 | 1.2 | 2.3 | 0.9 | 6.8 | 95.0 | 110.0 | 114.5 | 174.5 | 141.0 | 138.2 | 59.3 | 98.2 | | | | | | | RRM_2 x1 | AML1 | RNA binding | NA | NA | NA |
| 65 | BSTOLATCC_MAC17965 | Contig_5.g440 | 1692 | 5.2 | 6.5 | 5.5 | 15.8 | 131.3 | 231.9 | 256.1 | 415.8 | 559.4 | 476.3 | 329.9 | 97.5 | ⊙ | | | | | | DDE_3 x1 | NA | Transposase | NA | NA | NA |
| 66 | BSTOLATCC_MAC14246 | Contig_41.g1160 | 1854 | 0.3 | 0.4 | 1.3 | 4.8 | 13.0 | 23.9 | 23.8 | 47.3 | 61.1 | 57.8 | 43.6 | 94.2 | ⊙ | | | | | | DDE_1 x1 HTH_Tnp_Tc5 x1 CEN | JRKL | Jerky protein homolog-like | NA | NA | NA |
| 67 | BSTOLATCC_MAC5266 | Contig_20.g313 | 420 | 0.5 | 0.3 | 0.8 | 2.9 | 50.6 | 51.5 | 78.8 | 81.6 | 49.6 | 74.8 | 44.3 | 92.9 | | | ⊙ | | | | CBFD_NFYB_HMF x1 Histone x1 | NA | reference M ller, K., Lindauer, A | NA | NA | NA |

| | | | | | | | | | | | | | | | | | | | | | | | | | | | |
|---|---|---|---|---|---|---|---|---|---|---|---|---|---|---|---|---|---|---|---|---|---|---|---|---|---|---|---|
| 68 | BSTOLATCC_MAC3243 | Contig_18.g542 | 816 | 3.1 | 3.9 | 1.8 | 3.6 | 80.5 | 86.6 | 109.9 | 192.0 | 270.8 | 441.2 | 325.1 | 92.4 | | | | | | | Chromo x1 | NA | NA | NA | NA | NA |
| 69 | BSTOLATCC_MAC21908 | Contig_58.g1104 | 1626 | 0.2 | 0.5 | 0.3 | 0.4 | 4.1 | 9.9 | 12.1 | 20.5 | 30.0 | 44.0 | 42.2 | 91.6 | ◉ | | | | | | DDE_3 x1 | NA | DDE superfamily endonuclease | NA | NA | NA |
| 70 | BSTOLATCC_MAC15011 | Contig_42.g259 | 576 | 2.7 | 2.1 | 2.2 | 3.9 | 42.3 | 80.1 | 107.9 | 159.7 | 210.8 | 358.4 | 206.8 | 90.4 | | | | | | | BAH x1 PHD x1 PHD_2 x1 | NA | BAH domain | NA | NA | NA |
| 71 | BSTOLATCC_MAC14030 | Contig_40.g946 | 546 | 0.4 | 33.4 | 17.0 | 721.8 | 4701.6 | 2241.2 | 3048.8 | 3099.9 | 1508.7 | 1231.4 | 861.7 | 89.1 | | | ◉ | | | | HMG_box x1 HMG_box_2 x1 | NA | NA | NA | NA | NA |
| 72 | BSTOLATCC_MAC7803 | Contig_25.g1266 | 2151 | 7.1 | 8.5 | 8.2 | 77.0 | 402.3 | 335.0 | 316.1 | 601.5 | 699.6 | 448.7 | 254.6 | 88.3 | | | | ◉ | | | Spt5-NGN x1 KOW x2 Spt5_N x1 | NA | Transcription elongation factor S | NA | NA | NA |
| 73 | BSTOLATCC_MAC16643 | Contig_46.g251 | 1158 | 0.5 | 0.8 | 1.5 | 1.5 | 31.2 | 73.2 | 100.9 | 116.3 | 82.7 | 109.6 | 79.3 | 87.5 | | ◉ | | | | | DRMBL x1 Lactamase_B_2 x1 | DCLRE1I | DNA repair metallo-beta-lactama | NA | DRMBL domain-containing prote | 0.33 |
| 74 | BSTOLATCC_MAC3873 | Contig_19.g1170 | 2829 | 0.9 | 0.6 | 0.6 | 15.9 | 90.2 | 46.9 | 33.8 | 59.0 | 61.1 | 56.7 | 35.2 | 87.5 | | | | | | | Xpo1 x1 | NA | Exportin 1-like protein | NA | NA | NA |
| 75 | BSTOLATCC_MAC20542 | Contig_56.g976 | 342 | 4.1 | 12.5 | 12.5 | 34.7 | 602.2 | 487.7 | 782.3 | 865.4 | 846.2 | 1271.3 | 1381.0 | 87.3 | | | ◉ | | | | CENP-T_C x1 CENP-S x1 TAF x1 | NA | protein heterodimerization activi | NA | NA | NA |
| 76 | BSTOLATCC_MAC1496 | Contig_12.g193 | 1728 | 1.4 | 0.7 | 1.0 | 3.1 | 30.9 | 53.0 | 48.3 | 101.6 | 86.7 | 76.4 | 57.9 | 85.2 | | | | | | | POT1 x1 | NA | NA | NA | NA | NA |
| 77 | BSTOLATCC_MAC2159 | Contig_13.g847 | 3663 | 0.2 | 0.6 | 0.2 | 12.5 | 16.6 | 15.9 | 11.1 | 30.2 | 28.3 | 9.5 | 6.9 | 82.7 | | ◉ | | | | | AAA_23 x1 SMC_N x1 AAA_29 x | RAD50 | AAA domain | NA | NA | NA |
| 78 | BSTOLATCC_MAC2096 | Contig_13.g784 | 3102 | 1.3 | 1.3 | 2.5 | 15.9 | 83.5 | 74.9 | 60.6 | 136.6 | 139.5 | 74.9 | 54.3 | 82.4 | | | | | | | AAA_12 x1 AAA_11 x1 AAA_30 | NA | AAA domain | NA | NA | NA |
| 79 | BSTOLATCC_MAC18270 | Contig_50.g122 | 432 | 0.3 | 1.1 | 0.5 | 0.3 | 4.8 | 10.5 | 13.9 | 21.2 | 52.8 | 127.1 | 102.8 | 81.5 | | | ◉ | | | | HMG_box_2 x1 HMG_box x1 | NA | NA | NA | NA | NA |
| 80 | BSTOLATCC_MAC23585 | Contig_60.g667 | 1311 | 1.5 | 1.1 | 1.2 | 1.6 | 30.1 | 52.1 | 54.3 | 84.3 | 102.1 | 102.5 | 55.9 | 81.0 | | | | | | | PRiA4_ORF3 x1 | NA | NA | NA | NA | NA |
| 81 | BSTOLATCC_MAC2210 | Contig_13.g898 | 1035 | 0.3 | 0.4 | 0.0 | 0.4 | 5.2 | 11.3 | 14.2 | 23.6 | 19.8 | 20.2 | 11.7 | 80.7 | | | | | | | zf-RING_5 x1 zf-RING_2 x1 zf-RI | NA | NA | NA | NA | NA |
| 82 | BSTOLATCC_MAC6689 | Contig_23.g156 | 1770 | 7.6 | 13.2 | 12.4 | 55.7 | 1993.4 | 2038.7 | 1002.1 | 604.3 | 881.5 | 941.1 | 499.5 | 79.8 | | | | | | | DEAD x1 Helicase_C x1 zf-CCHC | DBP2 | Belongs to the DEAD box helica | NA | DEAD box RNA helicase-like pro | 0.54 |
| 83 | BSTOLATCC_MAC8561 | Contig_27.g656 | 3987 | 0.1 | 0.9 | 0.2 | 18.0 | 22.6 | 18.7 | 8.7 | 26.6 | 32.1 | 10.7 | 6.2 | 78.7 | | | | | | | Kinesin x1 Microtub_bd x1 | NA | Belongs to the TRAFAC class m | NA | NA | NA |
| 84 | BSTOLATCC_MAC13434 | Contig_4.g146 | 612 | 2.3 | 3.5 | 2.1 | 2.9 | 135.5 | 177.0 | 291.4 | 238.2 | 208.0 | 291.4 | 322.1 | 78.7 | | | | | | | zf-RING_2 x1 zf-C3HC4_2 x1 zf-R | NA | E3 ubiquitin-protein ligase RING | NA | NA | NA |
| 85 | BSTOLATCC_MAC17791 | Contig_49.g1390 | 1164 | 1.8 | 3.3 | 1.5 | 2.0 | 35.5 | 76.5 | 98.4 | 134.1 | 169.7 | 227.1 | 167.7 | 77.9 | | | | | | | Pre-SET x1 | NA | NA | NA | NA | NA |
| 86 | BSTOLATCC_MAC6422 | Contig_22.g1463 | 342 | 2.1 | 7.7 | 4.4 | 3.8 | 166.5 | 209.8 | 242.3 | 357.4 | 366.4 | 504.3 | 527.9 | 77.6 | | | ◉ | | | | CENP-T_C x1 CENP-S x1 TAF x1 | NA | protein heterodimerization activi | NA | NA | NA |
| 87 | BSTOLATCC_MAC25631 | Contig_8.g253 | 1584 | 2.4 | 1.6 | 3.3 | 27.8 | 142.7 | 91.8 | 91.4 | 158.7 | 182.8 | 156.2 | 89.5 | 76.5 | | | | | | | ThiF x1 UAE_UbL x1 UBA_e1_th | SAE2 | Ubiquitin/SUMO-activating enzyr | NA | SUMO-activating enzyme subun | 0.1 |
| 88 | BSTOLATCC_MAC15760 | Contig_44.g1003 | 576 | 7.8 | 29.4 | 16.7 | 102.1 | 822.0 | 697.4 | 840.6 | 1266.0 | 1370.2 | 1903.7 | 1632.8 | 76.3 | | | | | | | SAM_1 x1 SAM_2 x1 | NA | NA | NA | NA | NA |
| 89 | BSTOLATCC_MAC9851 | Contig_30.g279 | 570 | 1.9 | 4.0 | 5.1 | 9.4 | 195.4 | 227.4 | 379.4 | 360.8 | 280.1 | 512.2 | 409.2 | 76.1 | | | | | | | zf-C3HC4 x1 zf-RING_2 x1 | NA | zinc finger CCCH domain-contai | NA | NA | NA |
| 90 | BSTOLATCC_MAC22628 | Contig_6.g626 | 1203 | 0.8 | 0.4 | 0.3 | 1.2 | 4.5 | 8.4 | 8.3 | 17.2 | 39.4 | 64.1 | 56.9 | 76.0 | | | | | | | HUN x1 | NA | NA | NA | NA | NA |
| 91 | BSTOLATCC_MAC16275 | Contig_45.g1517 | 3216 | 0.4 | 0.5 | 0.1 | 14.4 | 23.1 | 15.6 | 12.3 | 28.0 | 25.2 | 15.5 | 9.9 | 75.9 | | | | | | | AAA-ATPase_like x1 | NA | An automated process has iden | NA | NA | NA |
| 92 | BSTOLATCC_MAC10232 | Contig_30.g661 | 939 | 5.1 | 5.0 | 4.2 | 5.7 | 56.9 | 109.8 | 123.3 | 242.5 | 362.2 | 415.0 | 273.5 | 75.8 | | | | ◉ | | | Med26 x1 | NA | Transcription elongation factor ( | NA | TFIIS N-terminal domain-contain | 0.62 |
| 93 | BSTOLATCC_MAC5461 | Contig_20.g508 | 372 | 0.1 | 0.1 | 0.3 | 0.8 | 10.9 | 8.3 | 16.5 | 13.2 | 11.4 | 17.9 | 16.7 | 74.6 | | | | | | | EF-hand_8 x1 EF-hand_6 x1 EF- | NA | NA | NA | NA | NA |
| 94 | BSTOLATCC_MAC6118 | Contig_22.g1159 | 888 | 2.2 | 1.4 | 2.5 | 3.1 | 55.6 | 95.9 | 91.8 | 160.0 | 148.2 | 158.3 | 93.7 | 73.6 | | ◉ | | | | | BRCT_2 x1 | NA | NA | NA | NA | NA |
| 95 | BSTOLATCC_MAC8474 | Contig_27.g569 | 912 | 0.9 | 2.9 | 1.4 | 40.0 | 223.2 | 127.6 | 163.6 | 169.1 | 126.4 | 179.2 | 117.1 | 73.5 | | | | | | | Pkinase x1 PK_Tyr_Ser-Thr x1 | CDC28 | cyclin-dependent protein serine/ | CDK1 | Cell division control protein 2 | 0.43 |
| 96 | BSTOLATCC_MAC20251 | Contig_55.g693 | 390 | 1.9 | 7.6 | 15.3 | 13.9 | 395.0 | 404.3 | 541.4 | 792.2 | 604.5 | 592.2 | 398.0 | 73.1 | | | ◉ | | | | Histone x1 CBFD_NFYB_HMF x1 | H2AFY | Histone H2A | NA | NA | NA |
| 97 | BSTOLATCC_MAC9213 | Contig_29.g58 | 3291 | 0.9 | 1.9 | 1.0 | 19.6 | 51.3 | 58.5 | 50.7 | 100.8 | 89.6 | 55.9 | 46.0 | 71.2 | | | | | | | AAA-ATPase_like x2 | NA | An automated process has iden | NA | NA | NA |
| 98 | BSTOLATCC_MAC10766 | Contig_33.g1191 | 408 | 0.0 | 1.1 | 0.6 | 0.3 | 8.7 | 18.4 | 36.6 | 34.9 | 39.8 | 85.9 | 53.1 | 71.1 | | | | | | | zf-C2H2_jaz x1 zf-Di19 x1 | NA | NA | NA | NA | NA |
| 99 | BSTOLATCC_MAC5921 | Contig_21.g965 | 387 | 0.0 | 0.1 | 0.4 | 0.8 | 7.3 | 4.9 | 2.8 | 8.2 | 10.7 | 6.7 | 2.6 | 69.4 | | | | | | ◉ | HSP70 x1 | NA | ATP binding | HSP70 | Heat shock protein 70 (Fragmen | 0.43 |
| ## | BSTOLATCC_MAC23646 | Contig_60.g728 | 945 | 0.6 | 0.7 | 0.6 | 0.4 | 14.0 | 32.8 | 40.8 | 47.3 | 45.5 | 102.2 | 100.5 | 69.1 | | ◉ | | | | | Exo_endo_phos x1 | APEX1 | double-stranded DNA 3'-5' exod | NA | DNA-(apurinic or apyrimidinic site | 0.18 |
