## Supplementary material for "Genome editing excisase origins illuminated by somatic genome of *Blepharisma*": Table S7

**Table S7.** Substitution rates between *Paramecium tetraurelia* PiggyMac-like genes and PiggyMac.

| Gene abbreviation | *P. tetraurelia* gene ID | $d_N/d_S$ | $d_N$ | $d_S$ |
|---|---|---|---|---|
| PGML2 | PTET.51.1.G0380073 | 0.0773 | 1.1082 | 14.3409 |
| PGML3a | PTET.51.1.G0010374 | 0.1245 | 1.0335 | 8.3021 |
| PGML3b | PTET.51.1.G0080308 | 0.0404 | 1.1559 | 28.6183 |
| PGML3c | PTET.51.1.G0020217 | 0.1508 | 1.0885 | 7.216 |
| PGML4a | PTET.51.1.G0340197 | 0.1161 | 0.9593 | 8.2612 |
| PGML4b | PTET.51.1.G0480099 | 0.2535 | 1.1062 | 4.3641 |
| PGML5a | PTET.51.1.G0570051 | 0.0141 | 1.1514 | 81.7442 |
| PGML5b | PTET.51.1.G0510172 | 0.0138 | 1.1642 | 84.3893 |

Reference gene: PGM - PTET.51.1.G0490162
