## Supplementary material for "Genome editing excisase origins illuminated by somatic genome of *Blepharisma*": Table S8

**Table S8.** Substitution rates between *Paramecium tetraurelia* and *Paramecium octaurelia* PiggyMac and PiggyMac-likes.

| **Gene abbreviation** | ***P. tetraurelia* gene ID** | ***P. octaurelia* gene ID** | **$d_N/d_S$** | **$d_N$** | **$d_S$** |
|---|---|---|---|---|---|
| PGM | PTET.51.1.G0490162 | POCT.K8.1.G7180000 2770580243 | 0.0234 | 0.0073 | 0.3106 |
| PGML2 | PTET.51.1.G0380073 | POCT.K8.1.G7180000 2770130227 | 0.0180 | 0.0045 | 0.2507 |
| PGML3a | PTET.51.1.G0010374 | POCT.K8.1.G7180000 2770510320 | 0.0229 | 0.0082 | 0.3600 |
| PGML3b | PTET.51.1.G0080308 | POCT.K8.1.G7180000 2770810134 | 0.0818 | 0.0245 | 0.2993 |
| PGML3c | PTET.51.1.G0020217 | POCT.K8.1.G7180000 2770610330 | 0.1052 | 0.0365 | 0.3469 |
| PGML4a | PTET.51.1.G0340197_ | POCT.K8.1.G7180000 2770180100 | 0.0425 | 0.0139 | 0.3262 |
| PGML4b | PTET.51.1.G0480099 | POCT.K8.1.G7180000 2770140101 | 0.0627 | 0.0153 | 0.2445 |
| PGML5a | PTET.51.1.G0570051 | POCT.K8.1.G7180000 2770010048 | 0.0393 | 0.0110 | 0.2800 |
| PGML5b | PTET.51.1.G0510172 | POCT.K8.1.G7180000 2769800173 | 0.0596 | 0.0123 | 0.2071 |

Observed $d_N/d_S$ values for orthologous pairs of PiggyMac and PiggyMac-like proteins from *P. tetraurelia* and *P. octaurelia*.(PGML1, 95.1% nucleotide identity, PGML3c, 89.7% nucleotide identity)
