## Supplementary material for "Genome editing excisase origins illuminated by somatic genome of *Blepharisma*": Table S9

**Table S9.** *Blepharisma* PiggyMac-like substitution rates.

| Gene ID | ENA accession | $d_N/d_S$ | $d_N$ | $d_S$ |
|---|---|---|---|---|
| Contig_3.g998 | BSTOLATCC_MAC9455 | 0.0093 | 0.7106 | 76.4367 |
| Contig_13.g879 | BSTOLATCC_MAC2191 | 0.0551 | 0.8871 | 16.0867 |
| Contig_13.g927 | BSTOLATCC_MAC2239 | 0.0261 | 0.5547 | 21.2267 |
| Contig_17.g391 | BSTOLATCC_MAC3091 | 0.0087 | 0.8394 | 96.9223 |
| Contig_17.g392 | BSTOLATCC_MAC3092 | 0.0076 | 0.8195 | 107.6866 |
| Contig_60.g827 | BSTOLATCC_MAC23745 | 0.1351 | 0.8401 | 6.2209 |
| Contig_61.g932 | BSTOLATCC_MAC23855 | 0.0836 | 0.7727 | 9.2391 |

Reference gene: Contig_49.g1063
