## Supplementary material for "Genome editing excisase origins illuminated by somatic genome of *Blepharisma*": Table S10

**Table S10.** Tree topology tests with ciliate PiggyBac homologs.

| Tree | logL | deltaL | bp-RELL | p-KH | p-SH | c-ELW | p-AU |
|---|---|---|---|---|---|---|---|
| Unconstrained | -97183.2 | 0 | 0.536 + | 0.691 + | 1 + | 0.535 + | 0.688 + |
| Monophyly of all ciliate PiggyBacs | -97536.0 | 352.77 | 0 | 0 | 0 | -2.98E-82 | 6.38E-59 - |
| Monophyly of all ciliate PiggyBacs with 4 non-ciliate interlopers | -97543.7 | 360.49 | 0 | 0 | 0 | -3.73E-101 | 7.36E-59 - |
| Monophyly of ciliate PiggyBacs without Tbp7 and with 4 non-ciliate interlopers | -97199.2 | 15.98 | 0.211 + | 0.309 + | 0.68 + | 0.212 + | 0.347 + |
| Monophyly of ciliate PiggyBacs without Tbp7 and without 4 non-ciliate interlopers | -97200.0 | 16.79 | 0.253 + | 0.324 + | 0.68 + | 0.253 + | 0.363 + |

"Non-ciliate interlopers": PBLEs PiggyBac-2 and PiggyBac-5 from *Chondrus crispus,* PiggyBac-1 from *Paracoccidoides brasiliensis* and PiggyBac-1 from *Mucor circinelloides*

deltaL : logL difference from the maximal log likelihood in the set.
bp-RELL : bootstrap proportion using RELL method (Kishino et al. 1990).
p-KH : p-value of one sided Kishino-Hasegawa test (1989).
p-SH : p-value of Shimodaira-Hasegawa test (2000).
c-ELW : Expected Likelihood Weight (Strimmer & Rambaut 2002).
p-AU : p-value of approximately unbiased (AU) test (Shimodaira, 2002).

Plus signs following numbers denote the 95% confidence sets.
Minus signs following numbers denote significant exclusion.
All tests performed 10000 resamplings using the RELL method.
